# Recovering the pre-colonial population structure of Khoe-San descendant populations

**DOI:** 10.1101/2025.10.05.680541

**Authors:** Stacy L. Edington, Dana R. Al-Hindi, Neus Font-Porterias, Alexandra Surowiec, William J. Palmer, Justin W. Myrick, Paul J. Norman, Caitlin Uren, Marlo Möller, Brenna M. Henn, Austin W. Reynolds

## Abstract

San populations from Botswana and Namibia retain exceptional linguistic, cultural and genetic diversity, but few Khoisan-speaking groups remain south of the Kalahari Desert. However, historically, far southern Africa was home to many San and Khoekhoe groups. Popular opinion often implies that such populations do not contribute to the ancestry of contemporary South Africans. Here, we characterize the genetic ancestry of self-identified South African Coloured groups and reconstruct pre– and colonial population structure from 620 newly sampled individuals. These groups retain majority Khoe-San genetic ancestry (>48%), suggesting the persistence of Khoe-San ancestry to the present day. By isolating the Khoe-San ancestry component, we show that it is intermediate between the ≠Khomani San and Nama, and distinct from Kalahari Khoe-San populations. We also find that signatures of the Indian Ocean slave trade can be traced to Indonesian islands such as Sulawesi, Java, and Flores, while the South Asian ancestry is regionally non-specific.

**Teaser:** Large-scale genomic study uncovers enduring indigenous ancestry in South African populations, exposing a complex past.

## Introduction

Present-day populations in South Africa reflect diverse origins, from the Indigenous San and Khoe peoples to migrants from varying regions throughout Africa, Europe, and Asia. The region’s ethnic mosaic results from a series of complex migrations over the last two thousand years (ky). The earliest migrations of pastoralist groups from eastern Africa into the region and later influx of agro-pastoralist groups of western African ancestry have been well documented in both the archaeological and genetics literature.(*1*, *2*) However, the impacts of more recent population movements during the colonial period have been less well studied from a genetic perspective.

The arrival of the Dutch East India Company (in Dutch: *Vereenigde Nederlandsche Geoctroyeerde Oostindische Compagnie* or *“*VOC”) in 1652 initiated South Africa’s colonial period in Table Bay, situated near modern day Cape Town.(*3*) At the time, the Company intended for the Cape to act as a refreshment post for ships traveling between Europe and Asia, as its primary colonies were situated in Sri Lanka and Jakarta – historically known as Ceylon and Batavia.(*4*) But by 1690, the resupply station had grown into a permanent Dutch colony. The VOC maintained its occupation of the Cape for nearly 200 years, its rule ending when the British seized control of the region in 1806.(*4*) During this period, the VOC is estimated to have imported upward of 63,000 to 80,000 enslaved individuals from eastern coastal Africa, Madagascar, India, Sri Lanka, and the Malay Archipelago.(*3*, *4*) While historians have reconstructed the port of origin for many enslaved individuals, significant gaps remain in our understanding of how they contributed to the genetic diversity of contemporary South Africans.

Over the next several decades, European colonists continued their expansion past Cape Town reaching the Cederberg mountains to the northwest by the 1720s, extending further north into the Richtersveld and southern Kalahari in the early 19th century, and advancing into the arid inland Karoo by the 1840s (Figure 1).(*5*) Their incursion resulted in population decline and the systematic displacement of the San and Khoe Indigenous groups.(*6*, *7*) In some cases, San and Khoe individuals (hereafter collectively referred to as Khoe-San, reflecting their shared genetic ancestry) were incorporated into colonial society, forming new genetically heterogenous communities with VOC-forced migrants and colonists, later subsumed under the term Coloured. This was a pre-apartheid term historically imposed to categorize people of mixed ancestry; that is, individuals who did not fit within the constructed White or Bantu racial categories.(*3*) The term is still in-use today and a formally recognized ethnic classification in South Africa, although we acknowledge that it may carry derogatory connotations in some contexts.

**Figure 1).**
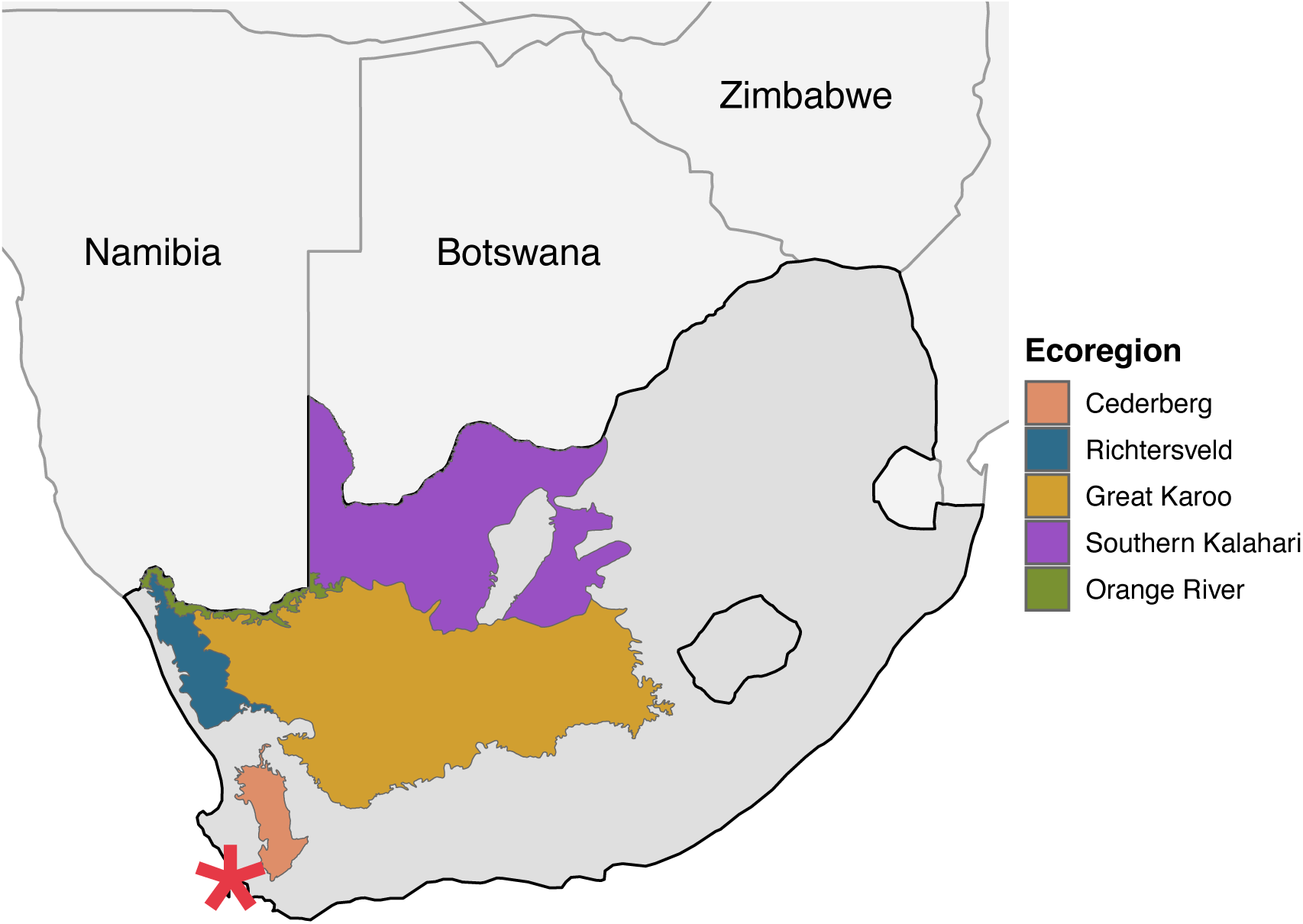
Map of South Africa highlighting the sampling regions included in this study. Colored markers indicate the geographical locations where participants were recruited: Cederberg (n=204), Richtersveld (n=206), Karoo (n=176), Southern Kalahari (n=310), and the Orange River (n=101). Total sample size across all regions was 997 individuals. The red star indicates Cape Town.

While this history is well established in academic contexts, South African Coloured (SAC) communities today have little formal exposure to this history beyond the initial founding of the VOC colony. Whether SAC outside of Cape Town have Khoe-San ancestry is often obscure unless individuals retain an oral history for their family. SAC individuals make up ∼41% of the population of the Northern and Western Cape Provinces(*8*) and are frequently described as five-way admixed,(*9*, *10*) comprised of Khoe-San, European, non-Khoe-San African, South Asian and other Southeast Asian ancestries. While the global genetic ancestry of these communities along the southern coast have been characterized (*9*, *11–13*), the temporal dynamics of admixture and the relative genetic contributions of different ancestral groups of SAC communities further from Cape Town remain largely unexplored.

We address these gaps in knowledge by analyzing 620 newly genotyped samples from self-identified SAC and Khoe-San individuals from the Western and Northern Cape Provinces, along with 22 reference populations that serve as putative sources of genome-wide ancestry from regions proximal to VOC operation sites and trade routes. These include populations from the Coromandel and Bengal coasts and across the Indonesian archipelago. Using this new robust reference panel, we investigate the genomic impacts of colonial dynamics in these regions and assess putative source populations for each of the five major distinct ancestries previously found among the SAC populations: Khoe-San, Bantu/non-Khoe-San African (NKA), South Asian, European, and East/Southeast Asian.

Specifically, we focus on the Khoe-San ancestry found in these communities. Compared to other global populations, the Khoe-San exhibit the highest levels of genetic heterogeneity, retain more ancestral alleles, and harbor autochthonous Y-chromosome and mitochondrial DNA (mtDNA) haplotypes.(*14*–*18*) Previous studies focused on Khoe-San substructure for groups in and around the Kalahari Desert, estimated to have diverged ∼20-30kya.(*19*, *20*) The Botswanan San–Tuu speaking !Xoo and Kx’a speaking Ju|’hoansi–harbor mostly northern-Kalahari Khoe-San ancestry, while the South African Tuu-speaking ≠Khomani San and Khoekhoe pastoralists Nama retain majority southern-Kalahari ancestry. Models including geographic location, language and subsistence strategies improved the predictive power of these genetic components than geography alone, suggesting that all of these factors contribute to genetic substructure within the Khoe-San.(*20*) However, the genetic affinity of Khoe-San peoples south of the Kalahari Desert, including our study region, remains largely unknown, despite this region historically being home to thousands of individuals.(*21*) We characterize pre-colonial population structure and haplotype sharing of these Khoe-San descendant individuals by isolating the Khoe-San ancestry haplotypes and assessing their genetic similarities relative to four Khoe-San reference populations: the ≠Khomani San, Nama, !Xoo, and Ju|’hoansi.

Finally, we estimate the proportion of Khoe-San ancestry that persists across more than a dozen contemporary communities, based on sampled individuals from the Cederberg, Karoo, Middle Orange River, and Upington regions and investigate how geographic and environmental factors influence gene flow between these communities. Previous efforts to characterize the Khoe-San ancestry among SAC populations were more heavily focused on the Western Cape(*11*, *22*) or were limited by small sample sizes from the Northern Cape Province (∼50 total),(*9*, *20*, *23*) potentially missing representation for the broader population.

## Results

### Genetic composition of Khoe-San descendant communities

We characterize the genetic composition of the largest set of SAC samples (n=997) derived from outside of Cape Town, including newly collected samples spanning four southern African regions (Cederberg, Karoo, Upington area, and Middle Orange River (n=620); Figure 1, Figure 2C). Newly collected southern African samples were assayed for 2.5 million markers on Illumina’s H3Africa array, then merged with a reference panel comprised of 1,165 individuals from inter– and intra-continental populations (see *Sample consolidation and PCA* Methods). Building upon previous studies, we assembled a comprehensive reference panel that aims to capture greater depth and fine-scale genetic diversity of southern African populations. The reference panel was also assembled with the goal of historically contextualizing the various migrations into southern Africa(*9*, *11*, *13*, *23*) Here, we include populations that represent six broad genetic ancestries: Niger-Congo, Khoe-San, European, South, Southeast (Indonesian), and East Asian ancestries, with >120 samples representing each ancestry (Figure 2A and 2B). Niger-Congo languages are the most commonly spoken language families spoken across sub-Saharan Africa, as a result of the Bantu-expansion. Populations included in this study and are those whose broader ancestry is labeled Niger-Congo reside western, eastern and southern Africa, but can largely trace back their genealogies to western Africa. We include populations to capture western and eastern African Bantu-expansion streams,(*24*, *25*) 5 East African populations, 130 Indonesian genomes, and over 500 individuals representing 4 Khoe and San populations to capture both northern– and southern-Kalahari genetic signals (*Supplemental Table 1*). The final merged dataset was comprised of 806,460 autosomal markers.

**Figure 2).**
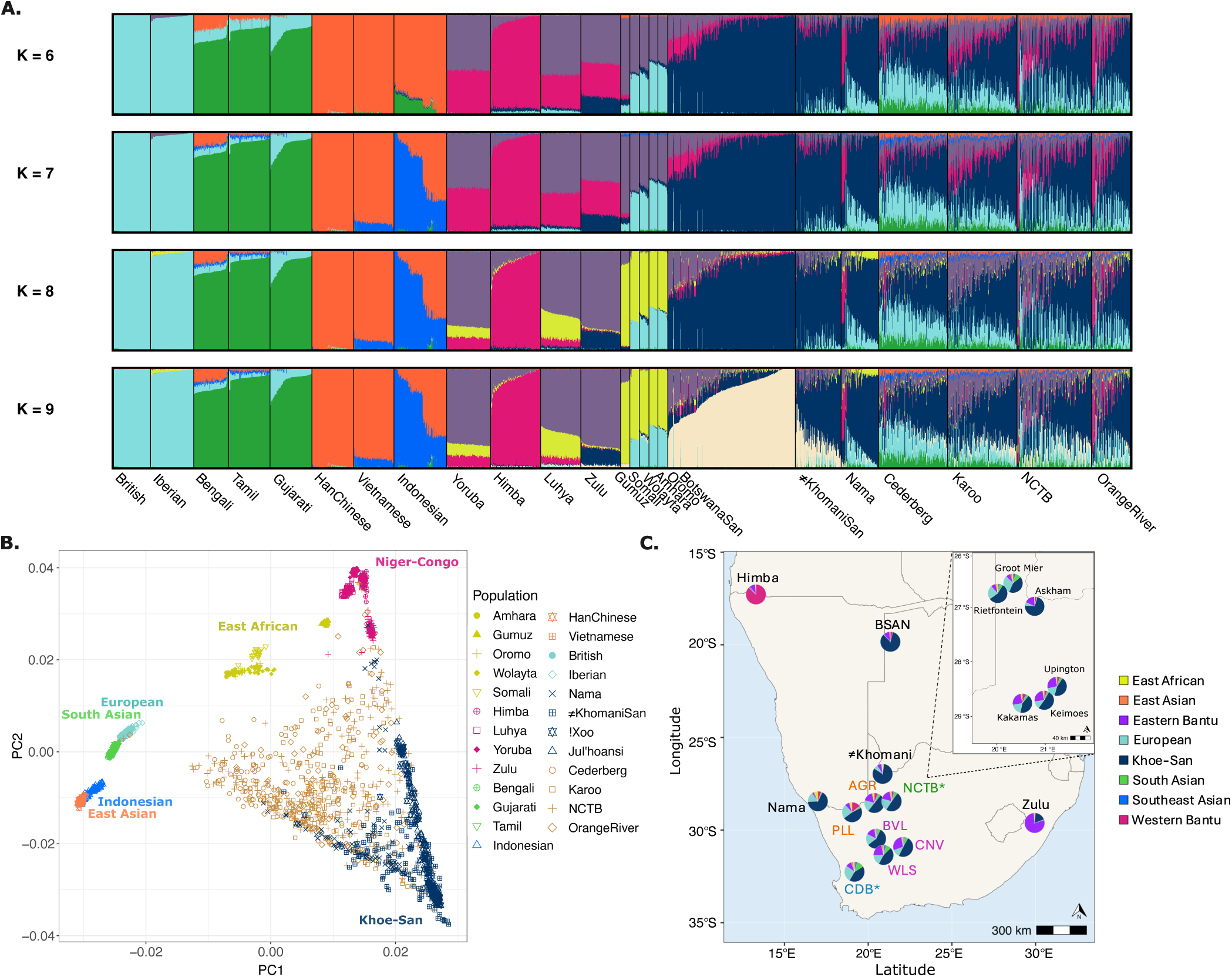
Characterizing global population structure within newly collected samples. Colors throughout the figures corresponding to broadly inferred ancestry from ADMIXTURE *k*=8. Population labels and abbreviations correspond the respective genome project (1000GP, African Genome Variation Project, and HGDP), with the exception of Indonesians (IND). In some instances, the !Xoo and Ju|’hoansi, who were inferred based on a series of analyses (*see Supplemental Text*) are merged into one population: Botswanan San (BSAN). **A)** Unsupervised ADMIXTURE run of *k*=6 through *k*=9. **B)** PCA on 806,460 congruent H3Africa array markers. Visualized are 1,165 individuals along PC1 and PC2, with shapes distinguishing populations within broader inferred ancestries (*k*=8). Study populations are colored beige, while shapes indicate sampling locality within South Africa. **C)** Approximate locations and ADMIXTURE *k*=8 results of southern African populations used in the study. Newly collected samples are segregated and colored within sampling regions, with the exception of the Cederberg (CDB) in blue and the Northern Cape Tuberculosis (NCTB) project in green. Towns sampled across the CDB are to close in proximity to segregate, while the inset map depicts sampling locations for the NCTB project. Orange River (orange) samples were collected from Pella (PLL) and Augrabies (AGR), while Karoo (purple) samples were collected across three towns: Williston (WLS), Brandvlei (BVL), and Carnarvon (CNV).

Following quality control filtering (see Methods), newly collected samples were assessed using principal component analysis (PCA). (*26*) Principal Component (PC) 1 segregates populations out-of-Africa from Khoe-San populations, while PC2 separates within-Africa populations, mainly the southern African Khoe-San groups from Niger-Congo speaking populations originating from western/central Africa (Figure 2B). The majority of Karoo (KRO), Cederberg (CDB), Upington area (NCTB), and Middle Orange River (ORG) communities are distributed across the center of PC1 and PC2. Similar distribution of reference population and sampled SAC communities has previously been observed.(*9*, *13*)

We further ran unsupervised ADMIXTURE(*27*) runs across multiple levels to infer ancestry clusters, setting *k*=4:10 (Figure 2A; *Supplemental Figure 6*), with *k*=9 and *k*=10 demonstrating the lowest cross-validation error rates (cv error=0.519, *see Supplemental Figure 7*). At *k*=5, we observe ancestries that correspond to European, South Asian, East and Southeast Asian, Khoe-San and non-Khoe-San African (NKA). Using ADMIXTURE k=5 results, we assessed these ancestry proportions within the South African sampled populations and their sampling region’s geographical distance from Cape Town. We find an increasing gradient of Khoe-San ancestry with increased distance from Cape Town, ranging from an average of 40% near Cape Town to 80% in the southern Kalahari (R^2^=0.74, p=0.013; Figure 2C and Figure 3). Conversely, South Asian and East and Southeast Asian ancestries show a negative correlation with increasing distance from Cape Town (R^2^=0.92, p=6.37×10^-4^ and R^2^=0.86, p=2.85×10^-3^, respectively; Figure 2C and *Supplemental Figure 9*), while distance from Cape Town explains a lower amount of the variability in European (R^2^=0.63, p=0.033) and NKA ancestry (R^2^=0.042, p=0.66; Figure 2C and *Supplemental Figure 9*).

**Figure 3).**
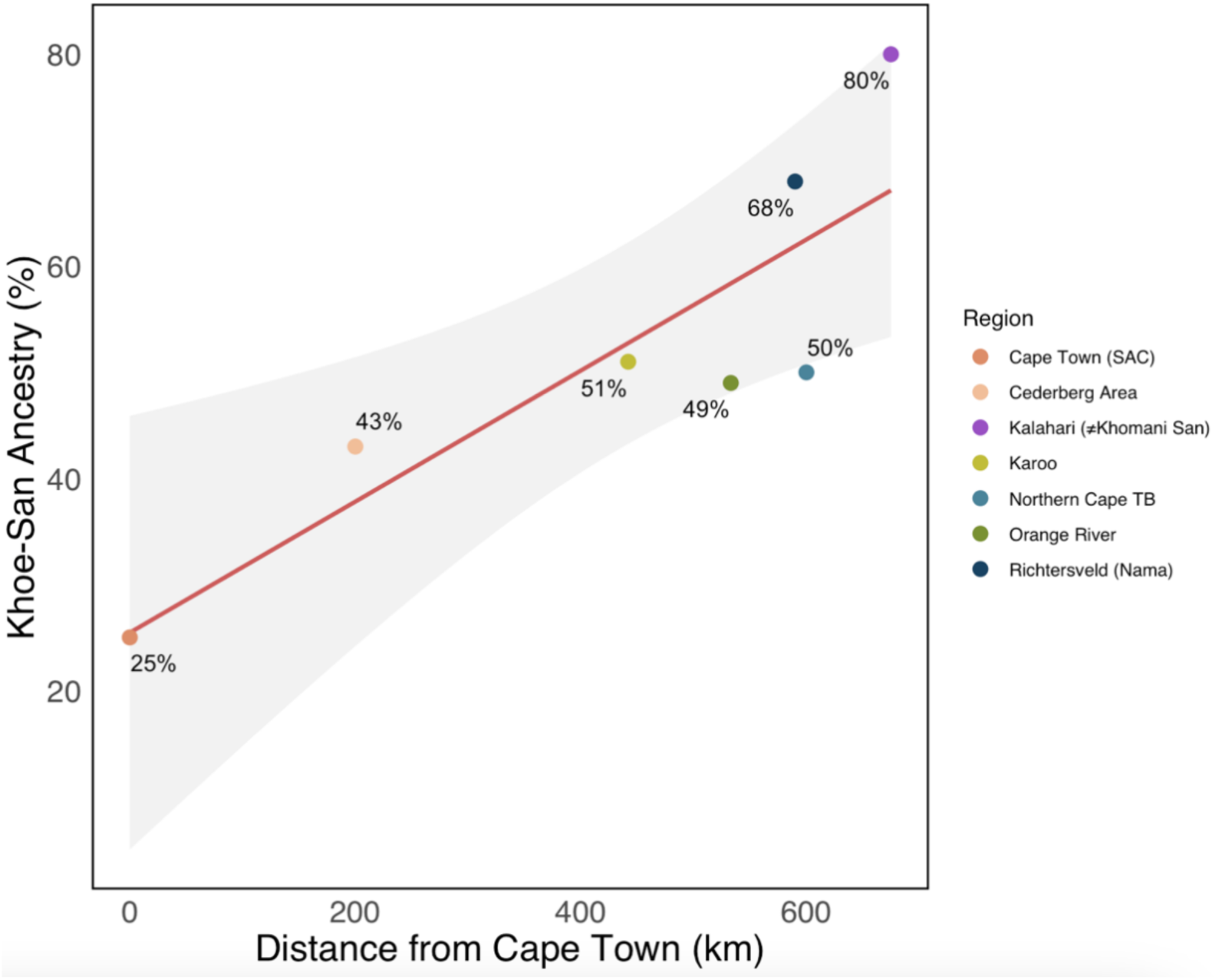
Precent of Khoe-San ancestry across southern African populations by distance from Cape Town, South Africa. R^2^ = 0.74.

The inferred NKA ancestry begins to segregate into two components at *k*=6, one of which is highest in the Himba from Namibia, likely a result of their endogamy and recent bottleneck.(*28*) At *k*=7, the southeastern Asian component distinguishes inferred ancestry in Indonesians from the Han Chinese and Vietnamese. A proportion of Indonesian ancestry is present in the sampled SAC populations, though the extent varies across groups. The CDB exhibits the highest proportion at 3.1% ± 2.1%, followed by the KRO at 2.1% ± 1.4%, while the NCTB and ORG communities show slightly lower proportions, 1.3% ± 1.4% and 1.1% ± 1.3%, respectively. Unsurprisingly, this likely has a correlation with distance from Cape Town, as previously published SAC samples from District 6, Cape Town were reported to have a higher Indonesian (∼20%) and South Asian ancestry (21%).(*13*)

Eastern African ancestry begins to differentiate from the other populations at *k*=8, shown in yellow (Figure 2A and 2C). The Gumuz show highest frequency of this component compared to other Ethiopian and Kenyan groups, with smaller proportions also present in the Yoruba, and Nama (∼6.5%). The relatively elevated East African ancestry in the Nama has been previously identified by genetic and archeological research, due to the sheep/goat/cattle pastoralist diffusion around 2kya.(*9*, *29*, *30*) However, this ancestry is a minority among the ≠Khomani San (∼1.5% ± 2.1%) and Botswanan San populations (∼1.5% ± 1.4%), with most of NKA ancestry similar to the Yoruba, Luhya and Zulu (and variably affiliated with the Himba across *k*’s). The ORG sampling sites (Pella and Augrabies) collectively show higher East African ancestry (3.4% ± 2.3%) compared to the other populations (CDB 2.4% ± 1.5%, NCTB 2.0% ± 1.9%, and KRO 1.0% ± 1.1%; see Figure 2A and 2C).

The northern– and southern-Kalahari Khoe-San ancestries segregate at *k*=9, primarily differentiating the ≠Khomani San and Nama from the two Kx’a speaking Botswanan San populations, with the latter representing a northern-Kalahari ancestry. Together, the Botswanan San populations harbor the most northern-Kalahari ancestry (76.4% ± 18.5%), relative to other Khoe-San reference populations, and minimal southern-Kalahari ancestry (10% ± 6.6%). The majority of the Khoe-San ancestry both the ≠Khomani San and the Khoekhoe-speaking Nama is southern Kalahari (60.0% ± 12.7% and 69.5% ±19.2%, respectively), while the ≠Khomani contain a notably higher proportion of northern Kalahari ancestry (25.5% ± 15.2%) compared to the Nama, who show only 2.2% ± 2.9%. Populations sampled from the CDB, KRO, ORG, and in Upington area (NCTB), share relatively consistent southern-Kalahari Khoe-San ancestry, ranging between 42.0% and 46.8%, while northern-Kalahari frequencies are uniformly lower, spanning from 2.6% in the CDB to 9.0% among NCTB participants, similarly to other sampled SAC populations from South Africa.(*9*) Interestingly, at this *k* value we find the Zulu encompass 16.7% ± 1.7% southern-Kalahari Khoe-San ancestry, compared to their low northern Khoe-San component of 2.3% ± 0.9% (Figure 2A; *Supplemental Figure 6*).

Finally, at *k*=10, a new cluster (lime green) appears in a subset of Indonesian individuals. At this resolution, the previously inferred East Asian component (orange, highest among the Han Chinese) found across the CDB, KRO, ORG, and NCTB participants *k*=7:9 now clusters with new Indonesian ancestry (*Supplemental Figure 6*). The CDB displays the highest proportions of this component 3.0% ± 1.8%, followed by 2.3% ± 1.7% in the KRO, 1.4% ± 1.8% in the NCTB, and 0.8% ± 1.3% in the ORG. The ≠Khomani San results replicate previously published findings, showing little to no Indonesian ancestry from either cluster (0.005%), while the Nama and Botswanan San show no detectable Indonesian clusters across all k’s (<0.001).(*13*)

### Identifying potential sources for African population structure

To investigate potential sources for African population structure, we performed multidimensional scaling (MDS) analysis on our sampled populations from the CDB, KRO, Upington area (NCTB) and the ORG. Ancestry specific MDS (asMDS) allows for a “zoomed in” view on specific ancestry components in admixed populations. Local ancestry inference (LAI) was preformed using GNOMIX with five broad reference groups (Khoe-San, European, South Asian, non-Khoe-San African and East and Southeast Asian).(*31*, *32*) Ancestry-specific haplotypes were then projected via ancestry-specific multidimensional scaling (asMDS) (*see Methods*; Figure 4A and 4B).(*31*) We aim to best characterize specific source populations for the five primary ancestries within the admixed study populations.

**Figure 4).**
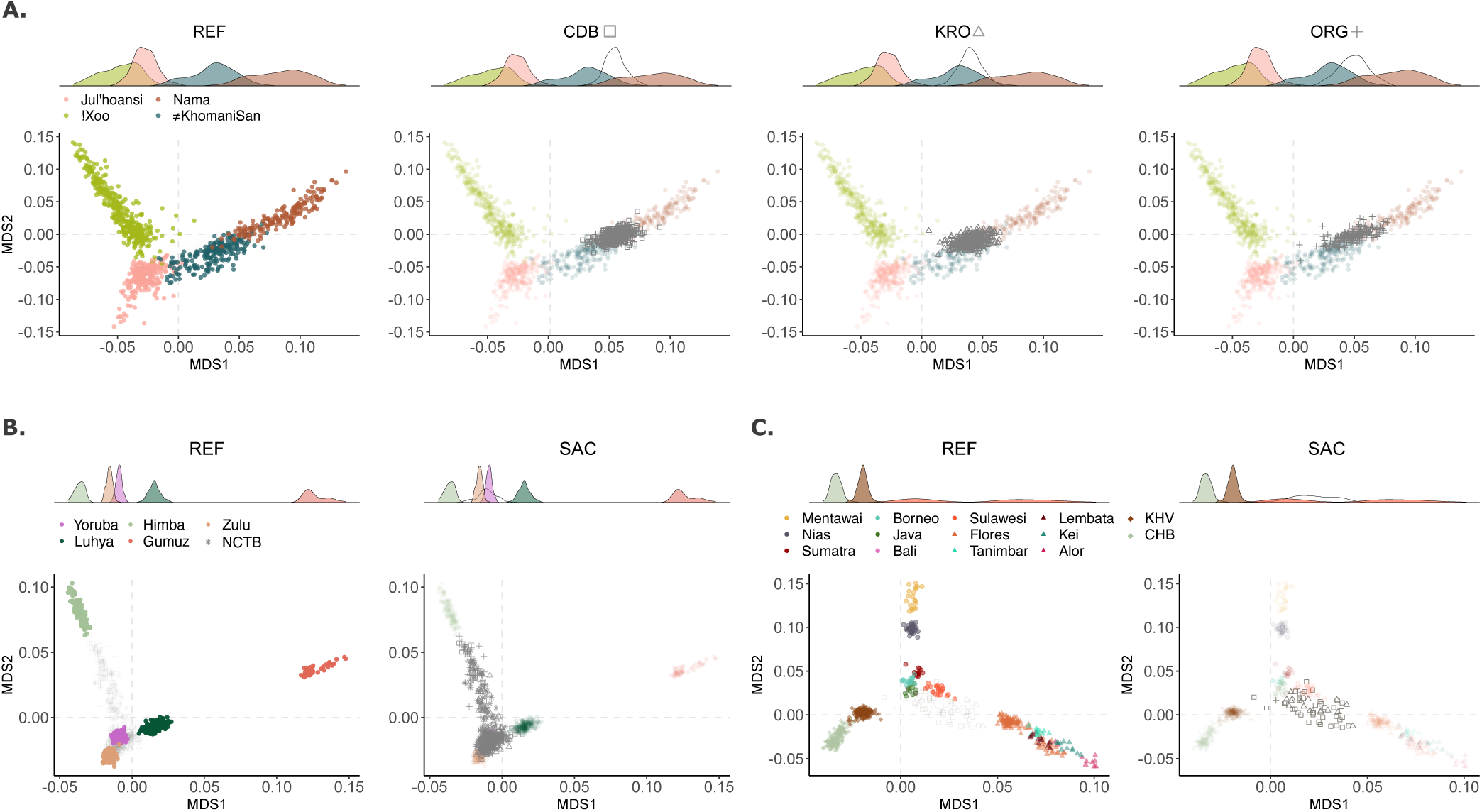
Ancestry-specific MDS analysis across three ancestries. In all panels, the left ancestry-specific MDS (asMDS) plot highlights the reference population, while the right focuses on the queried population’s projected haplotypes, mainly the Cederberg (CDB), Orange River (ORG), and Karoo (KRO). Haplotypes were filtered for a minimum 75% probability for the respective ancestry being assessed. Shapes for our sampled populations are found beside their respective asMDS. Each asMDS figure is accompanied by a density plot corresponding to asMDS1. **A)** Khoe-San specific MDS analysis with each of our sampled populations (labelled SAC) in the right three figures. **B)** Non-Khoe-San African (NKA) specific MDS analysis. The panel on the right jointly visualizes the CDB, ORG, and KRO projected haplotypes, with the addition of NKA haplotypes found among ≠Khomani San (KHM) and Nama individuals with less than 85% Khoe-San ancestry. **C)** Southeast and East Asian specific MDS analysis. Here, Indonesian samples are labeled and colored by island. Shapes geographically differentiate Asian populations into three regions: mainland (diamond), western Indonesians (circles), and eastern Indonesians (triangles).

Our Khoe-San asMDS analysis incorporates South African and Botswanan Khoe and San groups with diverse Khoisan languages. Specifically, the reference populations included are the !Xoo, ≠Khomani San, Ju|’hoansi, and the Nama (Figure 4A).(*33*, *34*) Additionally, included with our sampled populations was a control group, the Zulu, who have previously been characterized to have ∼20% Khoe-San ancestry(*35*) (Supplemental Figure 17). We replicate previously observed structure within the four Khoe-San reference populations, with the first axes largely distinguishing between the northern and southern Khoe-San ancestries (i.e. structure between the Botswanan San and the ≠Khomani San and Nama).(*19*, *36*) The refined !Xoo and Ju|’hoansi groups (originally subsumed under “BSAN”) centroids are distant from ≠Khomani San than the Nama (*see Supplemental Table 5*). The second axis separates the !Xoo and Ju|’hoansi from one another. Khoe-San inferred haplotypes from the Cederberg, Karoo, and NCTB, and Middle Orange River cluster between the overlapping tail ends of Nama and ≠Khomani San haplotype distributions along asMDS1 (Figure 4A), though all study populations’ centroids lie closer to the ≠Khomani San than the Nama (*Supplemental Table 5*).

The first dimension of our NKA asMDS segregates Yoruba and western Bantu-speaking populations (Zulu, Himba, and Luhya) from the eastern African reference populations (Gumuz) (Figure 4B). Along the second axis, we find the Himba pull away from the other broadly western African populations, likely due to a bottleneck which the population has experienced. Unexpectedly, the haplotypes from our sampled populations largely fall on top/near the Yoruba, Zulu, and Luhya, with some individual haplotypes’ exhibiting a cline towards the Himba (Figure 4B, *Supplemental Figure 19*). Notably, the Middle Orange River individuals exhibit a cline toward the Himba, though still more similar to other broadly southern African and western Bantu-speaking NKA groups (*Supplemental Figure 19*).

### Identifying potential ancestry sources for non-African population structure

Here we assess potential ancestry sources for non-African population structure with asMDS analyses on European, South Asian and East and Southeast Asian haplotypes. The European asMDS includes 5 reference populations: British from England and Scotland (GBR), Iberian populations in Spain (IBS), Toscani in Italia (TSI), Finnish in Finland (FIN), and the French. The inclusion of Dutch genomes would have been an ideal European ancestral reference; however, we were unable to obtain access of the available data for our analyses. The local ancestry prediction accuracy was the lowest for this ancestry (89.7%), with most of the model’s confusion lying on misassignment of South Asian segments as European 6.9% of the time; though the *k*=5 ADMIXTURE correlation is still high (R^2^=0.963; *see Supplemental Text and Figures 10-13*). We assess European haplotypes found among our sampled individuals and within ≠Khomani San and Nama individuals with less than 85% Khoe-San ancestry and filtered our study samples for those with at least 20% European ancestry. We find European-assigned haplotypes in our sampled populations cluster within and between the British and French populations (*Supplemental Figures 16A and 20*), though population centroids are closer to the French (Supplementary Figure 20 *and Table 7*). This pattern is concordant with historical migration from these regions into South Africa. (*37*)

The South Asian asMDS is less specific regarding the putative source populations (*Supplemental Figure 16B)*. We used 5 reference populations: the Bengali in Bangladesh (BEB), Indian Telugu in the UK (ITU), Punjabi in Lahore, Pakistan (PJL), Gujarati Indians in Houston, Texas, USA (GIH), and Sri Lankan Tamil in the UK (STU). As with the European ancestry analysis, we included haplotypes from ≠Khomani San and Nama individuals with less than 85% Khoe-San ancestry and filtered our samples for those with at least 20% South Asian ancestry. We observe a regional cline from North-West to South-East along MDS1, with the Gujarati (western India) and Punjabi (Pakistan) anchoring one end and the Bengali, Telugu, and Tamil anchoring the other. Along MDS2 the Bengali separate from the Telugu and Tamil. Haplotypes found within the Cederberg, Karoo, and Orange River participants generally pulled between the Bengali, Gujarati, and Punjabi but are generally scattered, with some clustering towards the Telugu and Tamil (*Supplemental Figure 16B*). Though we are unable to definitively state which source populations among these references best captures the South African haplotypes, there is a noticeable increase of South Asian haplotypes derived from Cederberg samples compared to our other sampled populations likely reflecting their closer proximity to Cape Town (*Supplemental Figure 16B*).

Included in the East and Southeast Asian asMDS analysis are the Han Chinese and Vietnamese from 1000Genomes alongside 130 Indonesians (Figure 4C). The East and Southeast Asian ancestry comprises a small fraction of the study population’s overall genetic composition, reaching 4.7% in the CDB and as low as 1.4% in the ORG. Given the region’s slave trade and historical record, we hypothesize Indonesians would best represent this component found among SAC populations. Our LAI model’s accuracy at assigning this ancestry is 96.3% and correlates well with reported *k*=5 ADMIXTURE proportions (R^2^=0.943) (*Supplemental Figures 10-13)*.

We refine previous research, which demonstrated SAC individuals from the Western Cape shift towards Indonesians,(*23*) by implementing an haplotype-based analysis using Indonesian genomes sampled across 12 islands and a larger, more diverse SAC cohort.(*13*, *23*) The first dimension separates eastern Indonesian islanders from mainland East Asians, while the second dimension captures previously observed regional variation within Indonesia, distinguishing western from eastern islanders.(*38*) We find Indonesians are more accurate proxies for the East and Southeast Asian components by observing clear haplotype clustering from the study populations near Indonesian samples as opposed to the Han Chinese or Vietnamese (Figure 4C). Here, we observe more specific clustering near western islands Sulawesi, Sumatra, and Java, with some trending towards Flores near regions that correspond with major ports during the Dutch East Indian Ocean trade routes.(*39*, *40*)

### Geographic predictors of population structure

To estimate how geographic and environmental factors impact human population structure, we used the machine learning tool SPRUCE (Spatial Prediction using Random Forest to Uncover Connectivity among Environments). SPRUCE uses the migration rates and demes outputs from MAPS (Migration and Population Surface estimation) as the dependent variable in the random forest regression (*see SPRUCE methods)*.(*41*) The inferred symmetric migration rate (m) is converted to dispersal distance (s) by MAPS “which can be interpreted roughly as the expected distance an individual disperses in one generation”(*41*). We estimated two time periods, approximately 56-13 generations ago (individual IBD segments 2-6cM) and less than 13 generations ago (individual IBD segments greater than 6cM).(*42*) The migration surfaces were similar for the two time periods. In Figure 5C, corresponding to ∼13 generations ago, we see there are slightly more barriers to migration in the Cederberg and relatively more migration within the Greater Karoo compared to 56-13 generations ago (Figure 5A). Random forest regression indicates that environmental variables remain similar across time periods, though some variation exists. The most important variables explaining variation in migration rates are mean temperature, slope, and minimum temperature during the coldest month (Figure 5B, 5D, *Supplemental Figure 27*).

**Figure 5).**
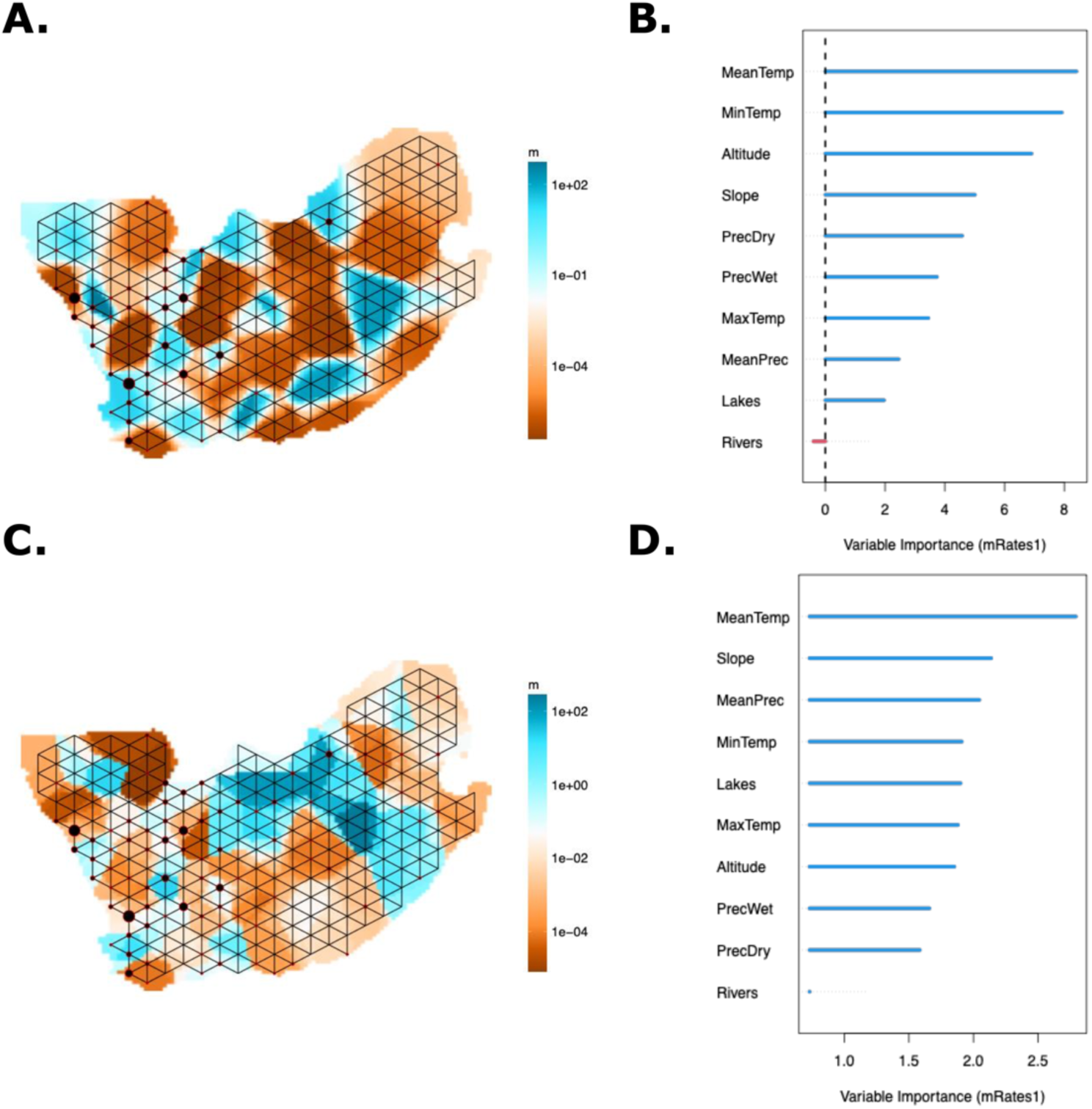
Environmental and geographical factors affect population structure. In both panels A and C, the blue color represents higher migration rates and fewer obstacles to migration; the orange color represents barriers to migration and a lower migration rate. **A)** MAPS results showing estimated migration rates for IBD segments 2-6cM representing 56-13 generations ago. **B)** SPRUCE Random Forest importance table with mean annual temperature and minimum temperature during the coldest month being the most important (Pearson Correlation R = 0.259). **C)** MAPS results showing estimated migration rates for IBD segments greater than 6cM representing less than 13 generations ago. **D)** SPRUCE Random Forest importance table with mean annual temperature and slope being the most important variables explaining the variation in estimated migration rate (Pearson Correlation R = 0.387).

To better understand the demographic connections in relation to geography, we further investigated IBD sharing within and between communities using GERMLINE2 (*see Methods)*. We see higher mean shared IBD segments within communities than between, with the Nama having the highest mean shared IBD (Figure 6; *Supplemental Figure 29*). The IBD network shows patterns of sharing between communities that are consistent with the expectation that geographically closer communities are more likely to share IBD than more distant groups. However, this relationship is relatively weak, suggesting that other social and geographic factors may play a role.

**Figure 6).**
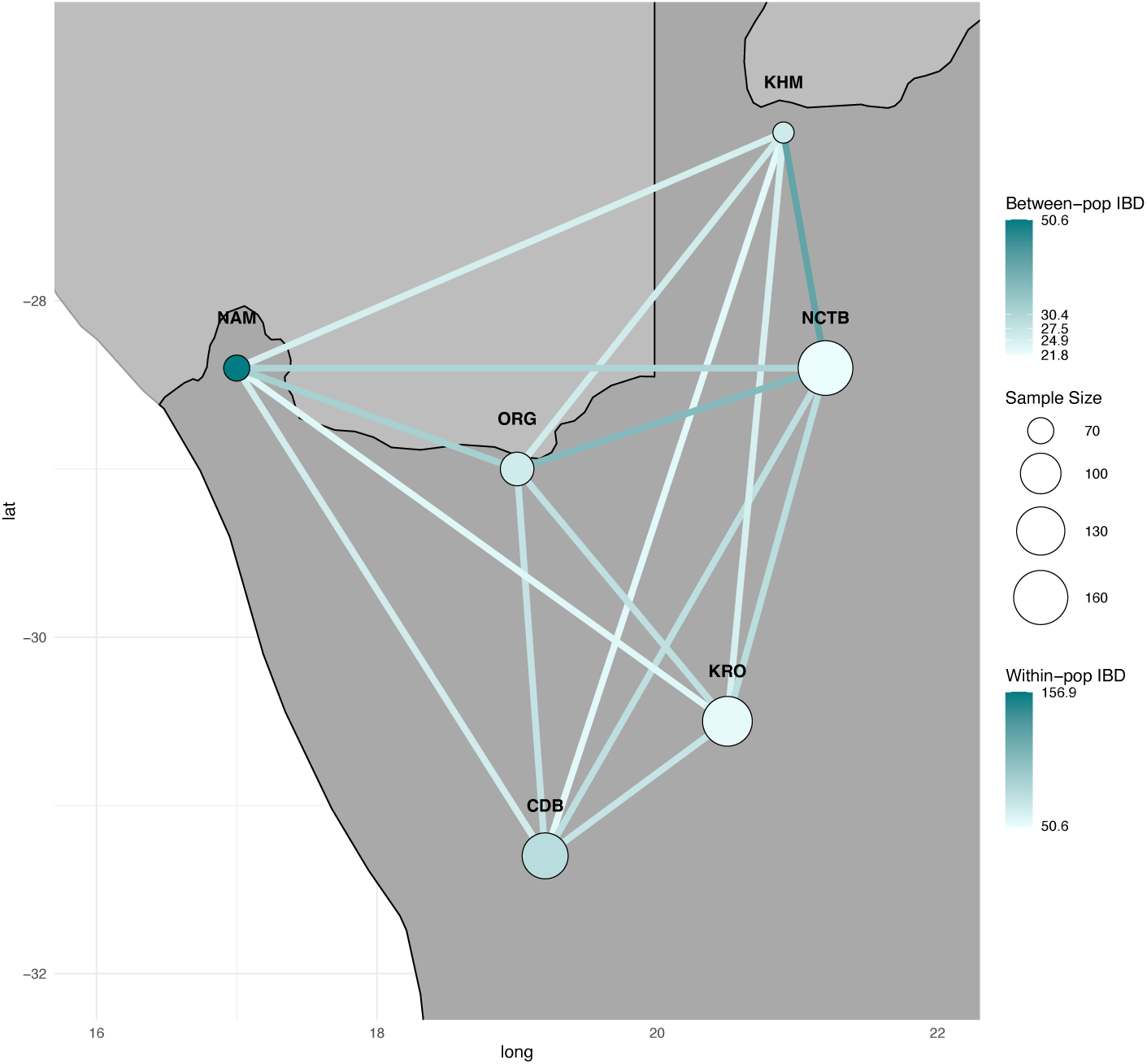
IBD sharing indicates low levels of migration across communities. The nodes are located on geographical sampling coordinates from each community and are scaled by the number of individuals sampled in that region. The nodes are colored based on within-community mean IBD sharing. The edges between groups are colored based on mean IBD sharing between communities.

## Discussion

The genetic diversity of present-day South African populations has been significantly shaped by a series of complex migrations over the past two thousand years. Previous studies have predominantly focused on disentangling Khoe-San population structure and understanding deep human evolutionary history with relatively less attention given to colonial impact and population dynamics.(*11*, *20*, *43–45*) There is limited knowledge of how colonization impacted the genetic ancestry of Indigenous populations in South Africa, largely due to the systematic displacement of these populations during and after the colonial period. Today, few individuals self-identify as members of South Africa’s Indigenous Khoe and San groups.

Previous studies have examined the complex ancestry components of SAC individuals in Cape Town and communities along the southwestern coast.(*9*, *30*) Based on a large-scale study of over 600 participants, we provide a comprehensive genetic analysis of SAC populations outside of Cape Town (n=620) and characterize genetic composition across multiple regions of South Africa (Cederberg, Karoo, Upington, Middle Orange River; Figure 1, Figure 2C). Our data demonstrates that the SAC population in the northwest of the country is shaped by multiple sources of gene flow, consistent with prior findings in Cape Town and the surrounding areas.(*11*, *20*, *22*, *46*, *47*) European colonization introduced not only European ancestry but also the forced migration of enslaved peoples through the Dutch East India (VOC) slave trade. This brought additional ancestry components into Indigenous Khoe-San populations, including gene flow from western and eastern Africa, South Asia, Malayasia, Madagascar, and Indonesia (*Supplemental Figure 6*).(*9*, *11*, *20*, *22*, *30*, *46*, *47*) Using ancestry-specific multidimensional scaling (asMDS) we were able to differentiate between putative ancestral source populations for the five ancestries commonly found within SAC communities(*46*, *47*) and identify better proxies for these sources.(*20*)

### Western African ancestry reflects both ancient Bantu expansion and colonial slave trade

Our analysis reveals distinct patterns of admixture timing and intensity across the Cederberg, Karoo, and Middle Orange River/Upington areas. NKA ancestry varies geographically, from approximately 11% in the Cederberg to about 22% in the Karoo and along the Middle Orange River (*Supplemental Table 2*, Admixture *k*=5). This distribution likely captures signatures of the Bantu expansion, with this component accounting for roughly 80% of genetic composition among southern Bantu-speaking Zulu, or it may represent NKA lineages introduced by the VOC’s slave trade.

Previous studies have found the NKA admixture into the Khoe-speaking hunter-gathers and Kxa-speaking hunter-gatherers to vary in timing, ranging from as early as 1.2k to 1.5kya.(*19*) The Richtersveld Nama show a pulse of NKA gene flow at 700 years ago. These dates suggest much of the gene flow observed in the northern Khoe-San populations dates to the initial wave of Bantu expansion across the Kalahari.(*24*, *25*) However, another wave of NKA ancestry was introduced to the Cape Colony as the VOC forcibly relocated enslaved individuals from Madagascar, Mozambique, and along the Swahili Coast.(*3*) These multiple waves of admixture are supported by tract length patterns in the Cederberg Mountains. The decay pattern of NKA tracts resembles that observed for South Asian ancestry, both suggesting colonial-era admixture events. This contrasts with tract length profiles in Karoo and Middle Orange River participants, which indicate older gene flow (*Supplemental Figure 14*).

After performing ancestry-specific MDS, the NKA component often closely resembles southeastern Bantu-speaking groups rather than southwestern Bantu-speaking populations (Figure 2). In the Karoo town of Carnarvon, this ancestry is consistent with self-reported Xhosa parents or grandparents among study participants. Conversely, the Nama and Middle Orange River participants have higher proportions of southwestern Bantu-speaking ancestry (∼5.6% and 9.5%, respectively) relative to other sampled populations (≤3.4%). This pattern positions their population centroids closer to the Himba rather than western African reference populations in NKA asMDS analysis (Figure 2C and *Supplemental Figure 19*). Ethnographic data from participants sampled along the Middle Orange River frequently report being “Nama-Dama” or having a Damara ancestor. The Damara are a historically mobile population who today reside in the semi-arid regions of Namibia and Northern Cape, South Africa.(*21*) Although the Damara are not included in our study, the closely related Himba serve as the best available proxy to reference the southwestern Bantu-expansion ancestral component.(*48*)

### Indonesian-ancestry and other Inter-Continental Migration

Initially established as a refueling station in 1652, the Cape Colony (now known as Cape Town) became the gateway for intercontinental migration into southern Africa. While many of the early colonists were Dutch, migrants from other western European regions also became part of the Cape Colony. Protestant refugees (known as the French Huguenots) arrived in the late 17^th^ century fleeing religious persecution. They lived in South Africa under Dutch rule in areas like Franschhoek and Drakenstein Valley in the mountainous region to the northeast of Cape Town.(*49*) The British gained control of the colony from the Dutch in 1806.(*50*, *51*) Reflecting the European colonial history at the Cape, our study populations retain European haplotypes intermediate to French and British populations in asMDS analysis, with population centroids positioned closer to the French (*Supplemental Figure 20*). Although French and British populations serve as reasonable proxies, a limitation of our study is the lack of access to Dutch reference genomes, which would have been most appropriate given Dutch colonial history.

The VOC’s extensive Indian Ocean trade network also facilitated the forced migration of enslaved populations from Asia to the Cape Colony.(*52*) Distinguishing among Asian sources has been challenging in previous studies, potentially due to methodological constraints, limited sampling, genetic drift following initial admixture in South Africa, or a combination of these factors.(*13*, *23*) We find a noticeably higher signal of South Asian haplotypes in the Cederberg samples compared to the other sampled locations, likely due to their proximity to Cape Town (*Supplemental Figure 22)*.(*9*) Although our analysis yields ambiguous results for distinguishing among specific South Asian regional ancestries, further methodological developments could potentially refine these haplotype assignments.

In contrast to the ambiguous South Asian patterns, our analysis is able to provide further resolution to East/Southeast Asian ancestry components in the SAC. The asMDS for this component clearly point to Indonesia as the best proxy source population, rather than the mainland Asian populations (e.g. Han Chinese) typically used to capture this signal (Figure 4C).(*9*, *11*, *47*) Inferred proportions of East/Southeast Asian ancestry in our study populations is relatively low, ranging from trace levels (mean <1%) in the ≠Khomani San to about 6% in the Cederberg, following a South-to-North gradient as seen in some other ancestry components (*Supplemental Figure 9B and Supplemental Table 2*). We included only those whose combined maternal and paternal haplotype proportions reflected at least 10% Southeast Asian ancestry, which is a limited proportion of the study cohort. Additionally, input segments were required to have a minimum probability of 0.75 to be classified as East/Southeast Asian. Although the low proportion of East/Southeast Asian ancestry and the applied thresholds introduced some limitations, they also increased confidence in the results.

Detection of this Indonesian component in the Cederberg and other sampled regions aligns with historical records.(*53*, *54*) An estimated quarter of all enslaved individuals at the Cape Colony were of Malay or Indonesian descent.(*55*) The clustering patterns observed in our southeastern Asian asMDS correspond with major VOC trade routes connecting South Africa to predominantly western Indonesian islands.(*39*, *40*) Islands such as Sulawesi, Sumatra, and Timor had formalized ‘slave-clauses’ with the Dutch in the 1600s.(*34*) Interestingly, we see higher proportions of the South Asian component than Southeast Asian, approximately double the Southeast Asian inferred global ancestry proportions across populations. This pattern may reflect enslaved individuals from Indonesia remained concentrated in the central Cape Colony rather than dispersing northward, or that those who did migrate north, whether through forced relocation or escape, experienced demographic decline resulting in their minimal genetic contribution to the SAC populations outside of present-day Cape Town.

### SAC populations retain high Khoe-San ancestry

The VOC initially relied on livestock trade with Indigenous populations during the first 30-40 years of their occupation at the Cape Colony, but made little effort to expand or incorporate Indigenous groups into the colony.(*56*) However, as European colonists began expanding outward from the Cape Colony in the 17th century, they successively forced Indigenous populations into the interior, and more marginal areas such as the Kalahari and Cederberg mountains.(*6*) However, colonial ambitions intensified significantly in the 1730s, leading to heightened Indigenous resistance. A frontier war erupted between the colonists and the Khoe-San living in the Cederberg in 1739,(*57*) followed by significant population decline due to smallpox epidemics. Historical records indicate that many surviving Khoe and San individuals were forced into servitude and employed as laborers on settler farms.(*58*) In some cases, these Indigenous individuals admixed with those enslaved by the VOC, forming new communities that lead to people who largely self-identify as SAC today.(*59*)

The systematic displacement and marginalization of Khoe-San populations during and after the initial European colonization has led to a widespread belief in contemporary South Africa that Khoe-San ancestry is largely absent from modern populations, yet our analysis demonstrates substantial Khoe-San genetic contributions across study populations. Our results indicate that SAC populations from the Western and Northern Cape Provinces retain a substantial proportion of their genetic ancestry from Khoe-San populations, averaging 41–53%.(*11*) Haplotype-based analysis of Khoe-San segments (with >75% posterior probability) finds this component to be more similar to the southern Kalahari ancestry –– represented by the ≠Khomani San and Nama –– rather than the northern and central Kalahari, represented in this study by !Xoo and Ju|’hoansi (Figure 4). This result corresponds with the sampling sites’ geographic location south of the Kalahari (Figure 1 and 2C).(*48*) Previous studies have shown a clear north-south structure in Khoe-San genetic diversity among modern populations, with more recent work suggesting an even more geographically complex pattern, reflective of the Kalahari Desert as a barrier to gene flow.(*19*, *20*, *60–62*) Interestingly, the population centroids of the Karoo and Orange River in our asMDS results are closer to the ≠Khomani San than the Nama (*Supplemental Figure 18* and *Supplemental Table 5)*, with the Cederberg situated intermediate.

We hypothesize the observed patterns could result from one or more of the following: 1) European disruption led to widespread displacement and increased gene flow among Khoe and San populations south of the Kalahari, resulting in a Khoe-San ancestry signature generally intermediate between the ≠Khomani San and Nama. 2) A gradient of ancestry between the Nama and ≠Khomani could reflect a demic diffusion of Khoekhoe pastoralists both along the western coast and along the Orange River into the interior. If the diffusion incorporated local hunter-gatherer populations, rather than with replacement, ((*63*) such a process could fit the asMDS. 3) Finally, the Richtersveld Nama have experienced a severe post-colonial bottleneck; it is possible their extreme MDS values represent localized drift separating them from similar groups (such as Cederberg Khoe-San). Future research collecting additional samples from individuals in Namibia, and formal demographic modeling is necessary to address these hypotheses.

### Colonial Impact on Ecogeographic Population Structure

Colonial-era processes, centered at Cape Town and extending outward through systematic displacement and the VOC trade network, have created spatial gradients in the distribution of the five main ancestry components. Using ADMIXTURE *k*=5 results, we considered how these ancestry proportions varied across sampling regions. We observe a gradient of increasing Khoe-San ancestry with increased distance from Cape Town, following colonial patterns of systematic displacement of the Khoe and San groups, ((*6*) but observe the opposite in European, South Asian and southeastern Asian ancestries (Figure 3 and *Supplemental Figure 9*). We find higher proportions of European and Asian ancestries in individuals closer to Cape Town, likely because Cape Town was the initial heart of the settlement.(*3*, *9*) Unlike the other ancestries, the proportion of non-Khoe-San African ancestry has no relation with the sampling site’s distance from Cape Town (*Supplemental Figure 9*), reflective of the demographic history mentioned above.

In addition to these geographic gradients of ancestry, our results reveal how European colonization fundamentally altered population movement and structure across the landscape. Using effective migration rate analysis of shared identical-by-descent tracts among our study populations, we compared two distinct time periods: pre-colonial (approximately 450 CE to 1650 CE; 56-13 generations ago) and colonial (1650 CE to present; <13 generations ago).(*55*, *64*) During the pre-colonial period, we observe higher migration rates along the Cederberg and within the arid Greater Karoo interior, with natural barriers to movement between these regions (Figures 1 and 5A). The Cederberg’s rugged landscape may have constrained population movement, while the comparatively flatter semi-arid Karoo allowed for greater mobility within the region.

Colonial contact dramatically altered these patterns, resulting in an overall decrease in migration rates (Figure 5C). European colonization systematically restricted Indigenous movement through natural resource monopolization, policies limiting individual mobility, and military outposts protecting colonial boundaries.(*6*, *65*) Colonial control of water resources became a particularly powerful tool for controlling Indigenous populations, limiting Khoe-San movement and undermining pastoralist autonomy.(*6*) This shift is reflected in our environmental modeling: water resources show low importance in predicting pre-colonial migration rates, but the presence of lakes becomes significantly more important post-contact, potentially reflecting increased sedentism during the colonial period (Figure 5B, Figure 5D).(*58*)

As the British took control of the Cape Colony from the Dutch in the early 19th century, land became property of the Crown and Indigenous peoples were viewed as obstacles to colonial success.(*66*) Legal restrictions further constrained mobility— the colonial government’s 1809 law requiring Khoe-San individuals to adopt sedentism institutionalized the restriction of movement and facilitated labor control.(*67*) These policies systematically dismantled the traditionally mobile lifestyles of Khoe and San populations, forcing cultural assimilation as European colonizers imposed colonial norms while devaluing Indigenous lifeways as “primitive.” The persistence of this disruption is evident today, as most Khoe-San descendant individuals speak Afrikaans and English rather than traditional click-phoneme languages.(*48*)

## Conclusion

This study provides the first large-scale characterization of genetic composition across multiple regions in South Africa revealing that Khoe-San ancestry constitutes a significant proportion of ancestry among SAC populations (41–53%) from Cederberg, Great Karoo, and Middle Orange River. Despite historical colonial disruption, our results show that the Khoe-San descendant communities retained a significant portion of Khoe-San genetic ancestry. Utilizing local ancestry assignment, we are able to further pinpoint possible sources for colonial-era migration into South Africa. For example, we identify Indonesia as the likely source population for the previous “East Asian” genetic component, further supporting the historical record that a fourth of all enslaved individuals at the Cape Colony were of Malay and Indonesian descent. This ancestry remains a small minority in our sampled individuals, however. The substantial Khoe-San ancestry persisting in modern populations, the geographic gradients reflecting colonial settlement patterns, and the disrupted mobility patterns evident in contemporary genetic structure all demonstrate how colonial processes fundamentally altered the demographic landscape of South Africa. We emphasize that genetic signatures alone do not fully capture the complex power dynamics that shaped these demographic changes. Understanding the genetic legacy of colonization requires recognizing both what these patterns reveal about historical processes and the limitations of genetic data in capturing the full scope of colonial impact.

## Methods

### Sample Collection

Genetic and demographic data from several towns in the Cederberg, Karoo, and Orange River were newly collected between 2017 and 2023; locations are indicated in Figure 1. DNA from saliva samples were collected with Oragene OGR-600 kits (DNA Genotek) and ethnographic information regarding ancestry, language, and parental place of birth were collected for all participants. Institutional review board (IRB) approval was obtained from Stony Brook University (727494), University of California – Davis (1220180–7) and the Health Research Ethics Committee of Stellenbosch University (N11/07/210 and N11/07/210A). All individuals gave signed written and verbal consent with a community witness present before participating. Participants were asked to self-identify without categorical prompts. Prior to sampling, municipal or non-profit leaders in each community were consulted on the aims and approach of the project. Community level results were returned to the majority of towns between 2023-2025, including community meetings, pamphlet and poster distribution paired with discussions involving local municipalities, high schools, and community members.

### Ethnic composition of study participants

Study participants were recruited from communities of predominantly Coloured census, the Middle Orange River, Cederberg, and Karoo (Figure 1). To avoid racial categories, participants were asked to self-identify their ethnicity without prompts/examples from interviewers. 79% of participants chose Coloured as their primary ethnicity, whereas 12% chose either “Khoe-San” or a specific Khoe-San descent ethnicity, such as Nama. The remaining individuals chose a Bantu-language ethnicity (<4%) or provided a more complex answer such “I am South African” or “I am mixed”. The self-identified ethnicity of participants included in this study from a clinical cohort of the Middle Orange River region are 88.4% Coloured, 4.6% Tswana, 4.2% Khoe-San (e.g., Nama, San), and 1.9% as “other”.(*68*)

### Sample consolidation, Genotyping, and PCA

Newly collected samples were genotyped for >2.2million SNPs using the Illumina H3Africa array. Raw genotype data was processed using Illumina’s GenomeStudio to call common variants (MAF>0.05) on the autosomes, X– and Y-chromosomes, followed by zCall to call rare variants. Details of this pipeline are available at https://github.com/hennlab/snake-SNP_QC. All datasets were aligned to the 1000 Genomes Phase 3 reference panel and all ambiguous variants were removed.

We consolidated a reference panel incorporating data from four published genome sequencing projects, the African Genome Variation Project(*69*) (AGVP), Human Genome Diversity Project(*70*) (HGDP), Indonesian Diversity Genome Project(*71*) (IDGP), and 1000 Genomes Project(*72*) (1000GP), an unpublished cohort of 94 ≠Khomani San (KHM) whole genome sequences [Soto et al., in preparation], and previously published African samples(*73*) genotyped using the Omni5 array.

We extracted populations of interest from each dataset listed above. The Zulu, Omoro, Amhara, Wolayta, Somali, Nama (NAM), and Gumuz were extracted from the AGVP(*69*). Indonesian samples were sourced from the IDGP(*71*). The British (GBR), Iberians from Spain (IBS), Han Chinese (CHB), Vietnamese (KHV), Yoruba (YRI), Luhya (LWH), Bengali (BEB), Tamil (STU), and Gujarati (GIH) were obtained from 1000GP(*72*). Lastly, the French were obtained from HGDP and the Botswanan San from previously genotyped African samples.(*73*) The Botswanan San include two San populations, the Ju|’hoansi and !Xoo. However, the published dataset(*73*) did not include population IDs, leaving us to infer these IDs using a combination of methods further detailed the supplement (*see Supplemental Text and Figures 1–4*). The Botswanan San samples were processed before the lifting or merging with the larger dataset in order to maximize the variants used in the analysis. This left the Botswanan San on hg19 with 2,993,754 variants, and the NAM, KHM, ORG, CDB, NCTB, and KRO aligned to hg38 with 1,015,964 markers. Sample sizes for each population can be found in supplemental table 1.

Datasets were filtered for biallelic variants congruent with markers assayed using Illumina’s H3Africa array. With the exception of the included 1000GP and HGDP populations, each dataset was lifted from hg19 to hg38 using Picard Tools LiftoverVCF.(*74*) Samples were then merged with the newly collected cohort of South African samples genotyped on H3Africa.(*75*) The newly collected samples include SAC individuals sampled along the Cederberg Mountains (CDB), Karoo (KRO), Orange River (ORG), as well as Northern Cape Tuberculosis Project (NCTB). The merged data was filtered for Hardy Weinberg Equilibrium (<0.05), minor allele frequency (<0.00047 to remove singletons and doubletons) and individual and SNP missingness (>5%) using plink (v1.9). The compiled dataset contained 806,460 autosomal single nucleotide polymorphisms (SNP) and 1,165 individuals. Merging was assessed using PCA’s 1-10, with PC loadings generated through smartPCA(*26*). VCF files were converted to ‘.ped’ and ‘.map’ files for eigensoft smartPCA input.

### Phasing

The dataset was phased using the program Shapeit4 (v4.2.2)(*76*) and the Consortium on Asthma among African Ancestry Populations (CAAPA2) reference panel (unpublish data). This reference panel includes 78 ≠Khomani San and 49 Nama individuals, in addition to other continental African populations, making it the most representative reference panel available for the individuals sampled in this study. All Shapeit4 parameters were set to default, with the exception of ‘––mcmc-iterations’ being set to ‘10b,1p,1b,1p,1b,1p,1b,1p,10m’.

### Identifying Relatives

We used Parent Offspring Pedigree Inference Robust to Endogamy (PONDEROSA), a software designed to identify relatives within more endogamous populations and/or populations that have undergone recent bottlenecks, resulting in elevated levels of shared identical-by-decent (IBD) regions.(*77*) The program provides an option to simulate related individuals from the population for various relationships classes (i.e. maternal/paternal half-siblings, maternal/paternal grandparents, avuncular relationships). It accomplishes this using integrating the program ped-sim.(*78–80*) The IBD distributions for the simulated relationships are then used to train the model that will be applied on the real data.

PONDEROSA was independently preformed on each of the southern African populations, including the newly collected SAC individuals. We segregated the ≠Khomani San, Nama, CDB, KRO, ORG, and NCTB samples by region/population and proceeded to run simulations on each of the 6 southern African groups included in this study.

With the exception of the Botswanan San, samples were lifted from hg38 to hg19, without variant loss, in order to use the PONDEROSA this required a sex-recombination map aligned to hg19.(*78–80*) IBD calling was then carried out using PONDEROSA’s built-in script initiating phasedIBD(*81*), all settings set to default. We simulate 200 pairs of individuals for each population and then use the training classifiers to identify relatives in the populations. Relatives having a greater than 80% probability were assigned a specific class, while those below the threshold were assigned either 2^nd^, 3^rd^, or 4^th^ degree relatedness. Finally, we used PONDEROSA ‘remove_relatives.py’ script to filter individuals who were related closer than 3^rd^ degree. This script does not remove all relatives but rather retains one of the related pair of individuals, in order to maximize the number of unrelated individuals. For example, if a sibling pair existed in one of the populations, the function would retain one of the siblings in the unrelated subset of samples, while placing the other in the relative’s subset.

### ADMIXTURE

Prior to running ADMIXTURE, we first identified all first– and second-degree relatives using KING(*82*) and partitioned relatives into multiple non-overlapping “running groups”. We then performed separate ADMIXTURE analyses for each running group. ADMIXTURE assumes that all samples are unrelated because shared IBD tracts between close relatives can bias allele-frequency estimates and consequently inflate or deflate inferred ancestry proportions. Excluding close kin from each run ensures unbiased allele-frequency estimates, while systematically rotating relatives across groups guarantees that every individual contributes to at least one model fit.

The three Khoe-San reference populations were independently assessed for relatedness using PONDEROSA (as above). An internal R script was utilized to generate: 1) an unrelated subset of individuals and 2) randomized groups of first– and second-degree relatives while ensuring minimal overlap of related individuals across groups 3) a list of unrelated individuals in the dataset. A total of 10 groups were created to divide relatives; each then merged with the larger list of unrelated individuals. Corresponding PLINK files were generated for each of the final 10 running groups (RG).

The three southern African reference populations underwent a similar process independently of one another. However, instead of partitioning all first– and second-degree relatives into different running groups, we retained one individual from each related pair in the unrelated subset using PONDEROSA ‘remove_relatives.py’ script. Individuals classified as relatives were divided into 10 randomized groups and added to their respective RG PLINK file with the. This method allows us to retain a maximal number of Khoe-San individuals in the core, an unrelated list of individuals shared across the RG, while the related individuals are randomly subset across the 10 running groups with little overlap in relatives. A flowchart of this pipeline is included in Supplemental Figure 5.

European and Asian populations were thinned to only include the GBR, IBS, GIH, BEB, SES, CHB, KHV, and Indonesians in admixture. Linkage disequilibrium (LD) was calculated on the merged dataset of 806,460 markers, including the reference populations and newly genotyped southern African samples using PLINK 1.9(*83*). Data were conservatively pruned using the PLINK ––indep-pairwise flag for markers with LD of 0.5 or greater within a window size of 50 SNP, with sliding window size of 20 SNPs. The pruned dataset contained 479,497 variants. Ten iterations of ADMIXTURE *k*=4:10, *k* being the number of clusters ADMIXTURE infers, were performed for each running group, all with fivefold cross-validation and randomly selected starting seed. With respect to the level of k, results for each running group were averaged across individuals. ADMIXTURE results were visualized using pong(*84*) (Figure 2A, Supplement Figures 6 and 7).

### Local Ancestry Inference

Local ancestry (LAI) was inferred using Gnomix(*32*), which has been shown to outperform other programs in accuracy and speed. Similar to other LAI programs, Gnomix treats each reference population as an independent ancestral group, without considering the relationships between populations. In other words, the individuals within each ancestral group must have a high ancestry proportion with respect to the ancestral group they are representing.

This was relatively simple requirement to meet for most of the references, but particularly difficult when using reference populations who are admixed, such as the southern African reference individuals. Based on the ADMIXTURE results for *k*=6, we filtered for ≠Khomani and Nama individuals having at least 85% Khoe-San ancestry, and the Botswanan San for minimum 90% Khoe-San ancestry. We chose to use varying cut-offs to retain the maximum number of individuals while accounting for within Khoe-San population structure and differences in sample sizes. Southern African individuals below the cut-off were then added, in addition to the Zulu as a control.

Given that the SAC populations are often described as five-way admixed, we assembled a LAI reference panel that accounts for each putative ancestry. Putative source populations for the KHS components in our study samples include the ≠Khomani San, Nama, and the Ju|’hoansi and !Xoo from Botswana.

Putative source populations for European ancestry include GBR and IBS (EUR), for South Asian ancestry include BEB, GIH, and STU (SAS), and for East/Southeast Asian ancestry included CHB, KHV, and Indonesians (EAS). We separated the KHS populations, that is the Nama, ≠Khomani San, and Botswanan San, from other Africans and aggregated all NKA populations into an independent category. We opted to use this method because 1) the Khoe-San are highly diverged from other NKA populations, providing confidence that we can distinguish between Khoe-San and NKA haplotypes, and 2) the study samples likely have NKA ancestry from various source populations, making it difficult to discriminate between the putative populations. Populations included in NKA are the Yoruba, Luhya, and Gumuz. The Omoro, Amhara, Wolayta, and Somali individuals were removed from the NKA reference panel due to the back-to-Africa(*85*) ancestry which weakened the model’s assignment of EUR and NKA segments. In summary, our five-binned local ancestry reference panel had populations representing EUR, SAS, EAS, KHS, and NKA ancestry.

Gnomix(*32*) was executed using the recommended array config, a logistic regression base and xgboost smoother modules. All settings were set to default, with the exception of adjusting for larger window (0.4cM) and smoother size (82). We assessed multiple models and found this model to have the most accuracy and to be more stable against phase switching due to the increased window and smoother size. These changes reduced sensitivity with decreased window size but increased model accuracy. Local ancestry results were evaluated using five methods: 1) a confusion matrix, 2) ADMIXTURE and Gnomix global ancestry correlations, 3) track lengths, 4) probability distribution for each ancestry, and 5) visualizing karyograms. See supplement for more detail on local ancestry assessment.

### Ancestry-specific MDS

We used multi-array ancestry specific multidimensional scaling (maaMDS)(*31*) to evaluate whether the projected ancestry-specific haplotypes produced by GNOMIX in our southern African study samples would gravitate towards one or more putative source populations relative to others. This method calculates the dissimilarities among the assigned reference populations, then incorporates ancestry-specific haplotypes from the study population by their distances relative to the existing reference clusters, effectively projecting them onto the pre-defined coordinate axes. The analysis was carried out using average pairwise genetic distances, including only haplotypes with at least 75% probability of the target ancestry. Each ancestry had an independent maaMDS run where only the respective putative reference populations were specified. For example, when running the non-Khoe-San African (NKA) maaMDS, the KHS, EUR, SAS, and EAS populations were removed from the analysis. The flag ‘IS_WEIGHTED’ was set to true. Reference populations were given a weight of ‘1’, while study samples were assigned ‘0.01’ in order to ensure they are projected within the MDS space of the reference haplotypes. Visualization of the output data was done using the ggplot2(*86*) library in R.

### Euclidean Distance Computation

Using the coordinates from the asMDS1 and asMDS2, we calculated the Euclidean distances between population averages distribution. Individuals were assigned to populations based on predefined labels, and for each population, the MDS1 and MDS2 centroid was calculated by computing the arithmetic mean of the individual MDS1 values. Pairwise Euclidean distances between population centroids were computed in one dimension (MDS1) using the absolute difference between the centroids. The final pairwise Euclidean distance matrix was visualized as a table. Using the gridExtra package in R, the formatted distance matrix was converted into a table with a custom theme.

### SPRUCE

To estimate and identify potential environmental factors contributing to gene flow, we used SPRUCE (Spatial Prediction using Random Forest to Uncover Connectivity among Environments)(*87*), a machine learning approach to integrate genetic and environmental data, to determine which factors may be influencing population structure in individuals from Nama, Khomani, and Khoe-San descendant communities. SPRUCE utilizes the MAPS output to extract environmental values from the coordinates corresponding to the produced demes.

MAPS (Migration and Population Surface estimation) ((*41*) estimates time-resolved dispersal rates and population densities based on the number of shared long Pairwise Coalescent Segments (lPSC). Prior to calling identical by descent (IBD) segments, we performed phasing with SHAPEIT4(*76*) removed first– and second-degree relatives using the output from PONDEROSA ((*77*) (see *Identifying Relatives* and *Phasing* methods). IBD segments were then called using hap-ibd v1.0(*88*) were repaired with the program merge-ibd-segments. For running MAPS, an IBD matrix was constructed using code provided by the SPRUCE authors and the IBD calls were split into two groups: 2-6 and greater than 6 cM. The categories were chosen based off of historical dates corresponding to pre– and post-European contact in South Africa. The mean ages of the categories are 56.25 and 12.50 generations. ((*64*) Following the recommendation of the SPRUCE authors, we chose to use 300 demes for higher resolution migration patterns. Migration surfaces were visualized using the R package plotmaps(*41*).

Climate data such as mean annual temperature, maximum temperature of the hottest month, minimum temperature of the coldest month, mean annual precipitation, precipitation in the wettest month, and precipitation in the driest month were sourced from the free, online repository CHELSA (Climatologies at High Resolution for Earth’s Land Surface Areas)(*89*). Slope data was obtained from the Geomorpho90m dataset,(*90*) and elevation was obtained from MERIT DEM (Multi-Error-Removed Improved Terrain Digital Elevation Models)(*91*). Processing steps were performed using the Geospatial Data Abstraction Library (GDAL/OGR Contributors 2021) in a Bash environment. The geographic and environmental data were clipped within the geographic range of interest and the demes.txt output file from the MAPS software.

The predictor variables for the random forest regression were the values of 10 spatial variables related to climate, ecology, and geography at each deme. All variables were numeric and continuous with the exception of lakes and rivers which were coded as “1” for a body of water being present and “0” for a body of water being not present. The response variable was calculated by using the migration rate at each deme from the “mRates.txt” file outputted by MAPS and converting them to 10^m^.(*87*) The R package randomForestSRC was used to perform the random forest regressions (https://CRAN.R-roject.org/package=randomForestSRC). We performed random forest regressions for both 56-13 generations ago (IBD segments greater than 6cM) and less than 13 generations ago (IBD segments between 6 and 2 cM), which were our time intervals of interest. All other model specifications and considerations were applied as described in Pless et al. (2023).

### Within and Between Population IBD Sharing

GERMLINE2(*92*) was used to call identity by descent (IBD) segments to investigate IBD sharing both within and between communities across South Africa. Phasing was performed using SHAPEIT (see *Phasing* methods). First and second-degree relatives were removed using PONDEROSA (see *Identifying Relatives* methods) and KING IBDseg. IBD segments were repaired using *merge-ibd-segments*.

GERMLINE2 reports all pairwise IBD segments for each chromosome. We filtered for IBD segments greater than 6cM and calculated the sum of the IBD segment lengths for each pair of individuals to get the total length of pairwise shared IBD across the genome. We then filtered the total length of pairwise shared IBD for segments greater than 13cM. A heatmap analyzing within and between community IBD sharing was generated in R using pheatmap (version 1.0.12). The mean of the total shared IBD segments was calculated for the heatmap to look at the average shared IBD within and between communities. IBD networks were generated in R using iGraph.

## Acknowledgments

We would like to thank all the participant communities in the Western and Northern Capes for their continued trust and support. We would especially like to thank our many community research assistants and translators who assisted in data collection for the project. This research was supported by a NIH grant R35GM133531 and R35GM133531-04S1 (to BMH), R01 AI151549 (to PJM and BMH), and NSF grant #2419417 (to SLE and AWR). The content is solely the responsibility of the authors and does not necessarily represent the official views of the National Institutes of Health. We acknowledge the UC Davis Genome Center High-Performance Computing Cluster for data analysis support. We also thank Alexander Ioannidis, Daniel Mas Montserrat, and Santiago Gerardo Medina-Muñoz for their help in assessing the local ancestry models, and Carmina Barberena-Jonas for guidance on running the ancestry-specific MDS analysis.

## Inferring Botswanan Populations

We began by running an unsupervised ADMIXTURE inferring 2 to 7 population clusters (k) with the entirety of the Crawford et al. (2017) data (Supplemental Figure 1), along with the GBR, BEB, STU, GIH, KHV, IND, YRI, HMB, LWK, Zulu, Gumuz, Somali, Wolayta, Omoro, KHM, and Nama, and query populations. From there, we subset the Crawford et al. (2017) data for the 314 individuals with the highest Khoe-San ancestry proportion (Supplemental Figure 1A), as that is the reported number of San individuals sampled in the study,^1^ and then re-ran ADMIXTURE k2 through k7 with those individuals (Supplemental Figure 1B).

We then filtered the Botswanan San, ≠Khomani, and Nama individuals with 85% or greater Khoe-San ancestry as inferred by ADMIXTURE k5 and plotted them on a PCA (Supplemental Figure 2A). We observe a previously reported cline among the Kalahari San populations, ≠Khomani, and Nama. The Botswanan San split into two clusters, one of which anchored a corner of PC2. From there, we were able to infer the cluster of Botswanan San individuals diving PC1 likely have greater genetic variation relative to the other high Khoe-San populations. This leads us to infer that cluster of individuals to be the !Xoo, who were reported to have a higher population ecective size relative to the Ju|’haonsi.^2^

We continued to our KHS ancestry-specific MDS. We add ≠Khomani, Nama, and Botswanan San individuals who did not meet the 85% Khoe-San ancestry cut-oc into the query. We project those haplotypes, along with all first– and second-degree relatives in the data set, onto an MDS with the 85% and greater KHS reference. Projecting these individuals minimizes the ecect relatives and admixture can have on dimensionality reduction. We use the mclust^3^ R package to read in MDS2 data for the Botswanan San, the dimension where the Botswanan San split into two clusters. This package is designed for complex clustering and classification using a Gaussian finite mixture, meaning it assumes the data is made up of several Gaussian distributions. We use it to model MDS2 distribution with two clusters and identify the mean and standard deviation of both clusters (Supplemental Figure 2B-D). We filtered our model results to include only individuals with a probability of 0.9 or higher of belonging to a cluster. Upon examining samples across both clusters, we found no overlap between individuals assigned to each cluster. Most individuals who did not meet the ≥0.9 probability threshold are those positioned between the two cluster peaks, i.e., more than one standard deviation away from the modeled mean. These 10 individuals were labeled as BSAN_UNK (Botswanan San, Unknown). The individuals within the cluster positioned on the right side of MDS2 were labeled !Xoo, as they represent the samples contributing most to the genetic variation in MDS1. This classification leaves the remaining cluster of individuals to be identified as Ju|’hoansi (Supplemental Figure 3).

In order to refine the clustering of the assigned BSAN_UNK individuals, we performed a dimensionality reduction with uniform manifold approximation and projection (UMAP)^4^ followed by genetic clustering with Hierarchical Density-Based Spatial Clustering of Applications with Noise implementation, HDBSCAN(!^).^5^ We used as input the first 100 principal components (PCs) obtained with SmartPCA program from EIGENSOFT 7.2.1^6^ and applied the same approach described in Diaz-Papkovich et al 2023.^7^ For visualization purposes, UMAP was run with 25 number of nearest neighbors, a minimum distance of 0.3, and a dimension reduction of 2 components. For clustering, UMAP parameters included 50 number of neighbors, minimum distance of 0.01 and 5 number of components. Next, HDBSCAN(!^) was used to extract clusters using an !^ of 0.3 and a minimum number of points in a cluster of 50.

**Supplemental Figure 1).**
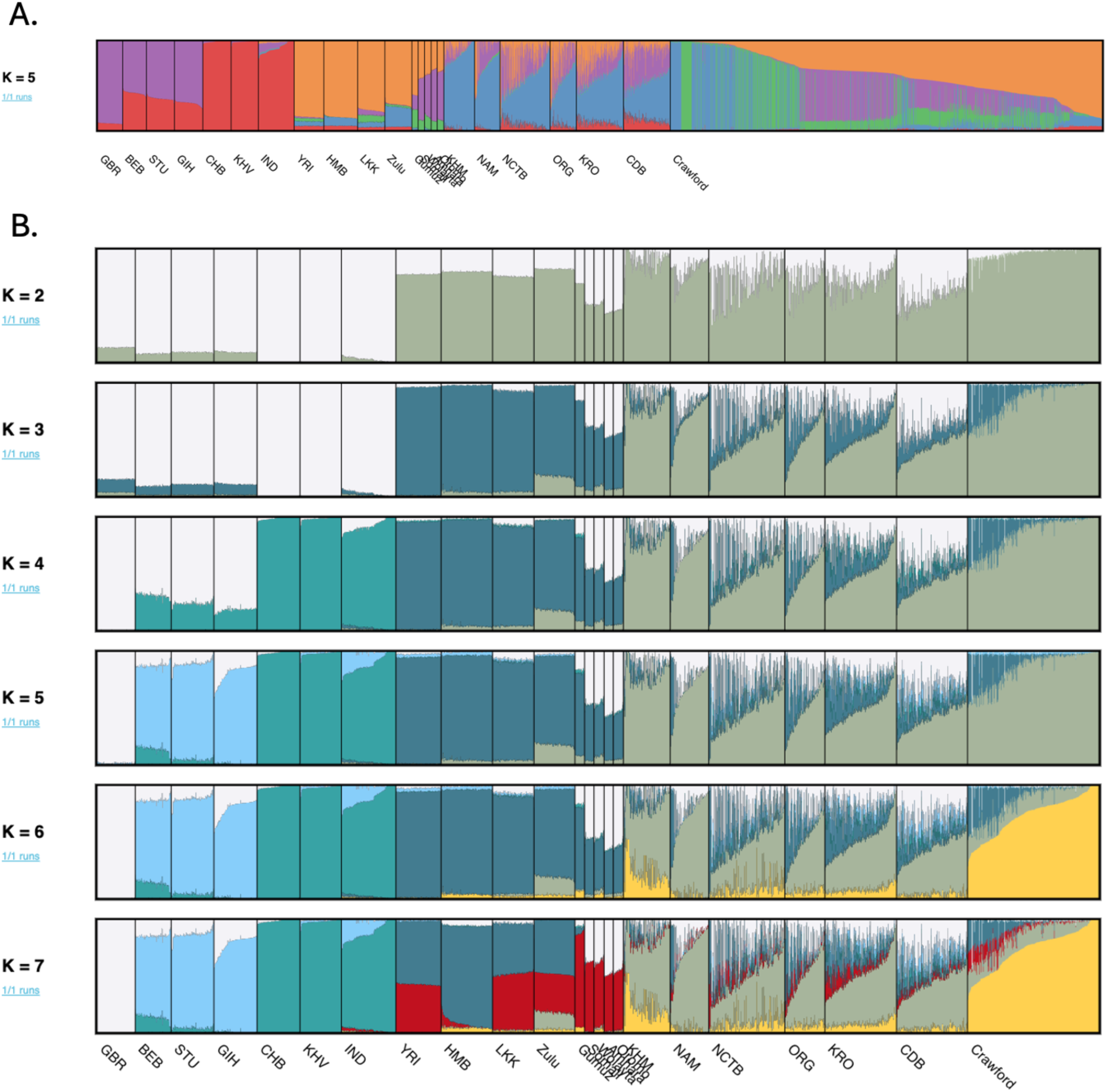
Identifying the Botswanan San using ADMIXTURE. **A)** ADMIXTURE at k5 with the entire Crawford et al. (2017) data. **B)** ADMIXTURE k2 through k7 with only 314 Botswanan San with the highest Khoe-San ancestry as inferred by figure A’s results.

**Supplemental Figure 2).**
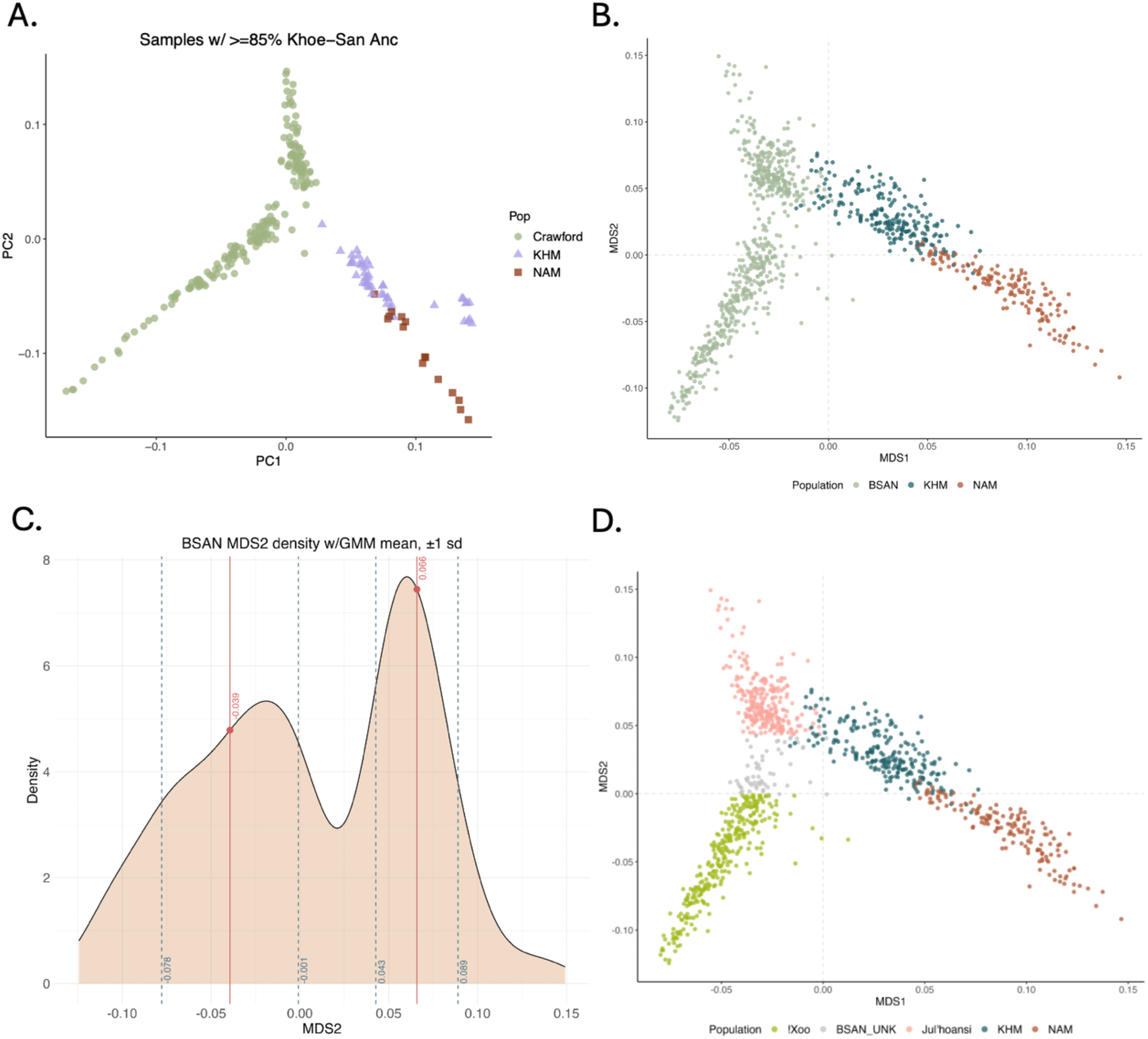
Dicerentiating Botswanan San Populations. **A)** Individuals with greater than 85% Khoe-San ancestry as inferred by ADMIXTURE k5 from Supplemental Figure 1A. **B)** Khoe-San (KHS) ancestry-specific multidimensional scaling (MDS) 1 and 2 of KHS reference populations. Individuals with less than 85% Khoe-San ancestry were added to the local ancestry inference query and projected into the MDS space, along with first– and second-degree relatives. **C)** mclust Gaussian mixture model identified mean and 1 standard deviation of MDS2 Botswanan San density. Red line specified averages while dashed blue lines highlight the 1 standard deviation cut ocs. **D)** Cluster/population assignment as inferred by the Gaussian mixture model from mclust R package and previously published population ecective size data^2^.

**Supplemental Figure 3).**
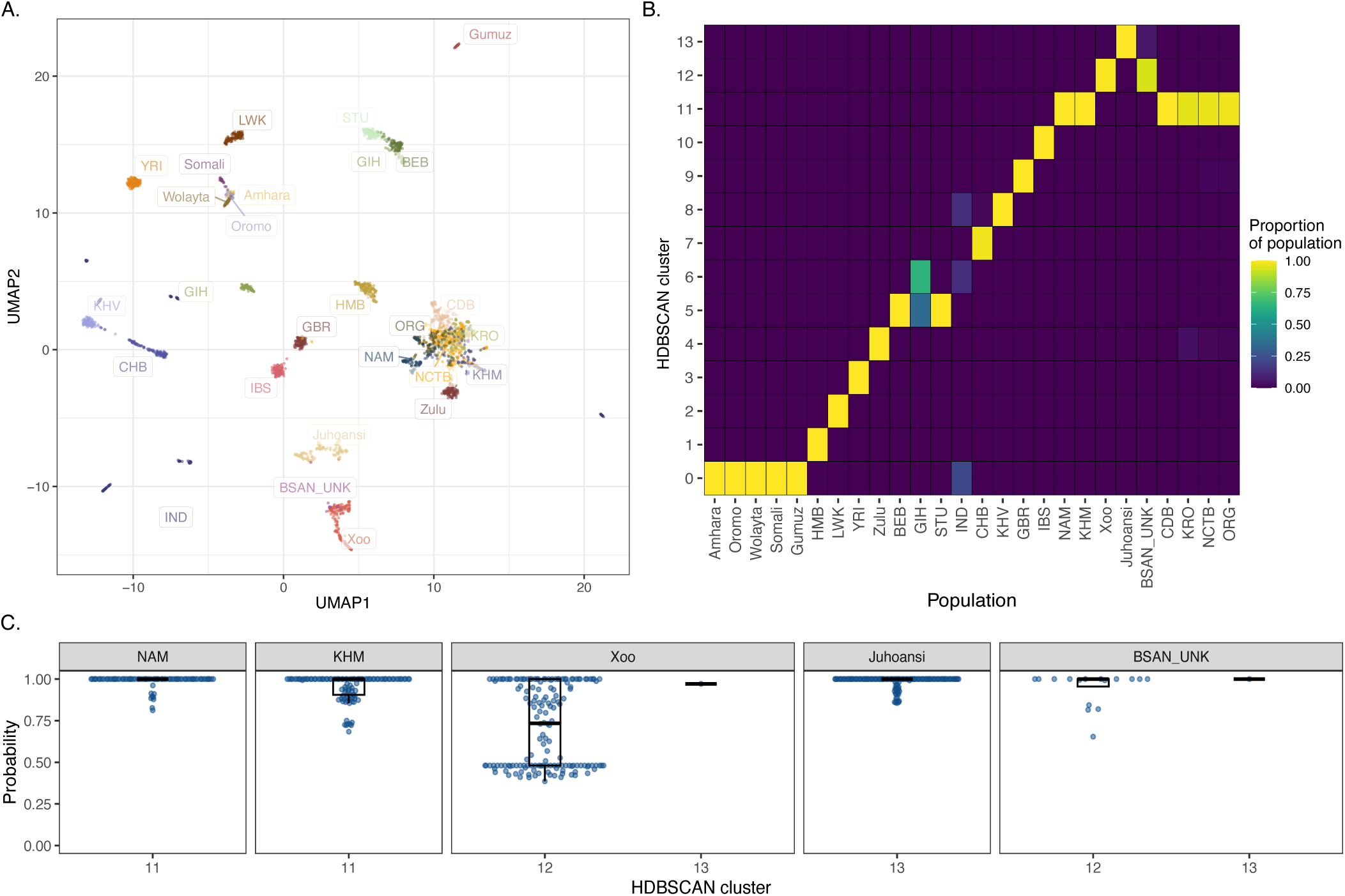
UMAP and HBDscan of reference and query populations. **A.** UMAP generated with 25 nearest neighbors, and minimum distance of 0.3. Labels refer to population labels. **B.** Proportion of individuals of each population contained within a given cluster inferred from HDBscan algorithm (see Methods for further details on the parameters used). **C.** Distribution probabilities per individual to be in the cluster inferred in B for each Khoe-San reference population.

**Supplemental Figure 4).**
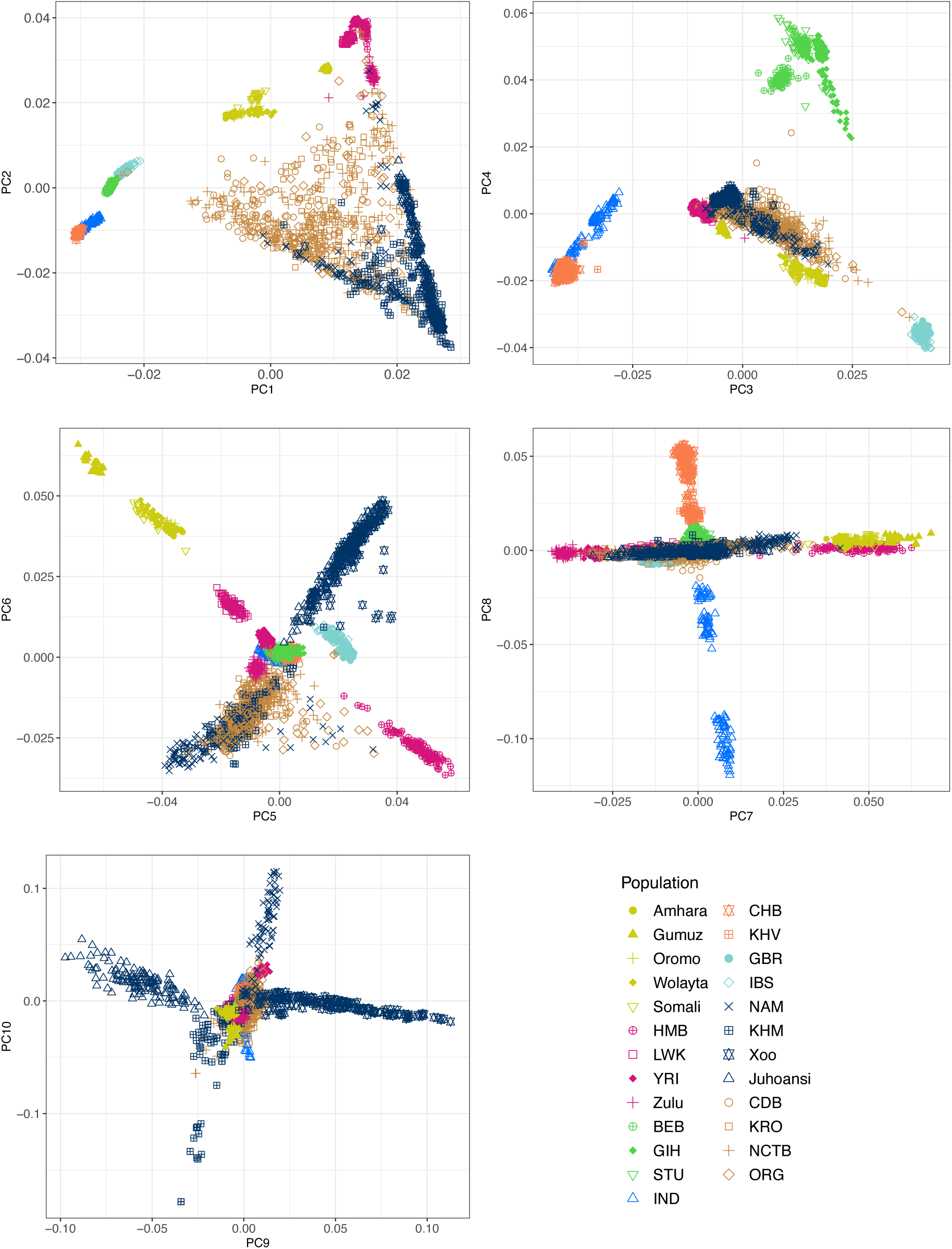
PC1 through PC10. PCs 1 through 10 were generated using smartPCA and plotted using ggplot2 in R. Colors roughly represent broader ancestry, while shapes represent population.

## Characterizing sampled Coloured populations

**Supplemental Figure 5).**
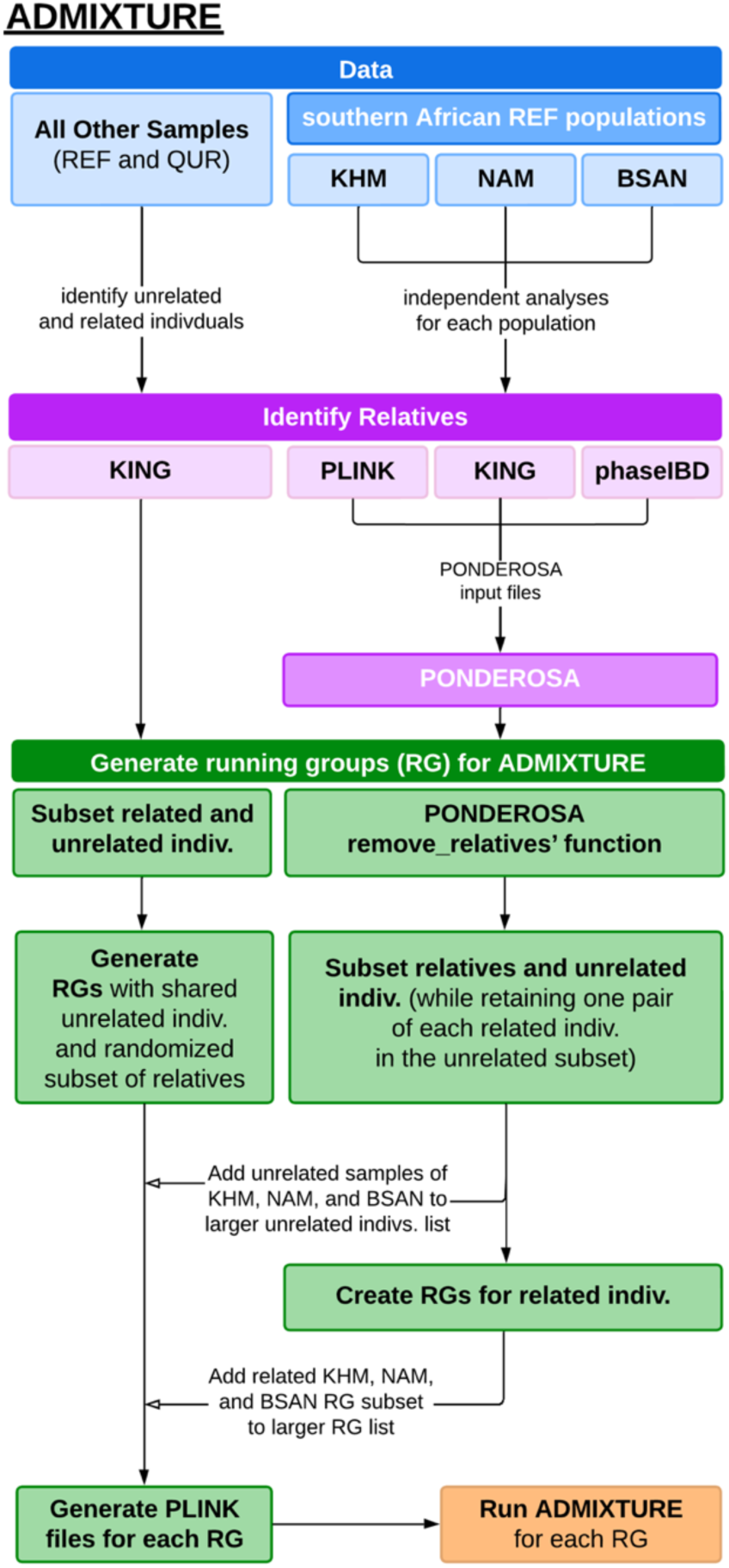
Identifying relatives’ workflow. Visual of applied workflow used to identify relatives and subset samples into running groups (RG) for ADMIXTURE. Each RG contained the same set of unrelated individuals but a dicerent subset of related individuals. They were all subject to 10 iterations of ADMIXTURE, all with a dicerent initial starting seed and cross-validated five times. PLINK LD pruning was completed prior to sub-setting samples, but after identifying relatives. PLINK LD pruning not depicted in the workflow.

**Supplemental Figure 6).**
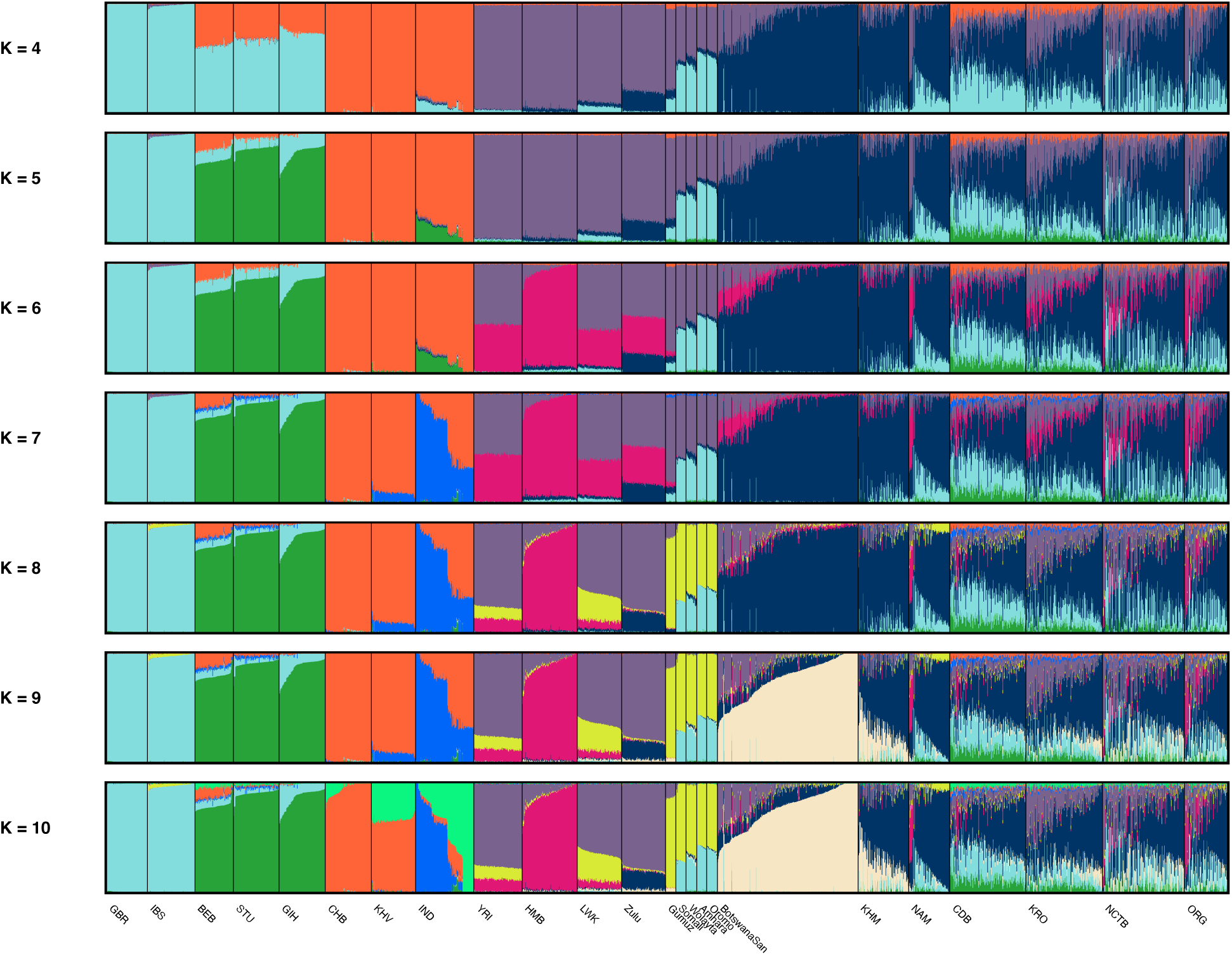
Unsupervised ADMIXTURE run of *k*=4 through *k*=10 using final merged dataset of 2,505 individuals and 806,460 markers congruent H3Africa array.

**Supplemental Figure 7).**
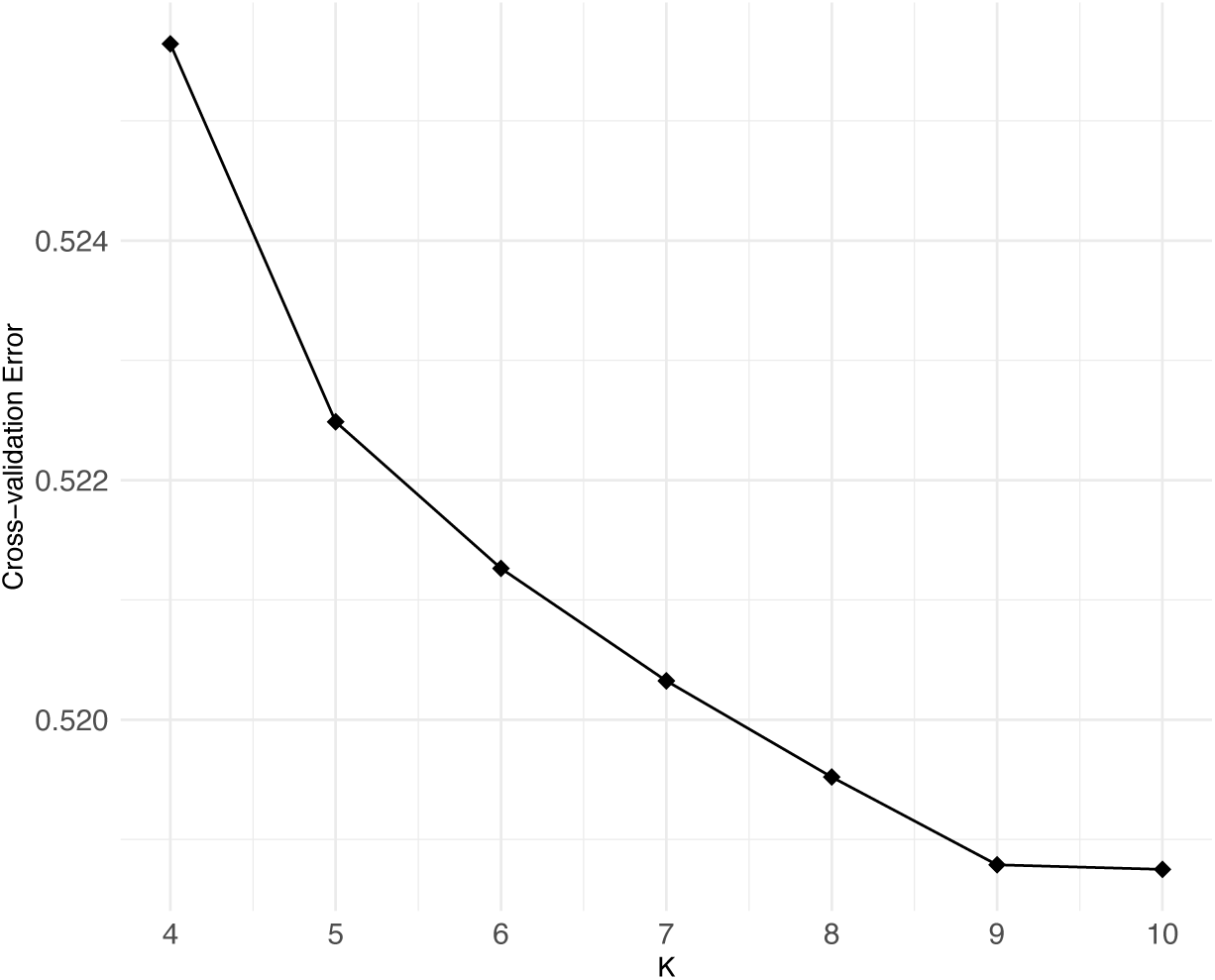
Cross-validation error rate for ADMIXTURE visualized in Supplemental Figure 6.

**Supplemental Figure 8).**
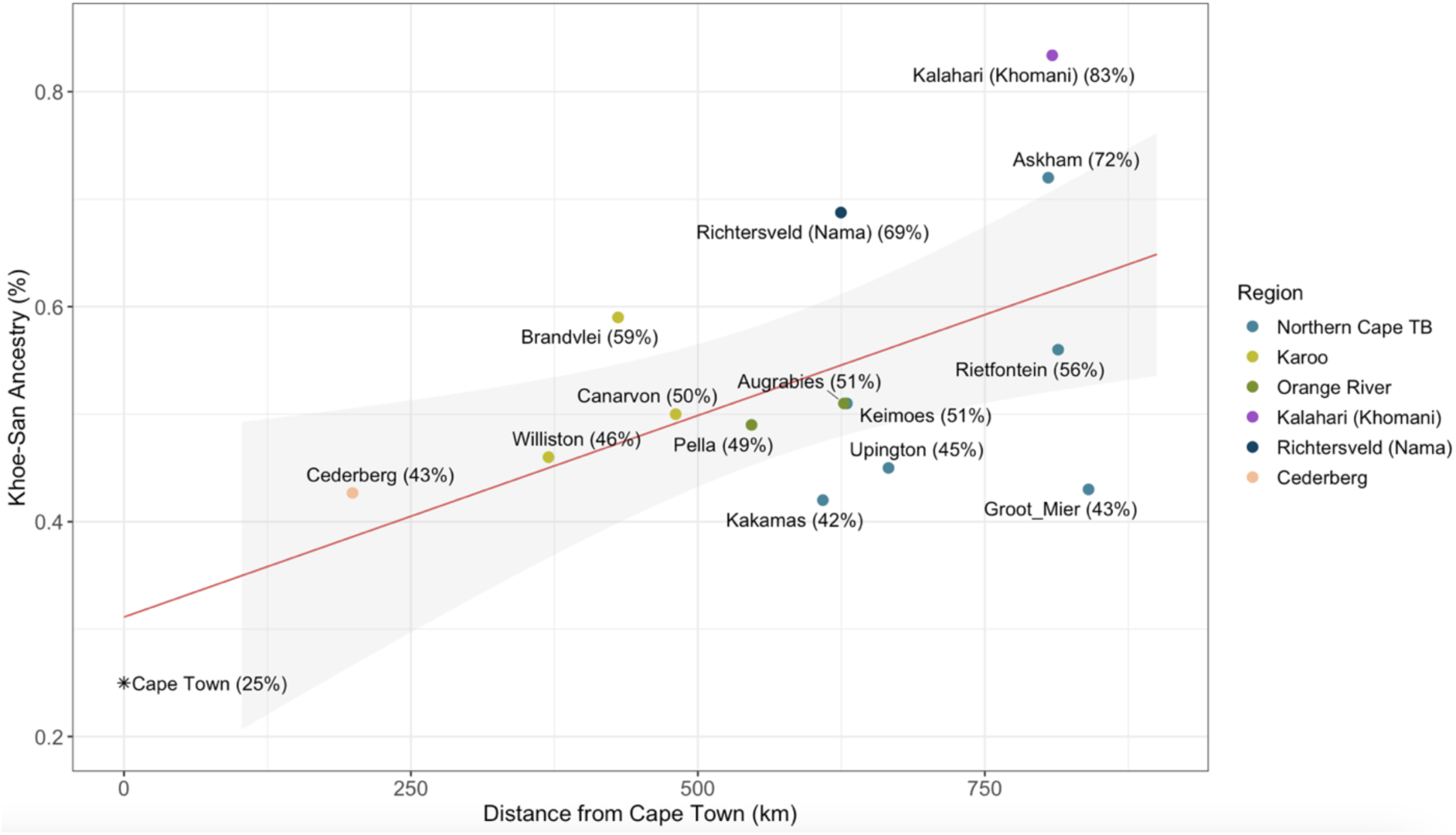
Proportion of Khoe-San ancestry by distance from Cape Town. Northern Cape Tuberculosis project participants are separated out by sampling site.

**Supplemental Figure 9).**
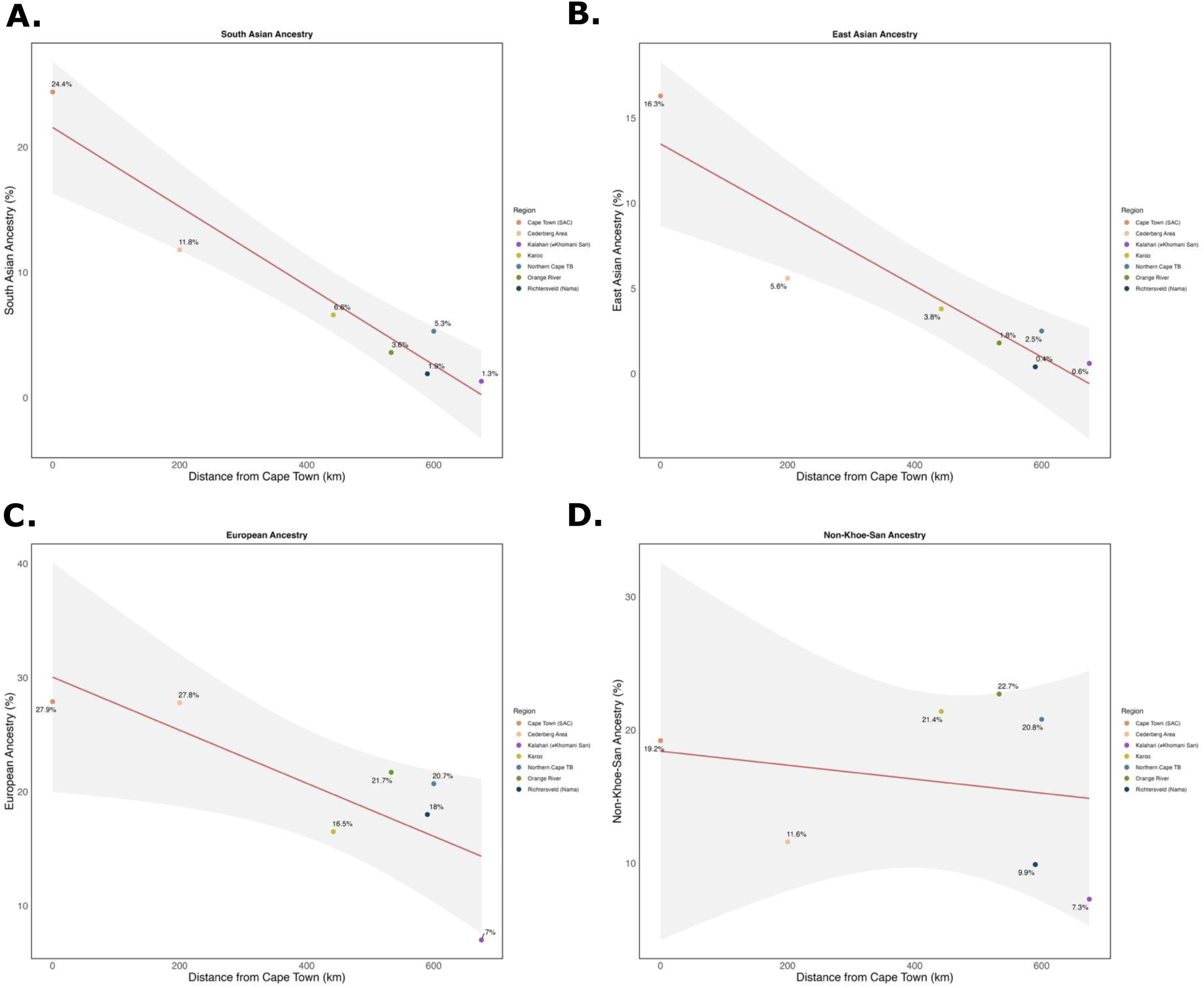
Linear regression showing ancestry by distance for. A) South Asian (R^2^ = 0.895), B) East Asian (R^2^ = 0.847), C) European (R^2^ = 0.0572), and D) Non-Khoe-San Ancestry (R^2^ = 0.004).

## Local ancestry results

Our model appears well-calibrated, with the confusion matrix showing >96% accuracy in predicting EAS, NKA, and KHS haplotypes, and ∼90% accuracy for EUR and SAS segments. Correlation between ADMIXTURE results for 5 population clusters and Gnomix global ancestry results is high for EUR, NKA, and KHS (r² ≥ 0.96), albeit slightly lower for smaller segments like EAS (r² = 0.94) and SAS (r² = 0.91), still indicating a strong correlation.

The highest local ancestry misassignment occurs between SAS and EUR ancestries, with 6.9% of SAS segments being assigned to EUR and approximately 3.5% of EUR segments assigned to SAS. This misassignment also contributes to the larger variance in EUR and SAS ancestry probability distributions (Supplemental Figure 11). We attributed this to the ‘Indian cline,’ referring to the geographical distribution of two primary ancestral groups across South Asia: the ‘Ancestral North Indians’ (ANI) and the ‘Ancestral South Indians’ (ASI)^8–11^. The ANI share closer genetic similarities with populations from the Near Eastern and European populations, while the ASI are more closely related to East Asians^8–11^, though many populations across South Asia retain a proportion of both ancestries. The ANI component among the BEB, STU, and GIH is visualized in our ADMIXTURE, highlighted in light green.

**Supplemental Figure 10).**
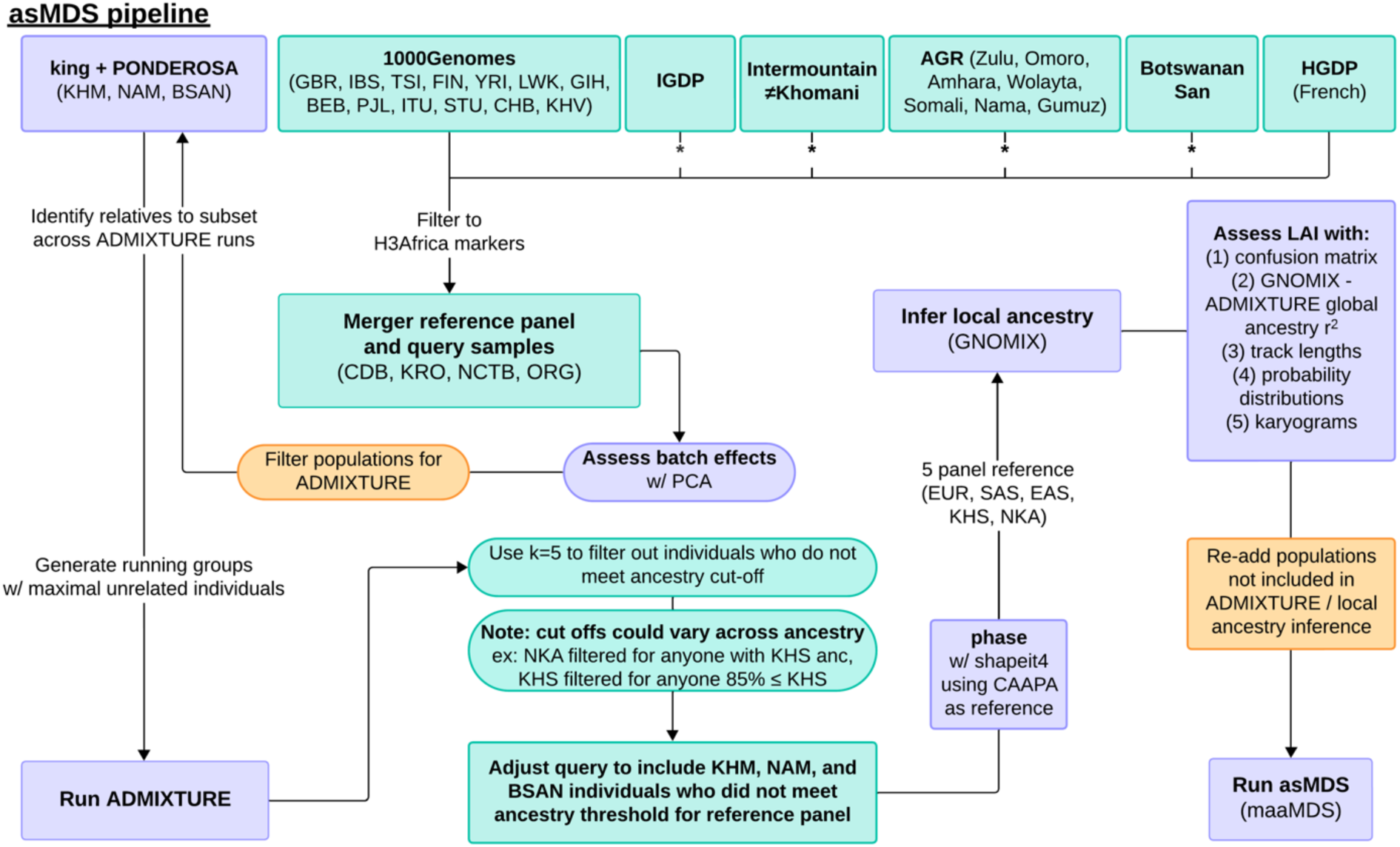
Local Ancestry and ancestry-specific MDS pipeline. Shapes in turquoise highlight steps taken regarding sample consolidation and filtering, while orange shapes are steps taken to filter or add entire populations. Shapes in purple major analyses executed throughout the pipeline. The * within lines represent whole genomes that were lifted from HG19 to HG38.

**Supplemental Figure 11).**
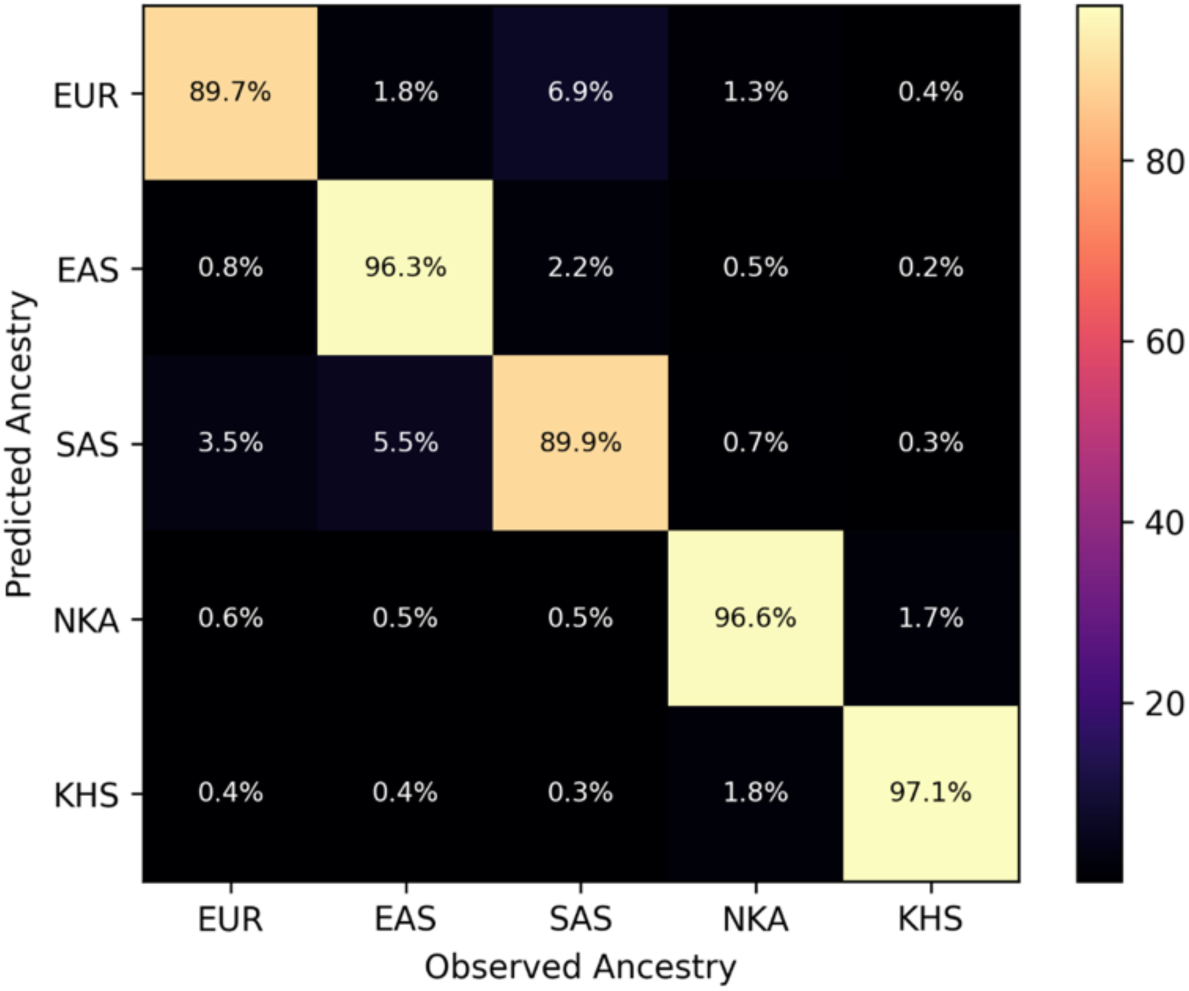
Local ancestry inference confusion matrix normalized across all chromosomes, where the x-axis is the ancestry observed, and the y-axis is the model’s predicted ancestry. The heat map highlights the accuracy of our model.

**Supplemental Figure 12).**
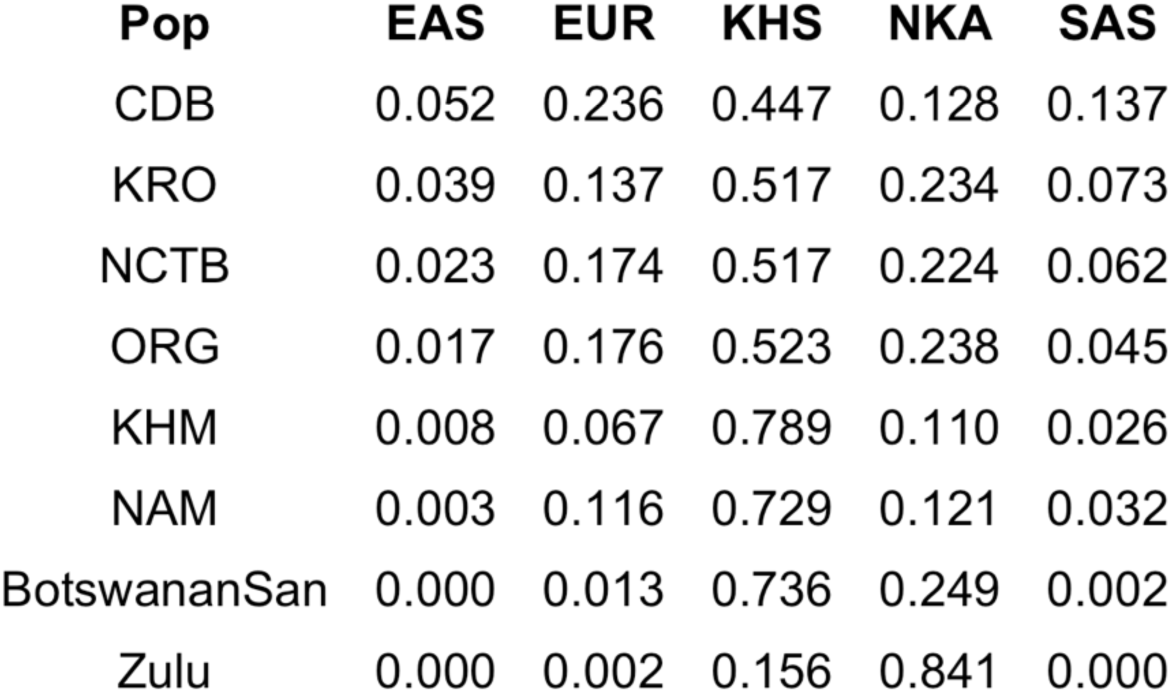
GNOMIX population-specific global ancestry average proportions.

**Supplemental Figure 13).**
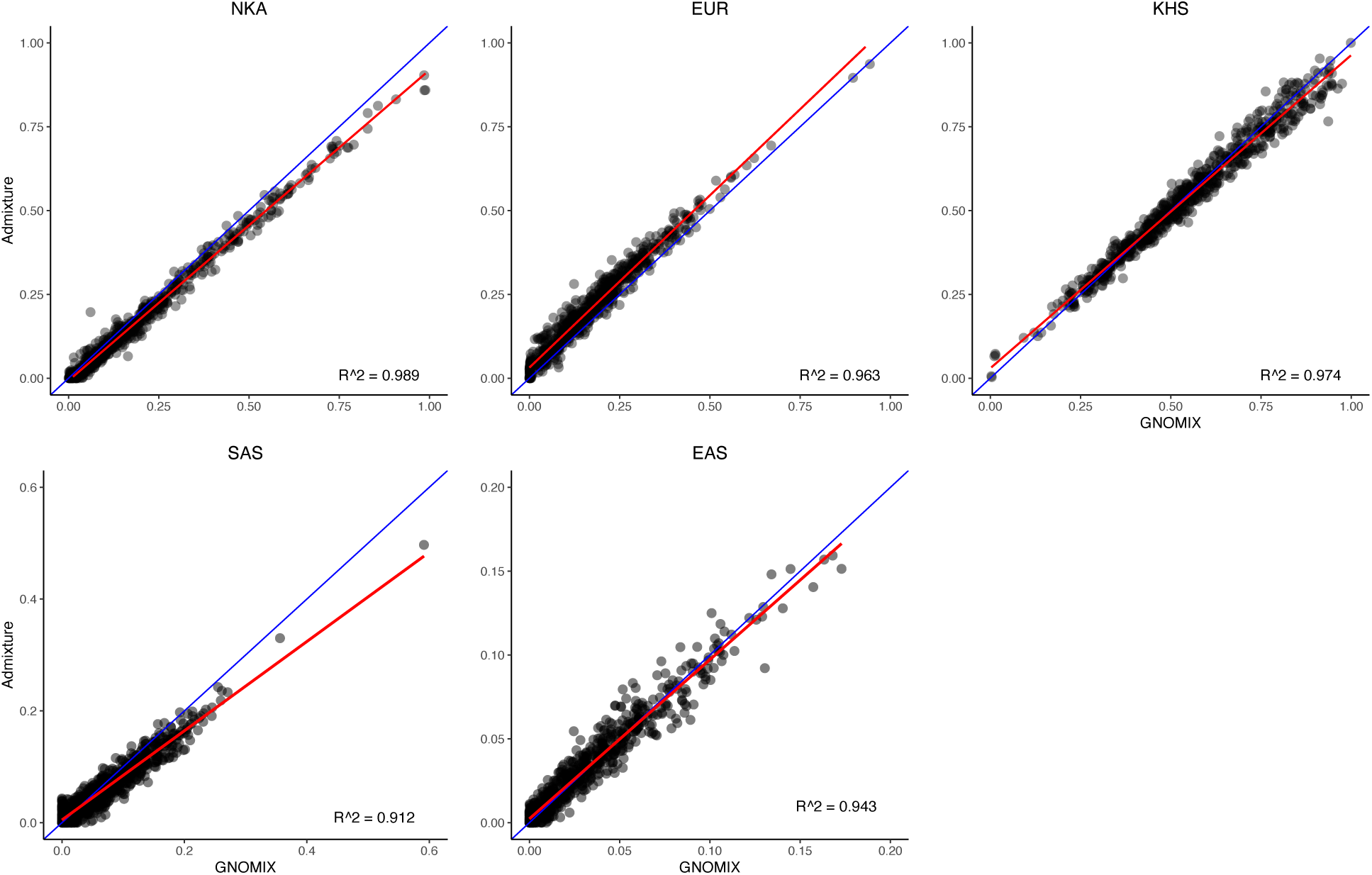
Correlation between GNOMIX global ancestry results and ADMIXTURE, where the red line is the best fit and blue line is x=y. R-squared values for each ancestry is reported on the bottom right of the respective plot.

**Supplemental Figure 14).**
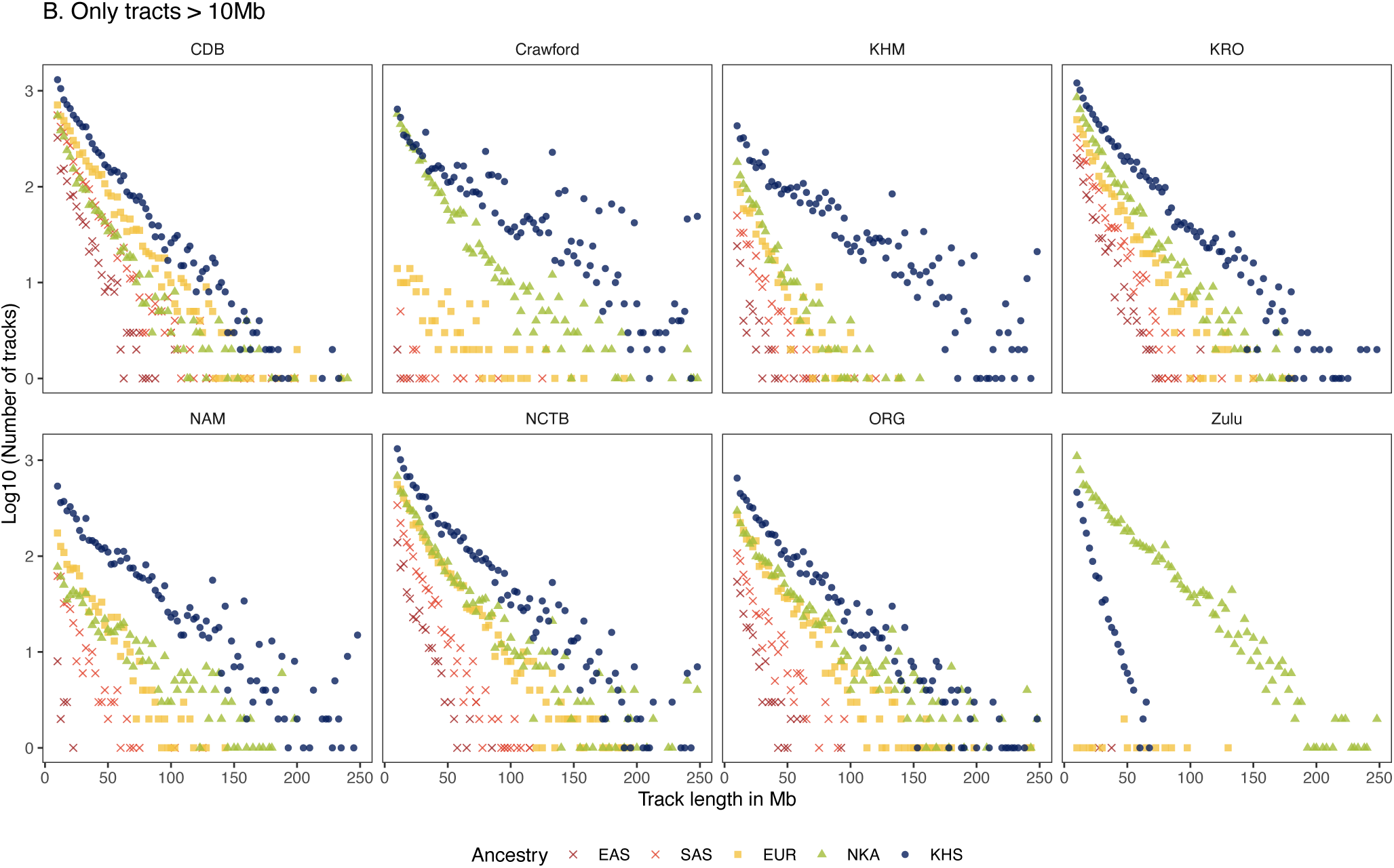
Track lengths with at least 75% probability of the respective local ancestry and greater than 10Mb across query populations. The x-axis is the track length in Mb and the y-axis is the log10 of the number of tracks.

## Ancestry-specific MDS results

**Supplemental Figure 15).**
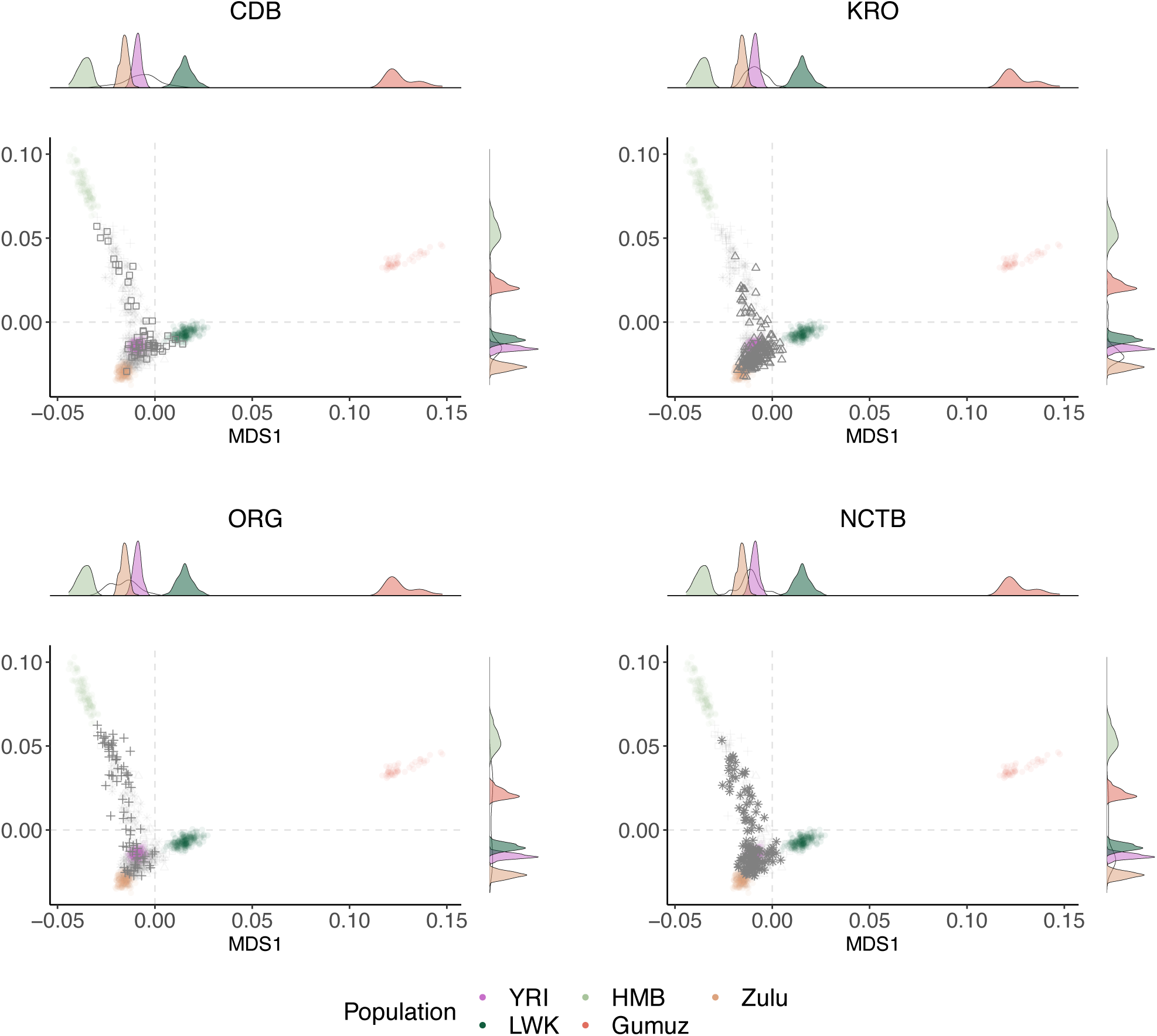
NKA asMDS results by population. Density plots for MDS1 and MDS2 are provided with the respective query population density visualized using a transparent line.

**Supplemental Figure 16).**
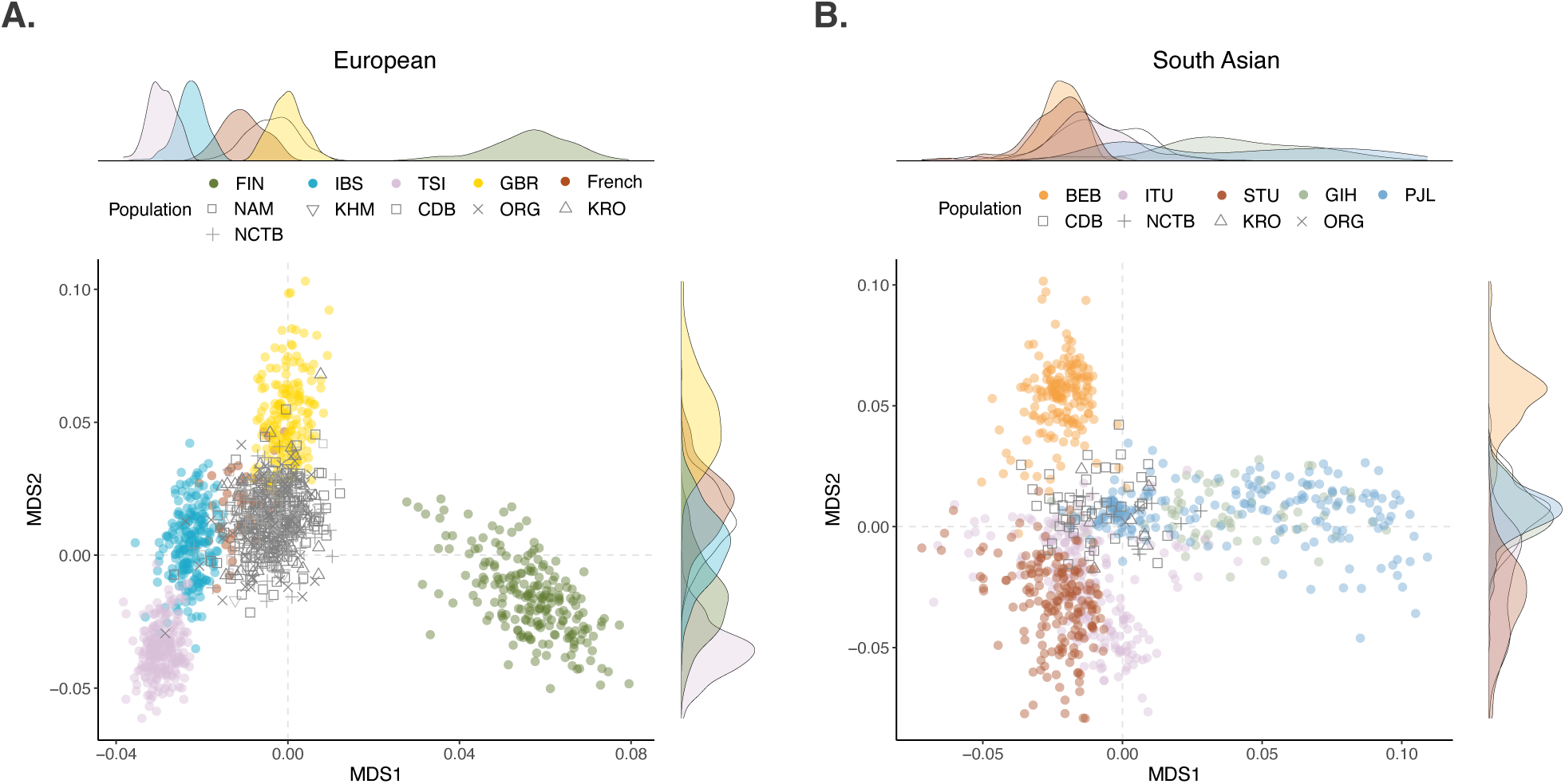
European and South Asian asMDS. Local ancestry results were filtered for haplotypes with 75% probability for the respective ancestry and individuals were filtered for at least 20% for the ancestry under investigation.

**Supplemental Figure 17).**
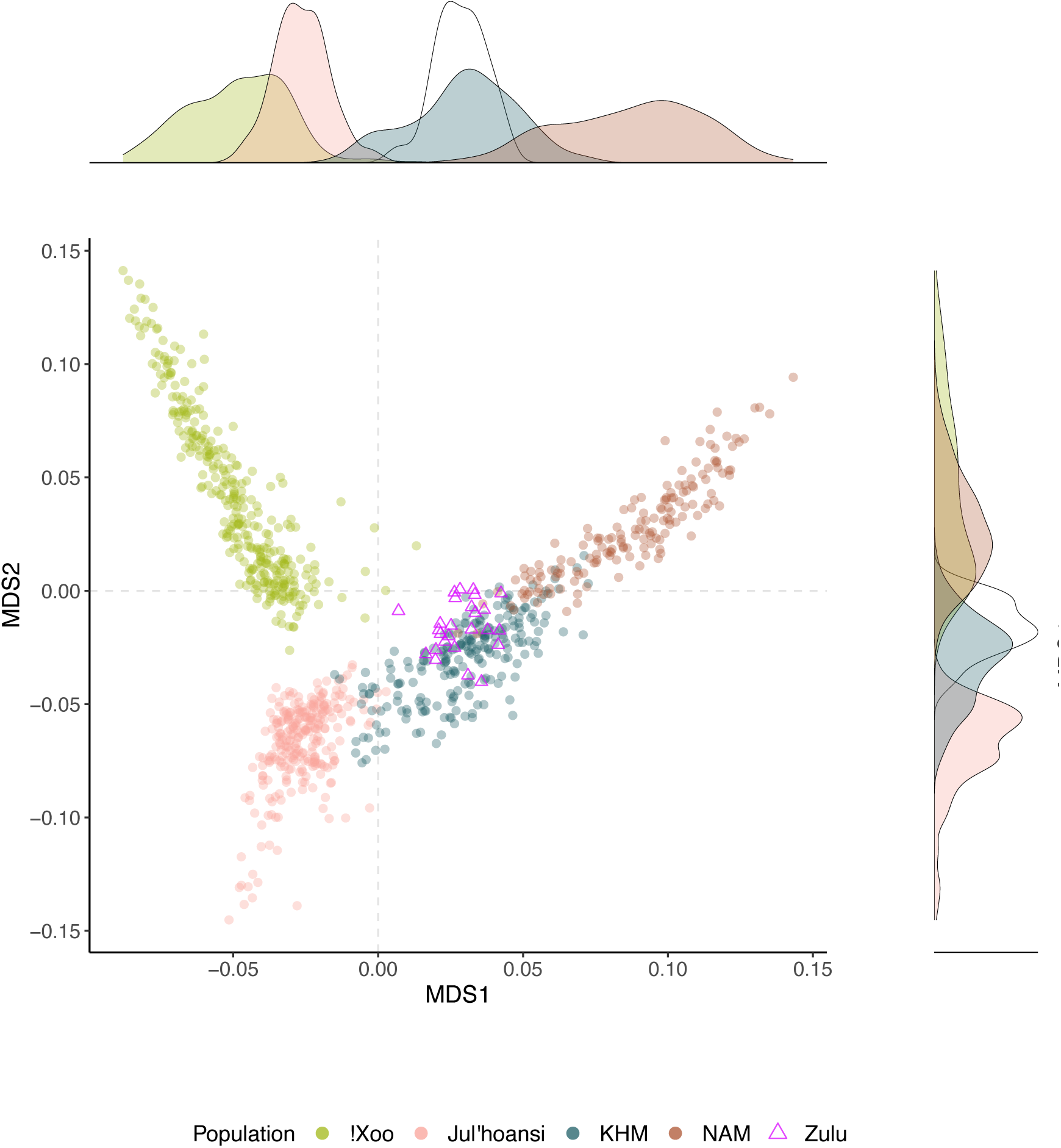
Zulu derived Khoe-San asMDS.

## Admixture Proportions

**Supplemental Table 1).**
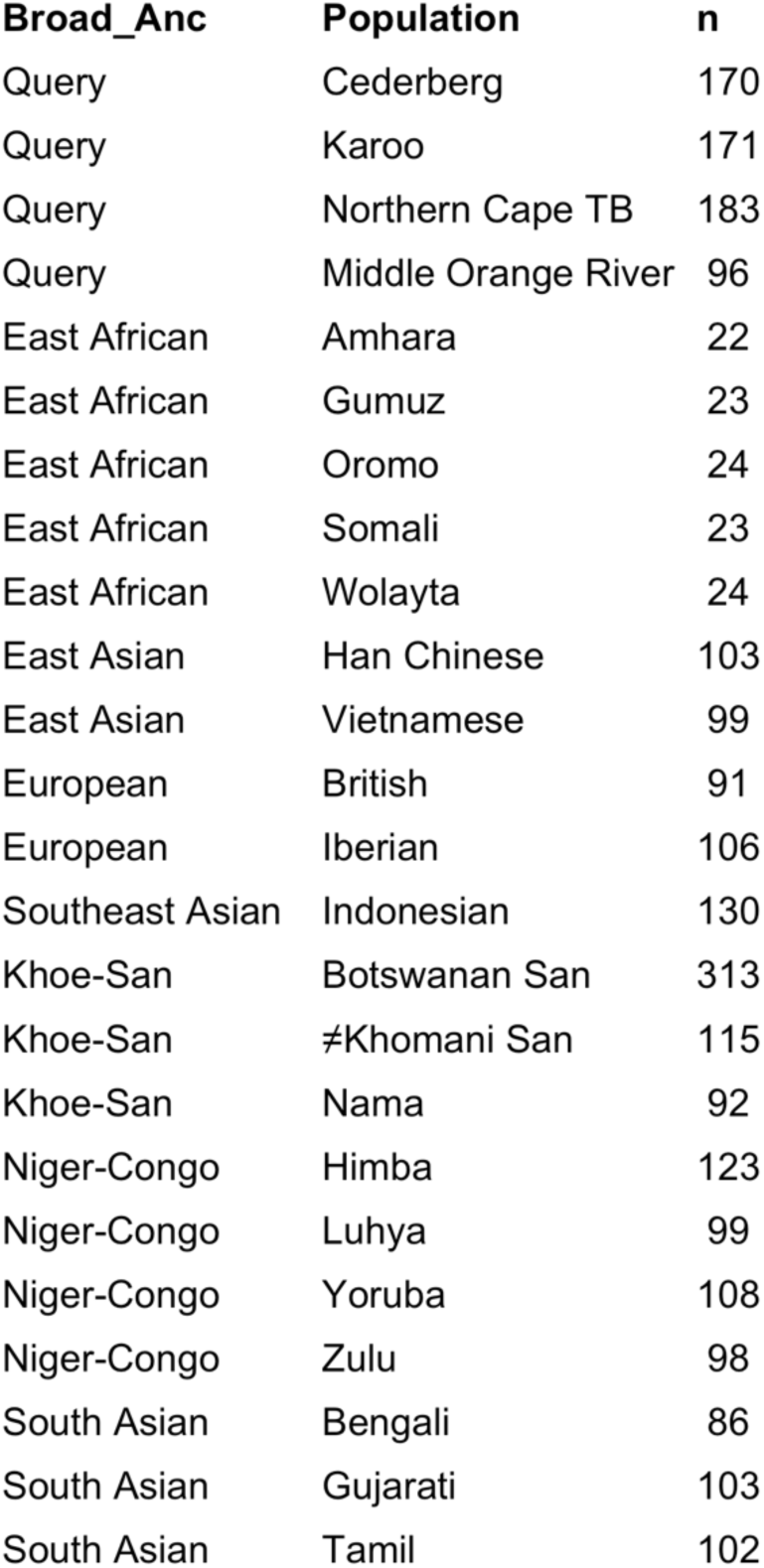
Sample Size for each population included in ADMIXTURE analysis, as well as their respective broader ancestry (Broad_Anc) category. Newly collected samples have the broader ancestry “Query”.

| Broad_Anc | Population | n |
| --- | --- | --- |
| Query | Cederberg | 170 |
| Query | Karoo | 171 |
| Query | Northern Cape TB | 183 |
| Query | Middle Orange River | 96 |
| East African | Amhara | 22 |
| East African | Gumuz | 23 |
| East African | Oromo | 24 |
| East African | Somali | 23 |
| East African | Wolayta | 24 |
| East Asian | Han Chinese | 103 |
| East Asian | Vietnamese | 99 |
| European | British | 91 |
| European | Iberian | 106 |
| Southeast Asian | Indonesian | 130 |
| Khoe-San | Botswanan San | 313 |
| Khoe-San | ≠Khomani San | 115 |
| Khoe-San | Nama | 92 |
| Niger-Congo | Himba | 123 |
| Niger-Congo | Luhya | 99 |
| Niger-Congo | Yoruba | 108 |
| Niger-Congo | Zulu | 98 |
| South Asian | Bengali | 86 |
| South Asian | Gujarati | 103 |
| South Asian | Tamil | 102 |

**Supplemental Table 2).**
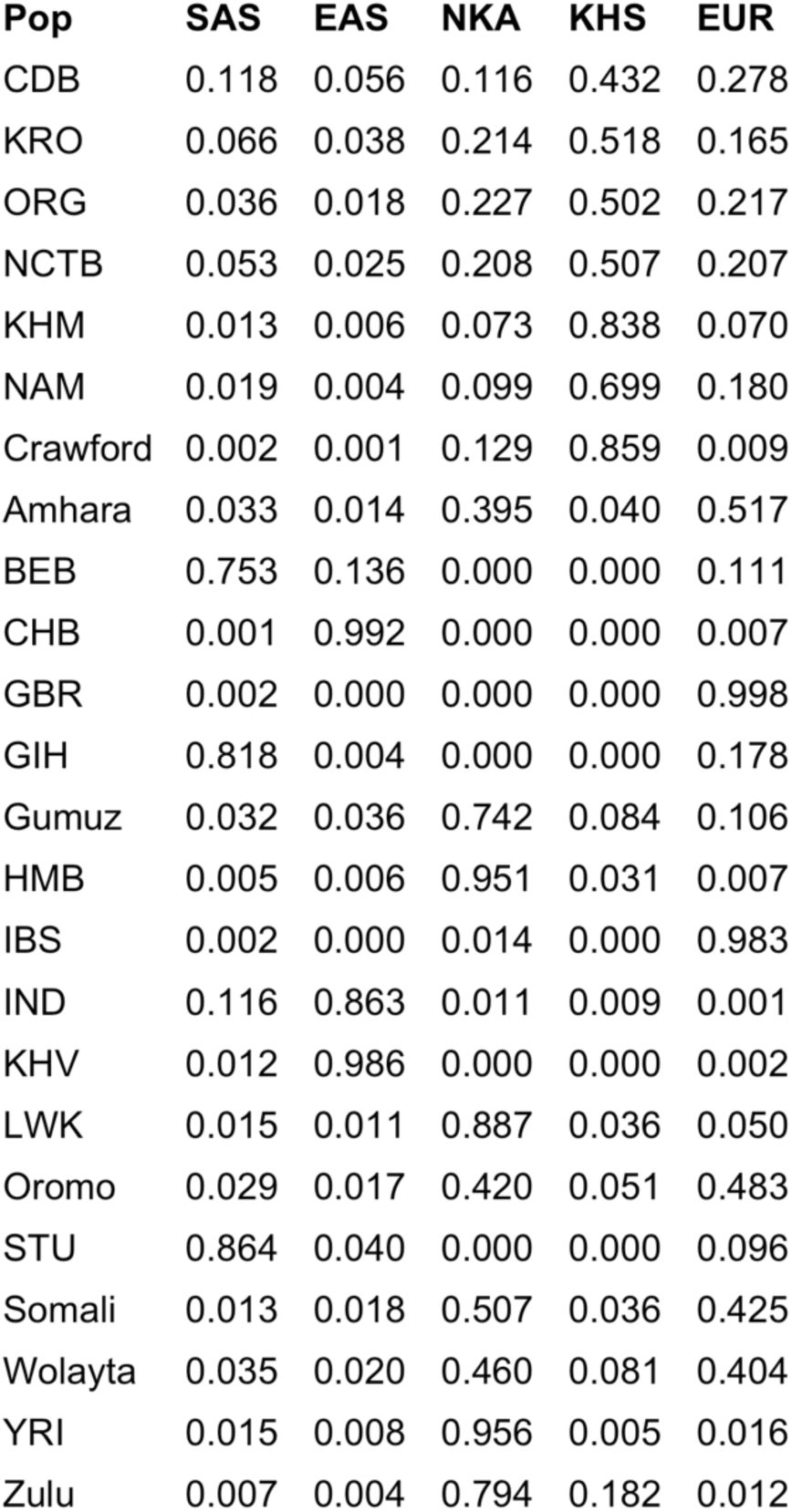
ADMIXTURE k=5.

**Supplemental Table 3).**
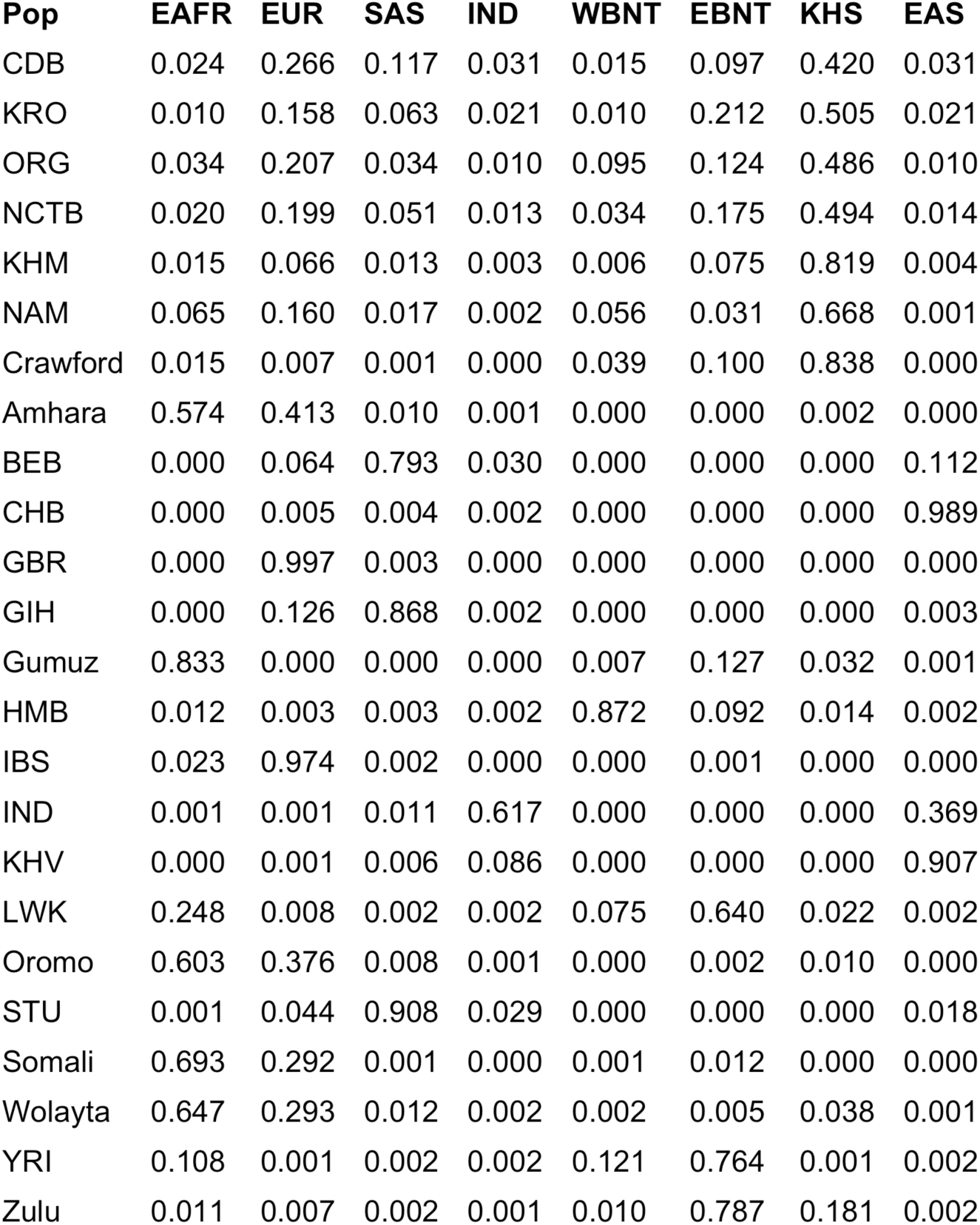
ADMIXTURE k=8.

**Supplemental Table 4).**
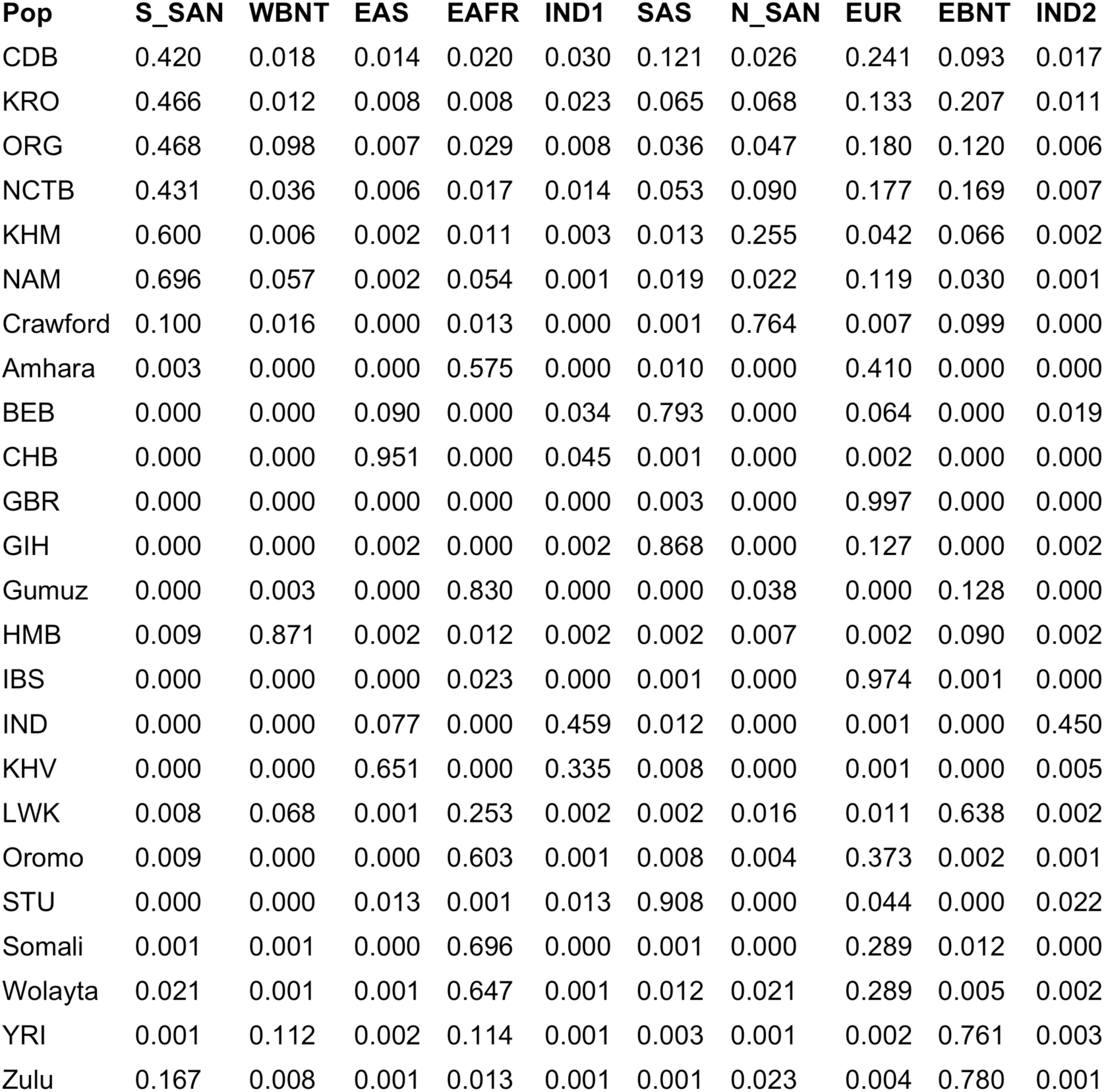
ADMIXTURE k=10.

## Population Centroids and Euclidean Distances

We aim to best characterize specific source populations for the five primary ancestries within the admixed study populations. We further investigated the relationships among populations by computing the Euclidean distances between each population’s centroid (i.e., a population’s mean coordinates along MDS1 and MDS2; see *Supplemental Methods*). For instance, the distances from the ≠Khomani San to the Cederberg (CDB), Karoo (KRO), and Orange River (ORG) populations are 0.036, 0.022, and 0.034, respectively, whereas the corresponding distances from NAM to these populations are 0.043, 0.059, and 0.046. These results suggest that, although Cederberg, Karoo, and Orange River are relatively similar in their overall MDS distribution, they share a more similar Khoe-San ancestry profile with ≠Khomani San than with the Nama. The Karoo (0.022) and NCTB (0.021) show the smallest distance to ≠Khomani San, indicating particularly close clustering.

Additionally, Zulu haplotypes fall with the ≠Khomani San (0.010), suggesting that the Khoe-San haplotypes present in Zulu individuals are genetically similar to ≠Khomani San. Among the query populations, we observe generally small distances to Yoruba (YRI), Luhya (LWK), and Zulu, compared to the Himba (HMB). Haplotypes from individuals sampled in the Karoo (0.002) demonstrate close similarities to Yoruba, while the Cederberg centroid appears slightly further from the Yoruba (0.009) and Luhya (0.017) (Supplemental Table 2). Participants sampled along Middle Orange River are more distant from the Yoruba (0.028) and Zulu (0.043), but still more similar to these groups than to Himba (0.072). Overall, these patterns indicate that the Cederberg, Karoo, and Orange River populations share closer ancestry profiles with Yoruba, Luhya, and Zulu than with Himba. Similarly, ≠Khomani San NKA haplotype centroid is closest to the Yoruba (0.004) and moderately more distant to the Zulu (0.021) and Luhya (0.025), with increasing distance to Himba (0.095). In contrast, the Nama NKA haplotype centroid is moderately distant to all NKA reference groups: Yoruba (0.036), Zulu (0.051), Luhya (0.040), Himba (0.064).

**Supplemental Figure 18).**
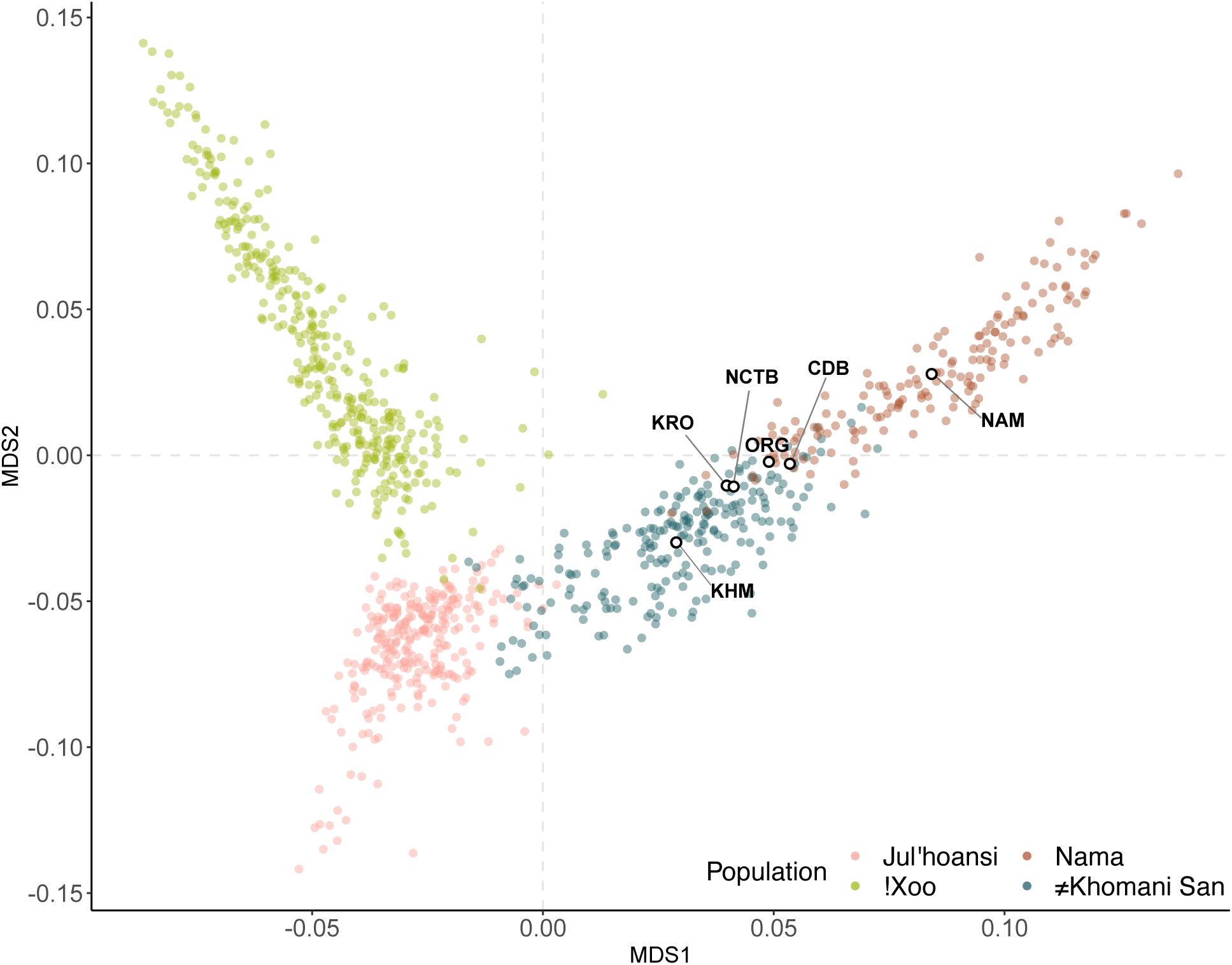
Khoe-San derived population centroids plotted against reference populations. Query sample haplotypes are not plotted for clearer visualization of reference populations distributions.

**Supplemental Figure 19).**
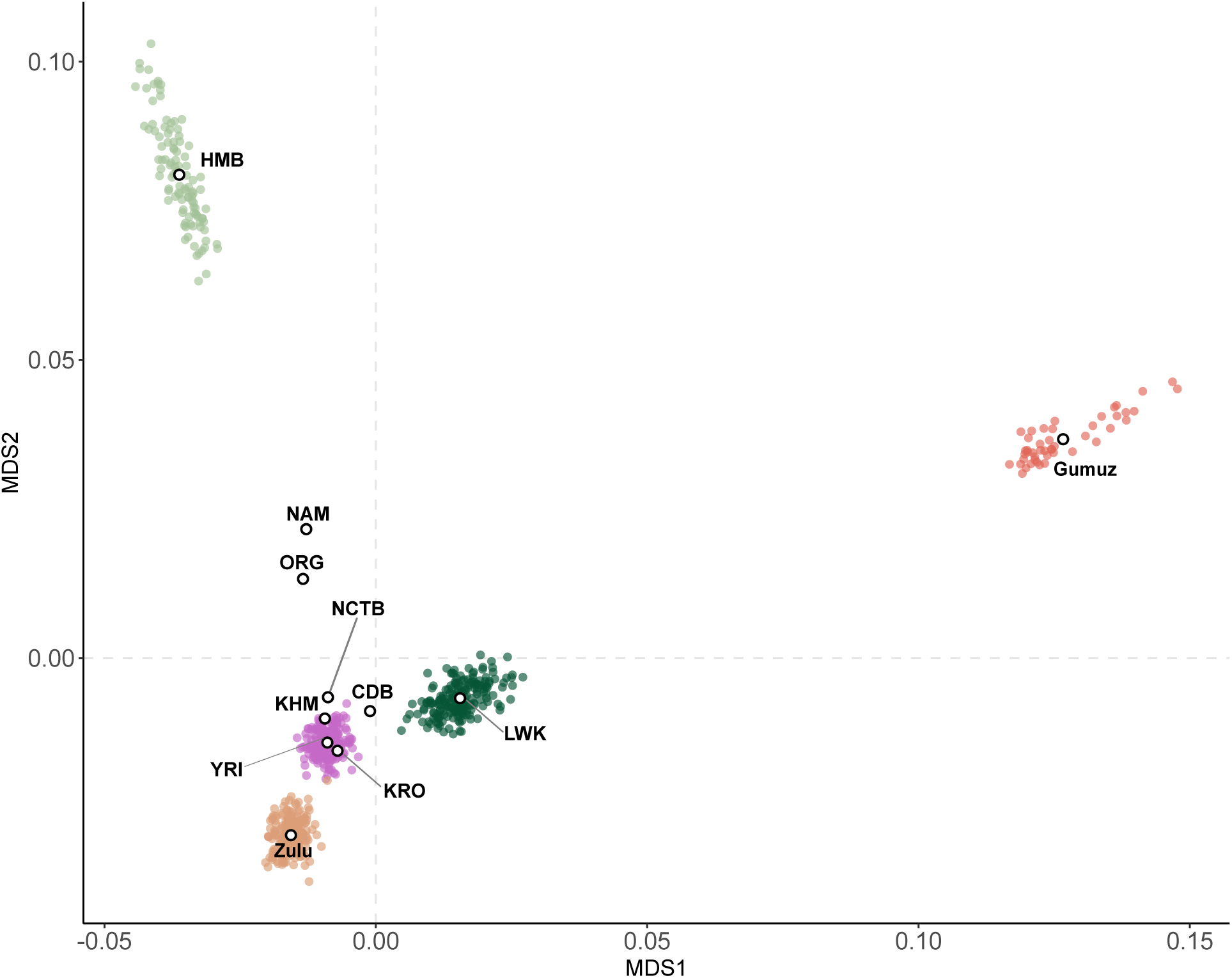
Non-Khoe-San derived population centroids plotted against non-Khoe-San reference populations. Query sample haplotypes are not plotted for clearer visualization of reference populations distributions.

**Supplemental Figure 20).**
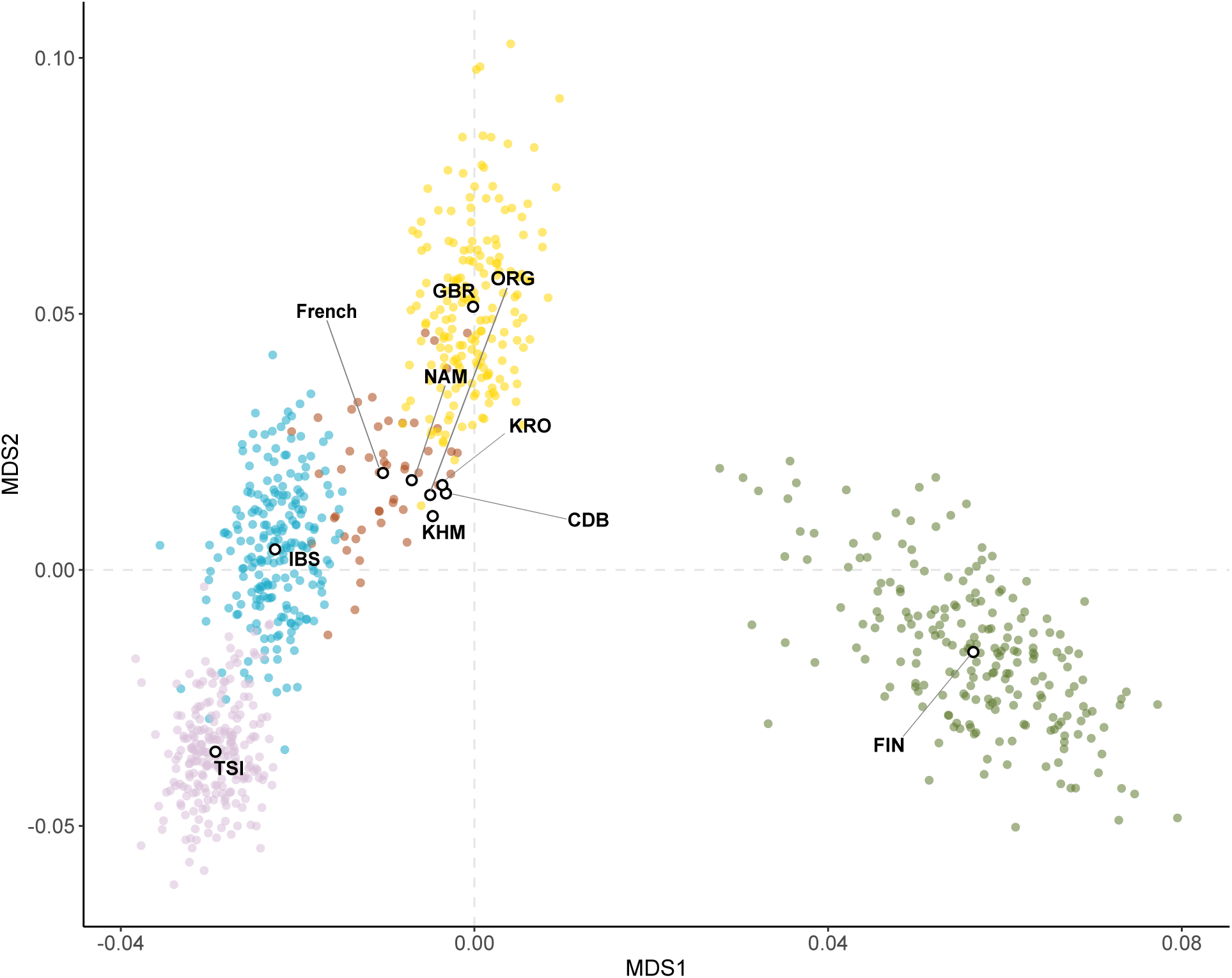
European derived centroids plotted against reference populations. Query sample haplotypes are not plotted for clearer visualization of reference populations distributions.

**Supplemental Figure 21).**
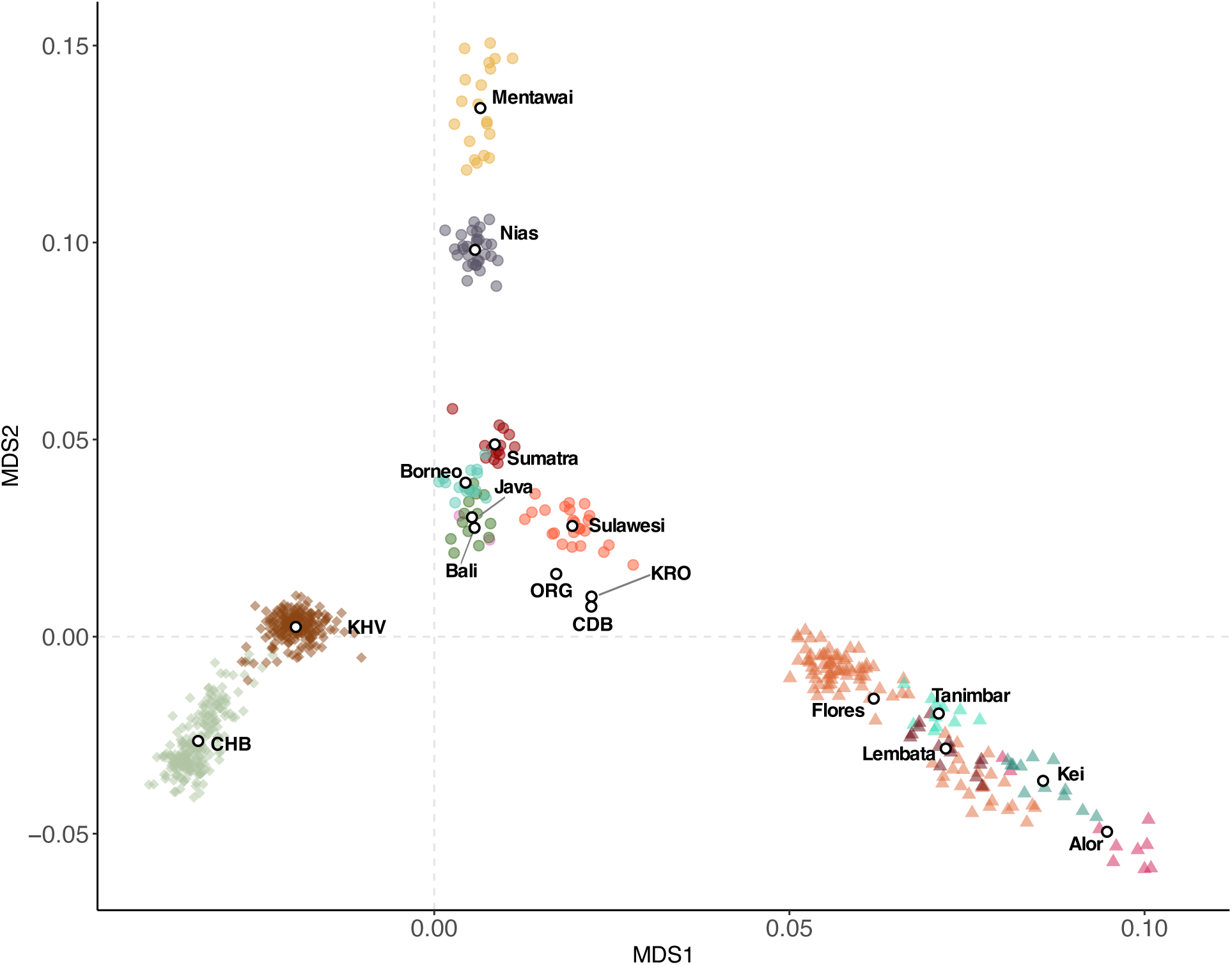
Other Asian derived centroids plotted against reference populations. Query sample haplotypes are not plotted for clearer visualization of reference populations distributions.

**Supplemental Figure 22).**
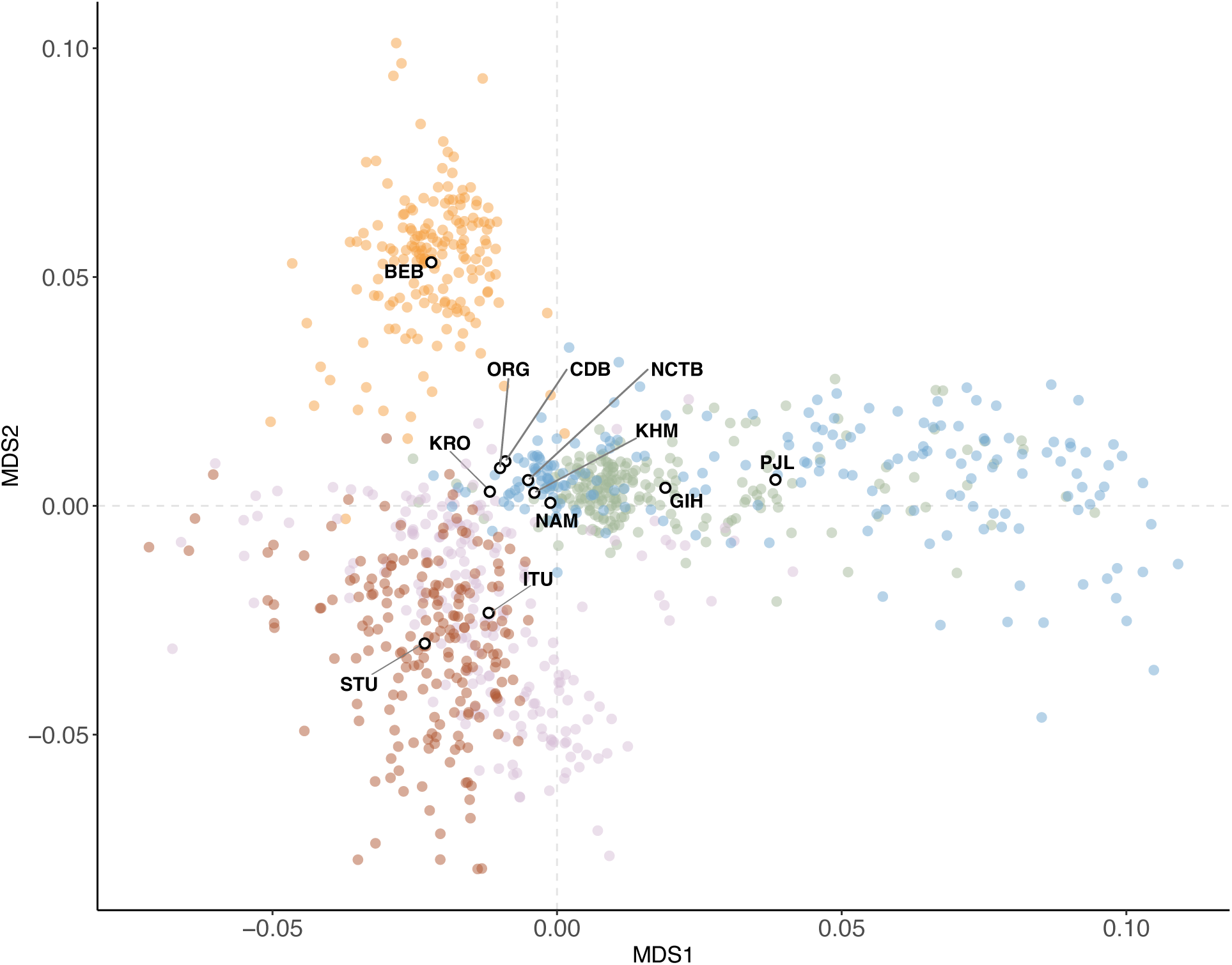
South Asian derived centroids plotted against reference populations. Query sample haplotypes are not plotted for clearer visualization of reference populations distributions.

**Supplemental Table 5).**
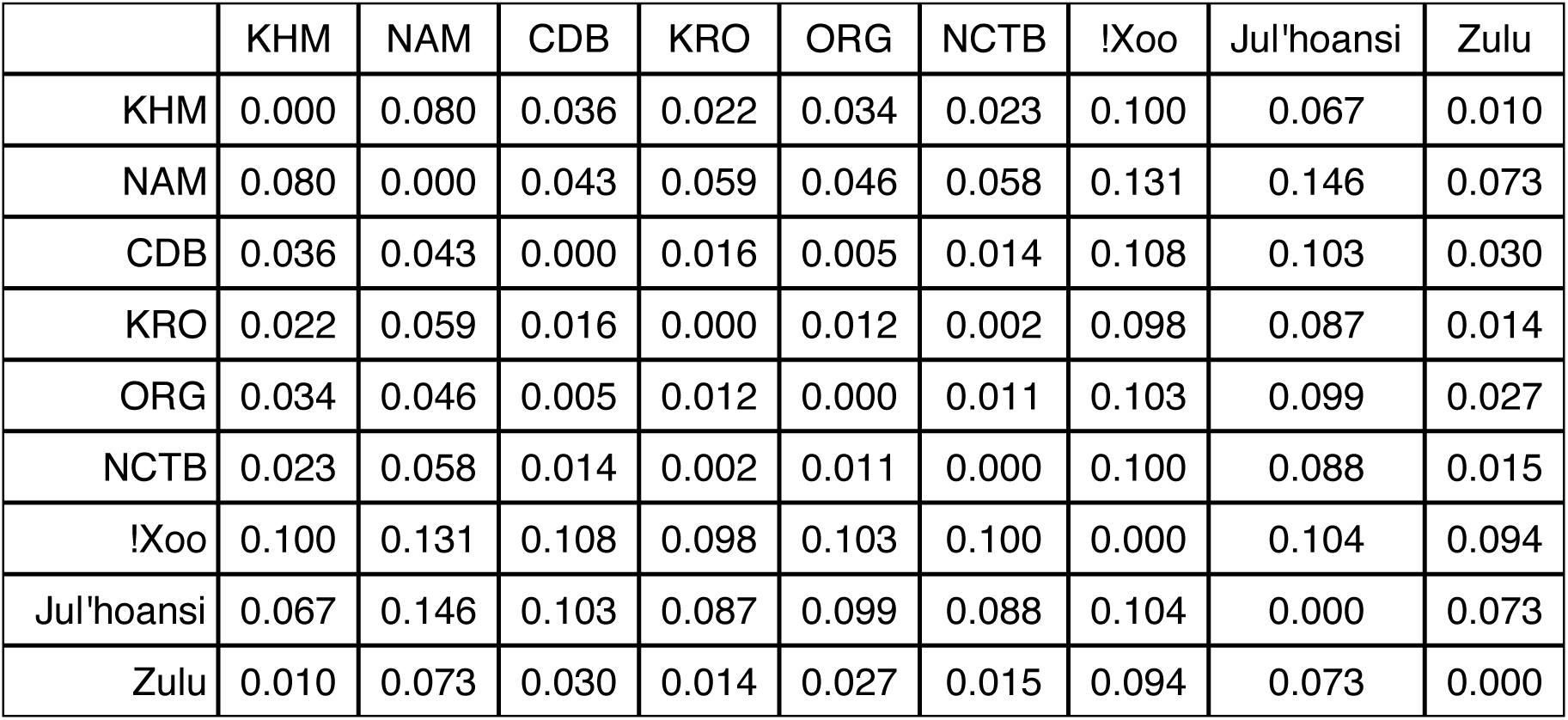
Khoe-San Euclidean distances between population centroids using Khoe-San ancestry-specific MDS1 and MDS2 coordinates.

**Supplemental Table 6).** Non-Khoe-San African Euclidean distances between population centroids using non-Khoe-San African ancestry-specific MDS1 and MDS2 coordinates.

|  | KHM | NAM | CDB | KRO | ORG | NCTB | Gumuz | HMB | LWK | YRI | Zulu |
| --- | --- | --- | --- | --- | --- | --- | --- | --- | --- | --- | --- |
| KHM | 0.000 | 0.032 | 0.008 | 0.006 | 0.024 | 0.004 | 0.144 | 0.095 | 0.025 | 0.004 | 0.021 |
| NAM | 0.032 | 0.000 | 0.033 | 0.038 | 0.008 | 0.028 | 0.140 | 0.064 | 0.040 | 0.036 | 0.051 |
| CDB | 0.008 | 0.033 | 0.000 | 0.009 | 0.025 | 0.008 | 0.136 | 0.097 | 0.017 | 0.009 | 0.025 |
| KRO | 0.006 | 0.038 | 0.009 | 0.000 | 0.030 | 0.009 | 0.144 | 0.101 | 0.024 | 0.002 | 0.017 |
| ORG | 0.024 | 0.008 | 0.025 | 0.030 | 0.000 | 0.020 | 0.142 | 0.072 | 0.035 | 0.028 | 0.043 |
| NCTB | 0.004 | 0.028 | 0.008 | 0.009 | 0.020 | 0.000 | 0.142 | 0.092 | 0.024 | 0.008 | 0.024 |
| Gumuz | 0.144 | 0.140 | 0.136 | 0.144 | 0.142 | 0.142 | 0.000 | 0.169 | 0.119 | 0.145 | 0.157 |
| HMB | 0.095 | 0.064 | 0.097 | 0.101 | 0.072 | 0.092 | 0.169 | 0.000 | 0.102 | 0.099 | 0.113 |
| LWK | 0.025 | 0.040 | 0.017 | 0.024 | 0.035 | 0.024 | 0.119 | 0.102 | 0.000 | 0.026 | 0.039 |
| YRI | 0.004 | 0.036 | 0.009 | 0.002 | 0.028 | 0.008 | 0.145 | 0.099 | 0.026 | 0.000 | 0.017 |
| Zulu | 0.021 | 0.051 | 0.025 | 0.017 | 0.043 | 0.024 | 0.157 | 0.113 | 0.039 | 0.017 | 0.000 |

**Supplemental Table 7).** European Euclidean distances between population centroids using European ancestry-specific MDS1 and MDS2 coordinates.

|  | KHM | NAM | CDB | KRO | ORG | NCTB | FIN | French | GBR | IBS | TSI |
| --- | --- | --- | --- | --- | --- | --- | --- | --- | --- | --- | --- |
| KHM | 0.000 | 0.005 | 0.003 | 0.004 | 0.002 | 0.003 | 0.066 | 0.011 | 0.042 | 0.020 | 0.052 |
| NAM | 0.005 | 0.000 | 0.003 | 0.003 | 0.003 | 0.003 | 0.070 | 0.006 | 0.037 | 0.020 | 0.055 |
| CDB | 0.003 | 0.003 | 0.000 | 0.001 | 0.002 | 0.000 | 0.067 | 0.009 | 0.038 | 0.021 | 0.055 |
| KRO | 0.004 | 0.003 | 0.001 | 0.000 | 0.002 | 0.001 | 0.067 | 0.008 | 0.037 | 0.022 | 0.056 |
| ORG | 0.002 | 0.003 | 0.002 | 0.002 | 0.000 | 0.001 | 0.067 | 0.009 | 0.040 | 0.020 | 0.054 |
| NCTB | 0.003 | 0.003 | 0.000 | 0.001 | 0.001 | 0.000 | 0.067 | 0.009 | 0.039 | 0.021 | 0.055 |
| FIN | 0.066 | 0.070 | 0.067 | 0.067 | 0.067 | 0.067 | 0.000 | 0.075 | 0.088 | 0.082 | 0.088 |
| French | 0.011 | 0.006 | 0.009 | 0.008 | 0.009 | 0.009 | 0.075 | 0.000 | 0.034 | 0.019 | 0.058 |
| GBR | 0.042 | 0.037 | 0.038 | 0.037 | 0.040 | 0.039 | 0.088 | 0.034 | 0.000 | 0.052 | 0.092 |
| IBS | 0.020 | 0.020 | 0.021 | 0.022 | 0.020 | 0.021 | 0.082 | 0.019 | 0.052 | 0.000 | 0.040 |
| TSI | 0.052 | 0.055 | 0.055 | 0.056 | 0.054 | 0.055 | 0.088 | 0.058 | 0.092 | 0.040 | 0.000 |

**Supplemental Table 8).** South Asian Euclidean distances between population centroids using South Asian ancestry-specific MDS1 and MDS2 coordinates.

|  | KHM | NAM | CDB | KRO | ORG | NCTB | BEB | GIH | ITU | PJL | STU |
| --- | --- | --- | --- | --- | --- | --- | --- | --- | --- | --- | --- |
| KHM | 0.000 | 0.004 | 0.009 | 0.008 | 0.008 | 0.003 | 0.054 | 0.023 | 0.027 | 0.042 | 0.038 |
| NAM | 0.004 | 0.000 | 0.012 | 0.011 | 0.012 | 0.006 | 0.057 | 0.020 | 0.026 | 0.040 | 0.038 |
| CDB | 0.009 | 0.012 | 0.000 | 0.007 | 0.002 | 0.006 | 0.045 | 0.029 | 0.033 | 0.048 | 0.042 |
| KRO | 0.008 | 0.011 | 0.007 | 0.000 | 0.005 | 0.007 | 0.051 | 0.031 | 0.026 | 0.050 | 0.035 |
| ORG | 0.008 | 0.012 | 0.002 | 0.005 | 0.000 | 0.006 | 0.047 | 0.029 | 0.032 | 0.048 | 0.041 |
| NCTB | 0.003 | 0.006 | 0.006 | 0.007 | 0.006 | 0.000 | 0.051 | 0.024 | 0.030 | 0.043 | 0.040 |
| BEB | 0.054 | 0.057 | 0.045 | 0.051 | 0.047 | 0.051 | 0.000 | 0.064 | 0.077 | 0.077 | 0.083 |
| GIH | 0.023 | 0.020 | 0.029 | 0.031 | 0.029 | 0.024 | 0.064 | 0.000 | 0.041 | 0.019 | 0.054 |
| ITU | 0.027 | 0.026 | 0.033 | 0.026 | 0.032 | 0.030 | 0.077 | 0.041 | 0.000 | 0.058 | 0.013 |
| PJL | 0.042 | 0.040 | 0.048 | 0.050 | 0.048 | 0.043 | 0.077 | 0.019 | 0.058 | 0.000 | 0.071 |
| STU | 0.038 | 0.038 | 0.042 | 0.035 | 0.041 | 0.040 | 0.083 | 0.054 | 0.013 | 0.071 | 0.000 |

**Supplemental Table 9).** Southeast and East Asian distances. Euclidean distances between population centroids using Southeast and East Asian ancestry-specific MDS1 and MDS2 coordinates.

|  | CDB | KRO | ORG | NCTB | Sulawesi | Java | Flores | Sumatra | Alor | Bali | Borneo | CHB | KHV | Kei | Lembata | Mentawai | Nias | Tanimbar |
| --- | --- | --- | --- | --- | --- | --- | --- | --- | --- | --- | --- | --- | --- | --- | --- | --- | --- | --- |
| CDB | 0.000 | 0.003 | 0.010 | 0.010 | 0.021 | 0.028 | 0.046 | 0.043 | 0.092 | 0.026 | 0.036 | 0.065 | 0.042 | 0.077 | 0.062 | 0.127 | 0.092 | 0.056 |
| KRO | 0.003 | 0.000 | 0.008 | 0.008 | 0.018 | 0.026 | 0.047 | 0.041 | 0.094 | 0.024 | 0.034 | 0.066 | 0.042 | 0.079 | 0.063 | 0.125 | 0.089 | 0.057 |
| ORG | 0.010 | 0.008 | 0.000 | 0.003 | 0.012 | 0.019 | 0.055 | 0.034 | 0.101 | 0.016 | 0.026 | 0.066 | 0.039 | 0.086 | 0.070 | 0.119 | 0.083 | 0.064 |
| NCTB | 0.010 | 0.008 | 0.003 | 0.000 | 0.011 | 0.019 | 0.054 | 0.033 | 0.101 | 0.017 | 0.026 | 0.068 | 0.041 | 0.086 | 0.070 | 0.117 | 0.082 | 0.064 |
| Sulawesi | 0.021 | 0.018 | 0.012 | 0.011 | 0.000 | 0.014 | 0.061 | 0.023 | 0.108 | 0.014 | 0.019 | 0.076 | 0.047 | 0.093 | 0.077 | 0.107 | 0.071 | 0.070 |
| Java | 0.028 | 0.026 | 0.019 | 0.019 | 0.014 | 0.000 | 0.073 | 0.019 | 0.120 | 0.003 | 0.009 | 0.069 | 0.037 | 0.105 | 0.089 | 0.104 | 0.068 | 0.082 |
| Flores | 0.046 | 0.047 | 0.055 | 0.054 | 0.061 | 0.073 | 0.000 | 0.084 | 0.047 | 0.071 | 0.079 | 0.096 | 0.083 | 0.032 | 0.016 | 0.160 | 0.127 | 0.010 |
| Sumatra | 0.043 | 0.041 | 0.034 | 0.033 | 0.023 | 0.019 | 0.084 | 0.000 | 0.131 | 0.021 | 0.011 | 0.086 | 0.054 | 0.115 | 0.100 | 0.085 | 0.049 | 0.093 |
| Alor | 0.092 | 0.094 | 0.101 | 0.101 | 0.108 | 0.120 | 0.047 | 0.131 | 0.000 | 0.118 | 0.126 | 0.130 | 0.125 | 0.016 | 0.031 | 0.204 | 0.172 | 0.038 |
| Bali | 0.026 | 0.024 | 0.016 | 0.017 | 0.014 | 0.003 | 0.071 | 0.021 | 0.118 | 0.000 | 0.012 | 0.067 | 0.036 | 0.103 | 0.087 | 0.107 | 0.071 | 0.081 |
| Borneo | 0.036 | 0.034 | 0.026 | 0.026 | 0.019 | 0.009 | 0.079 | 0.011 | 0.126 | 0.012 | 0.000 | 0.076 | 0.044 | 0.111 | 0.096 | 0.095 | 0.059 | 0.089 |
| CHB | 0.065 | 0.066 | 0.066 | 0.068 | 0.076 | 0.069 | 0.096 | 0.086 | 0.130 | 0.067 | 0.076 | 0.000 | 0.032 | 0.119 | 0.105 | 0.165 | 0.131 | 0.104 |
| KHV | 0.042 | 0.042 | 0.039 | 0.041 | 0.047 | 0.037 | 0.083 | 0.054 | 0.125 | 0.036 | 0.044 | 0.032 | 0.000 | 0.112 | 0.097 | 0.134 | 0.099 | 0.093 |
| Kei | 0.077 | 0.079 | 0.086 | 0.086 | 0.093 | 0.105 | 0.032 | 0.115 | 0.016 | 0.103 | 0.111 | 0.119 | 0.112 | 0.000 | 0.016 | 0.188 | 0.157 | 0.023 |
| Lembata | 0.062 | 0.063 | 0.070 | 0.070 | 0.077 | 0.089 | 0.016 | 0.100 | 0.031 | 0.087 | 0.096 | 0.105 | 0.097 | 0.016 | 0.000 | 0.175 | 0.143 | 0.009 |
| Mentawai | 0.127 | 0.125 | 0.119 | 0.117 | 0.107 | 0.104 | 0.160 | 0.085 | 0.204 | 0.107 | 0.095 | 0.165 | 0.134 | 0.188 | 0.175 | 0.000 | 0.036 | 0.167 |
| Nias | 0.092 | 0.089 | 0.083 | 0.082 | 0.071 | 0.068 | 0.127 | 0.049 | 0.172 | 0.071 | 0.059 | 0.131 | 0.099 | 0.157 | 0.143 | 0.036 | 0.000 | 0.135 |
| Tanimbar | 0.056 | 0.057 | 0.064 | 0.064 | 0.070 | 0.082 | 0.010 | 0.093 | 0.038 | 0.081 | 0.089 | 0.104 | 0.093 | 0.023 | 0.009 | 0.167 | 0.135 | 0.000 |

## SPRUCE

**Supplemental Figure 23).**
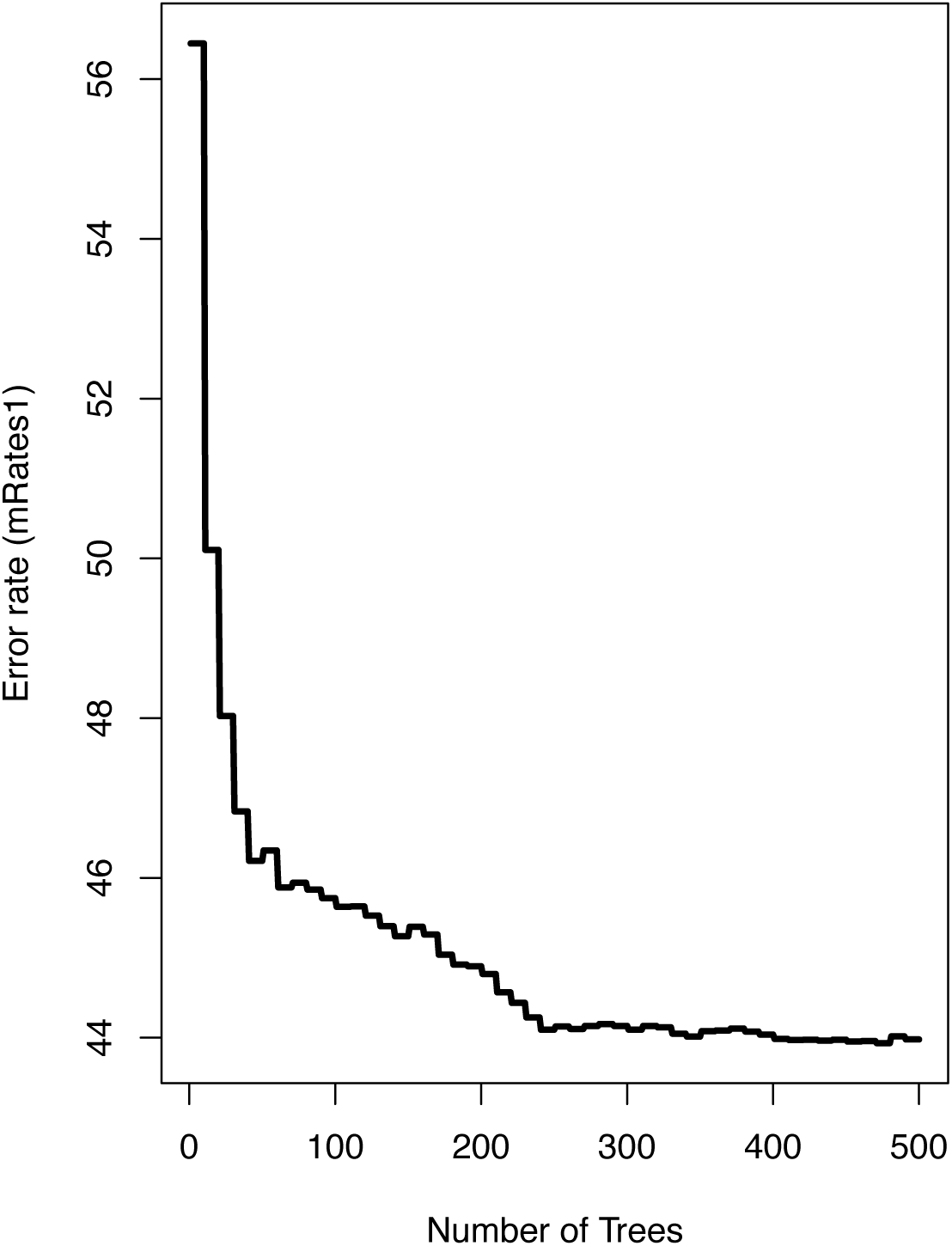
Error rate plot showing the error rate decreasing as the number of trees in the random forest increases for IBD segments 2-6 cM representing 56-13 generations ago.

**Supplemental Figure 24).**
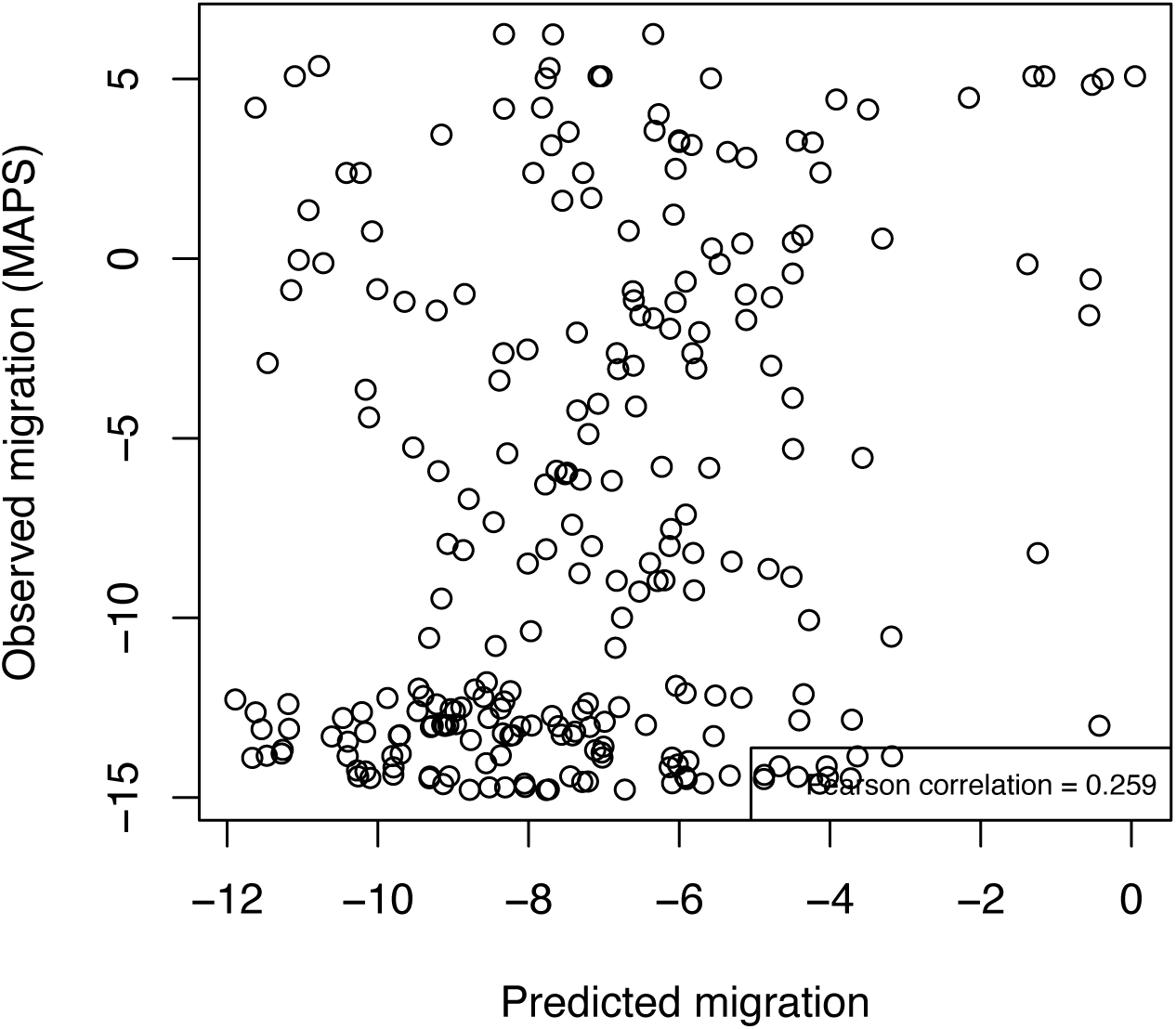
Scatterplot showing predicted migration plotted against observed migration (Pearson correlations = 0.259) for IBD segments 2-6 cM/56-13 generations ago.

**Supplemental Figure 25).**
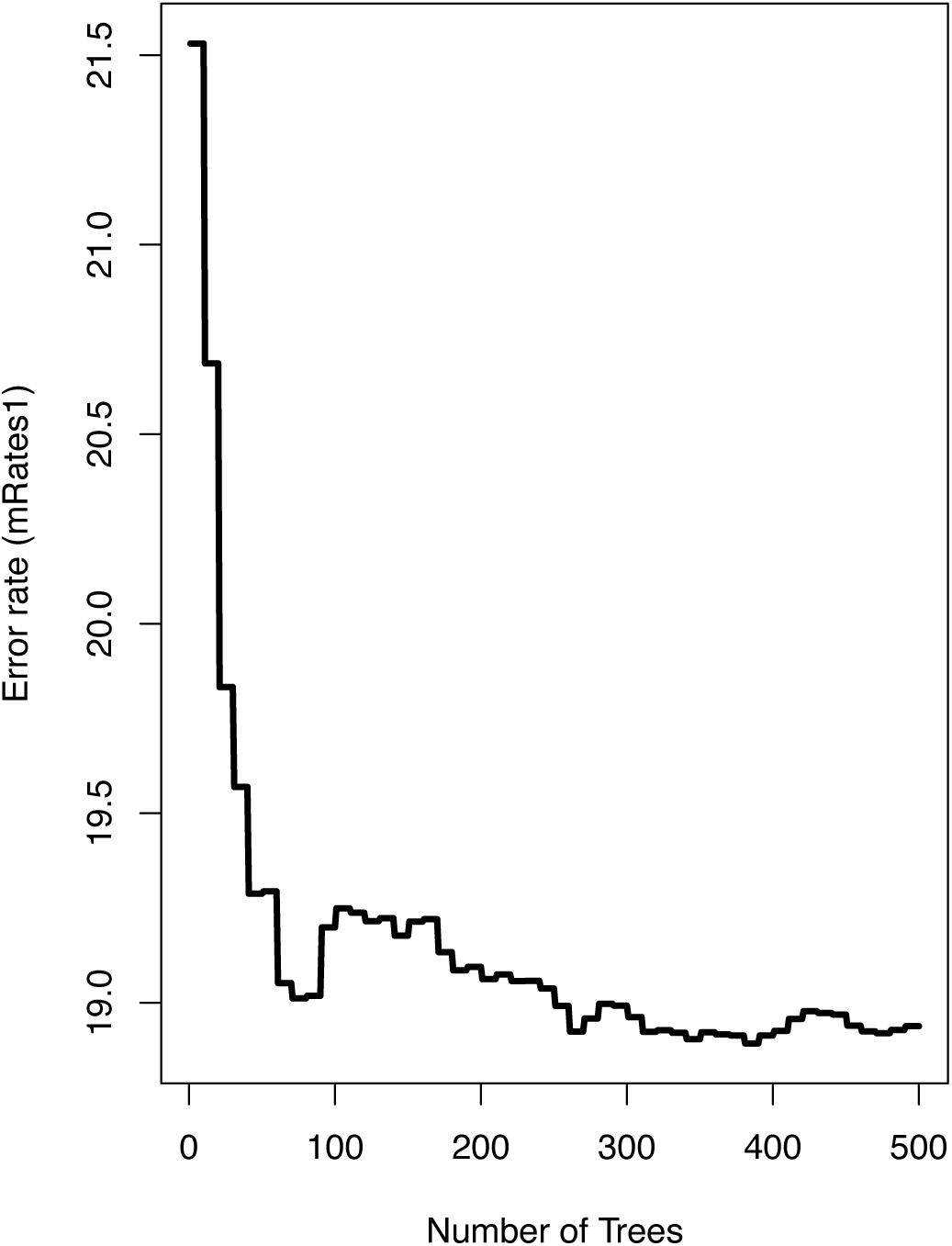
Error rate plot showing the error rate decreasing as the number of trees in the random forest increases for IBD segments greater than 6 cM representing less than 13 generations ago.

**Supplemental Figure 26).**
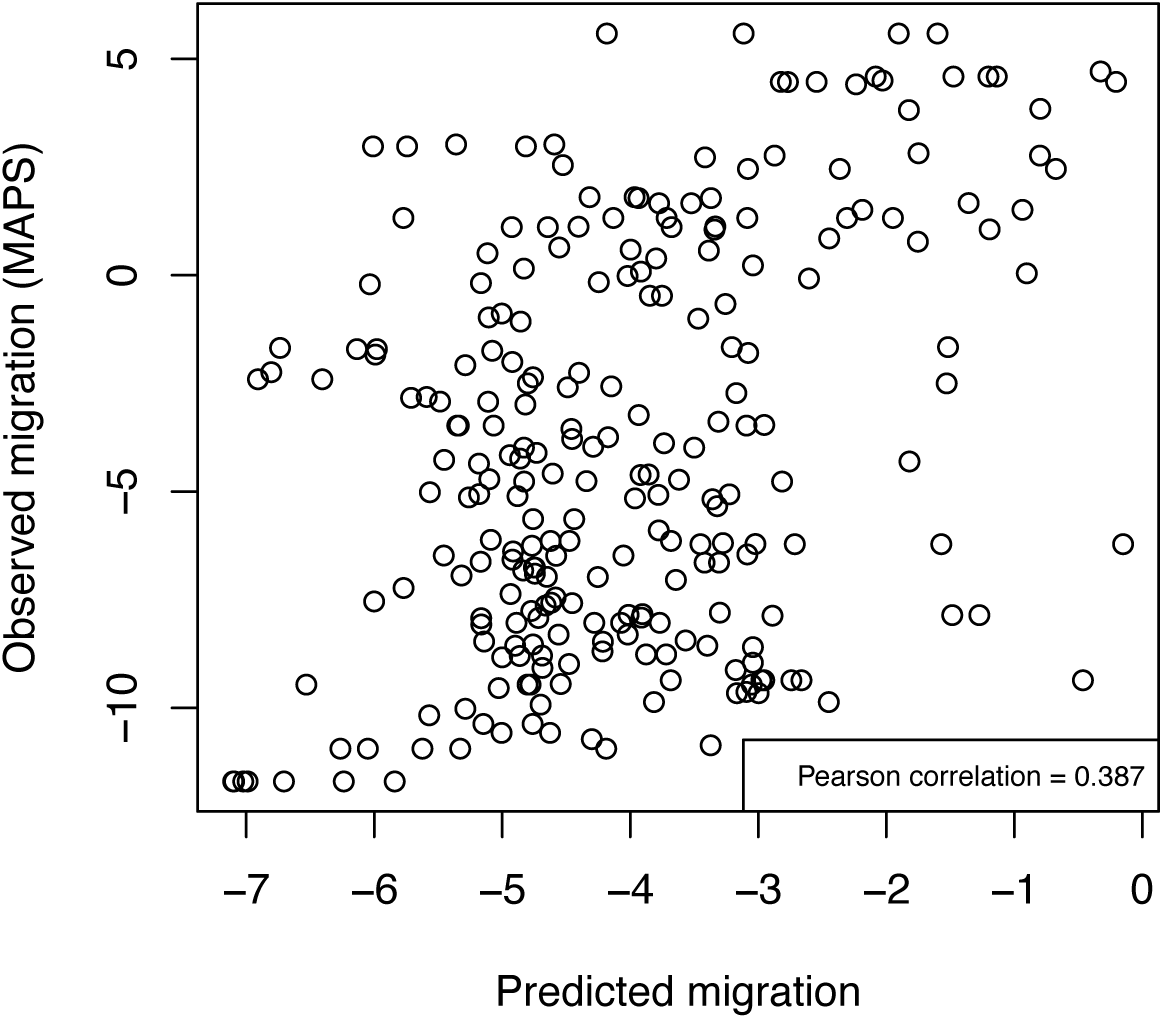
Scatterplot showing predicted migration plotted against observed migration (Pearson correlations = 0.387) for IBD segments greater than 6 cM/ less than13 generations ago.

**Supplemental Figure 27).**
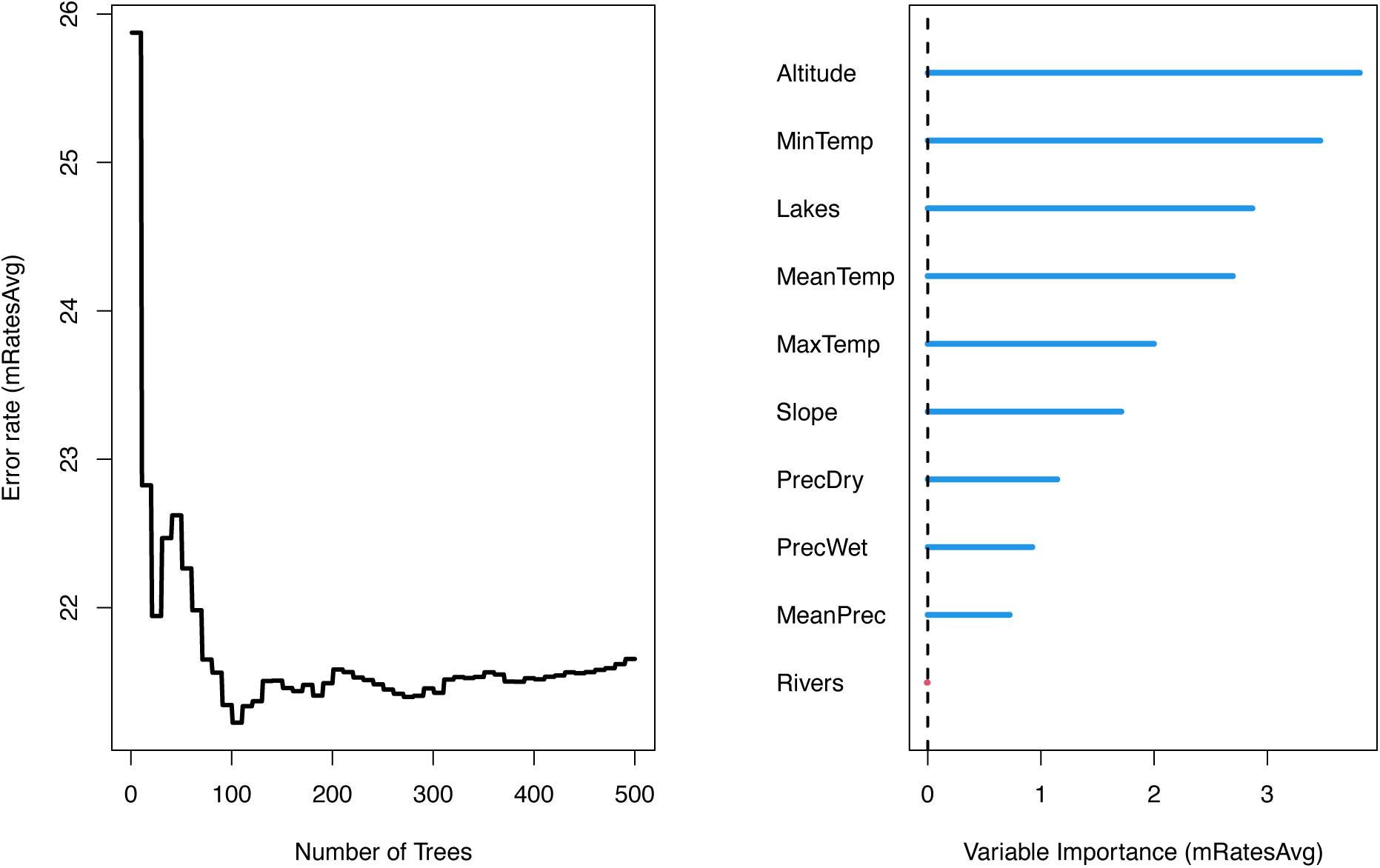
SPRUCE Random forest importance table average across time periods (2-6cM and greater than 6cM), 56> generations ago showing altitude and minimum temperature during the coldest month being the most important variables explaining the variation in estimated migration rate.

**Supplemental Figure 28).**
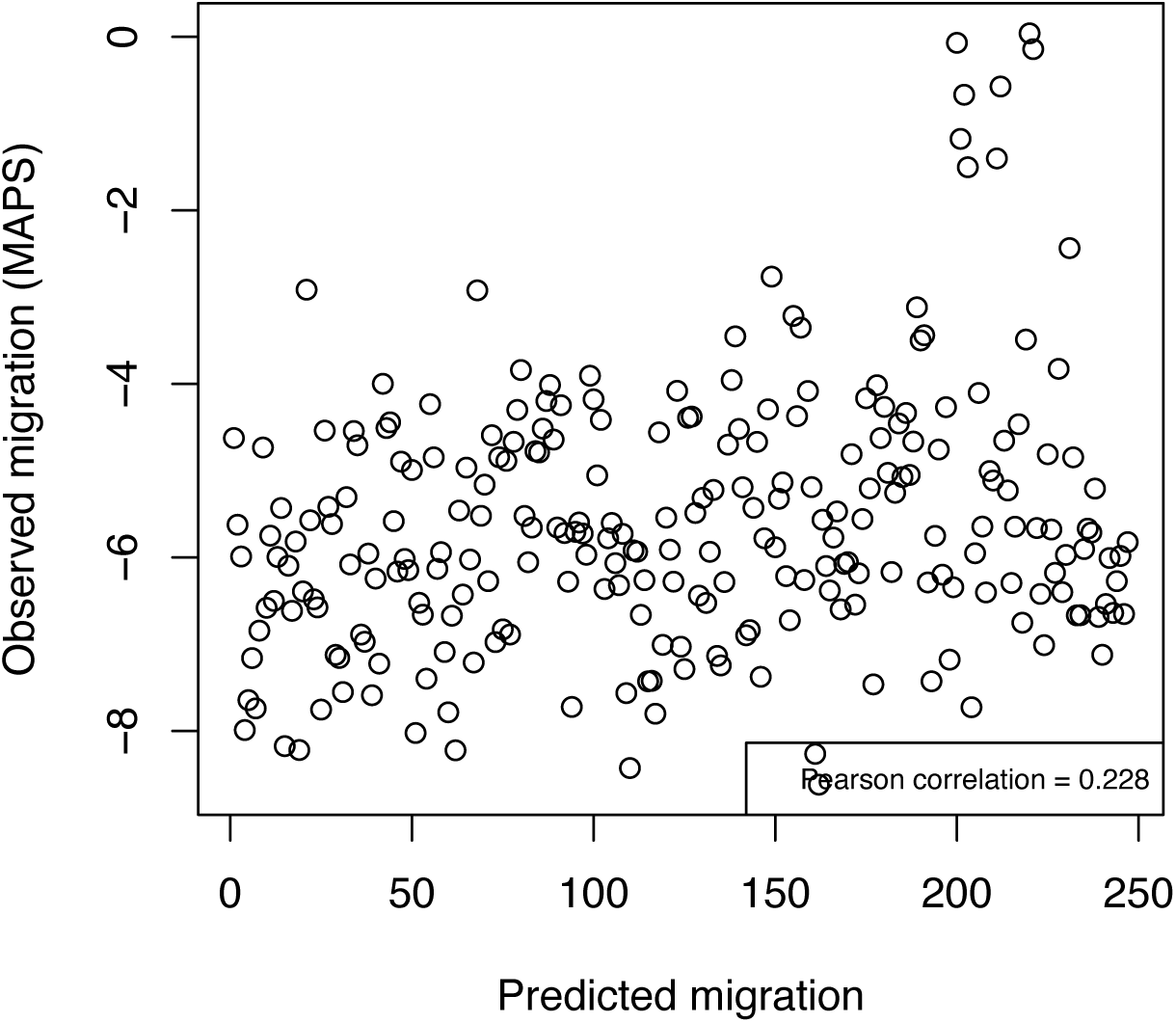
Scatterplot showing predicted migration plotted against observed migration (Pearson correlations = 0.228) on average.

**Supplemental Figure 29).**
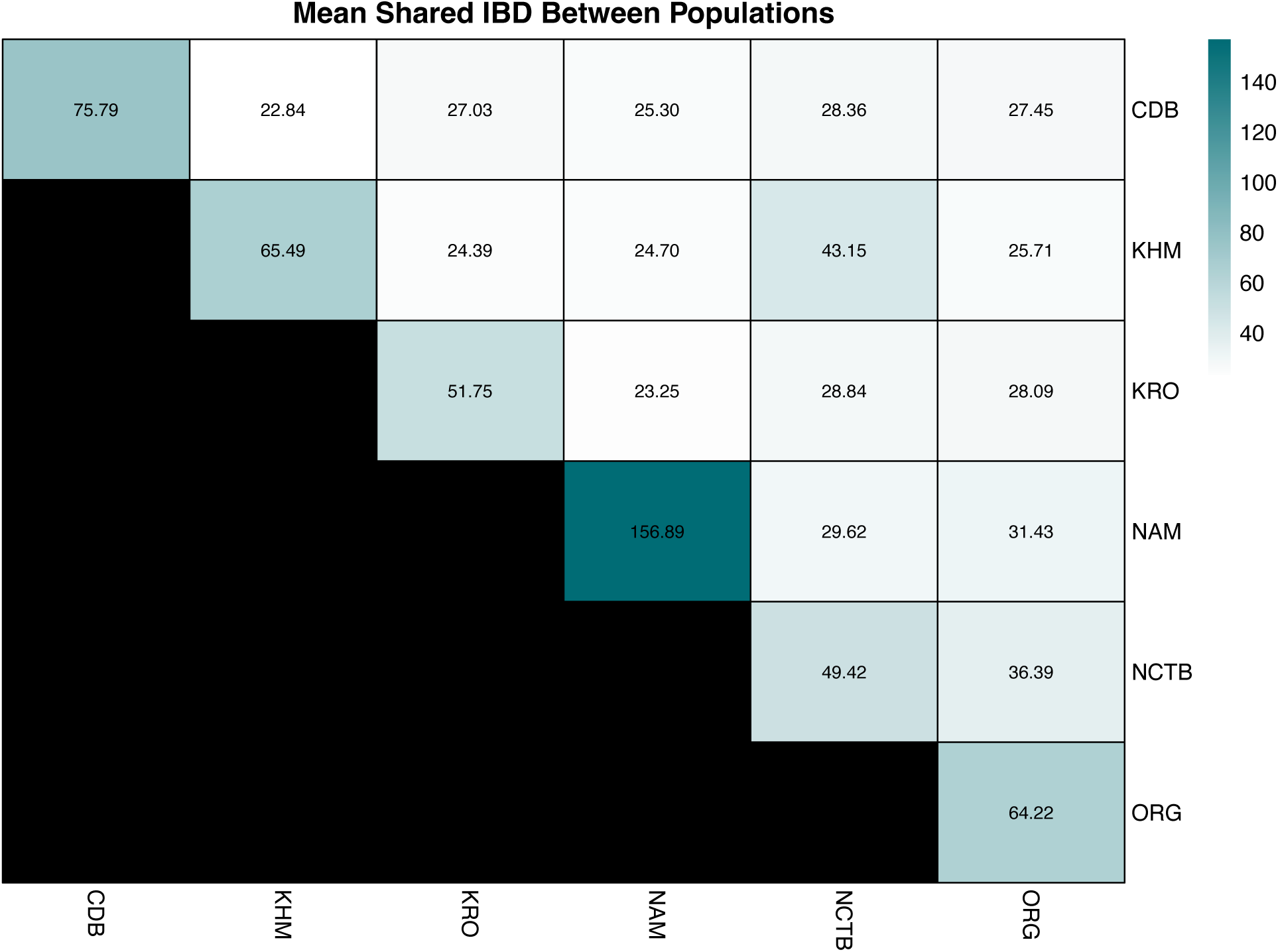
Mean shared IBD length between and within Khoe-San and Khoe-San descendant communities.

